# In plants distal regulatory sequences overlap with unmethylated rather than low-methylated regions, in contrast to mammals

**DOI:** 10.1101/2020.03.24.005678

**Authors:** Rurika Oka, Mattijs Bliek, Huub C.J. Hoefsloot, Maike Stam

## Abstract

**Background:** DNA methylation is an important factor in the regulation of gene expression and genome stability. High DNA methylation levels are associated with transcriptional repression. In mammalian systems, unmethylated, low methylated and fully methylated regions (UMRs, LMRs, and FMRs, respectively) can be distinguished. UMRs are associated with proximal regulatory regions, while LMRs are associated with distal regulatory regions. Although DNA methylation is mainly limited to the CG context in mammals, while it occurs in CG, CHG and CHH contexts in plants, UMRs and LMRs were expected to occupy similar genomic sequences in both mammals and plants.

**Results:** This study investigated major model and crop plants such as *Arabidopsis thaliana*, tomato (*Solanum lycopersicum*), rice (*Oryza sativa*) and maize (*Zea mays*), and shows that plant genomes can also be subdivided in UMRs, LMRs and FMRs, but that LMRs are mainly present in the CHG context rather than the CG context. Strikingly, the identified CHG LMRs were enriched in transposable elements rather than regulatory regions. Maize candidate regulatory regions overlapped with UMRs. LMRs were enriched for heterochromatic histone modifications and depleted for DNase accessibility and H3K9 acetylation. CHG LMRs form a distinct, abundant cluster of loci, indicating they have a different role than FMRs.

**Conclusions:** Both mammalian and plant genomes can be segmented in three distinct classes of loci, UMRs, LMRs and FMRs, indicating similar underlying mechanisms. Unlike in mammals, distal regulatory sequences in plants appear to overlap with UMRs instead of LMRs. Our data indicate that LMRs in plants have a different function than those in mammals.

## Background

Epigenetic marks, including DNA methylation and histone modifications, are important factors involved in regulating chromatin structure and gene expression. In both animals and plants, high levels of DNA methylation lead to less accessible chromatin and have been associated with transcriptional repression, while low DNA methylation has been observed at active distal and proximal gene regulatory regions and gene bodies [1–6].

Overall, functions of DNA methylation, such as silencing of transposable elements, genome stability and transcription regulation, are similar in plants and mammals [1,2,6–8]. There are also functions that are identified for mammals, but not yet reported for plants, such as a widespread tissue-specific demethylation of regulatory elements and inhibition of cryptic initiation of transcription [6,9–11]. One well-known difference between animals and plants is that while animals have primarily methylation in CG context in most tissues, plants also have abundant methylation in CHG, and low methylation levels in CHH context (where H is A, T or C) [1,12]. Although the DNA methylation pathways are quite conserved through the plant kingdom they also display differences [11–13]. As the vast majority of the mechanistic insight into DNA methylation is obtained in Arabidopsis, below the pathways are discussed for Arabidopsis.

In the RNA-directed DNA methylation (RdDM) pathway, small interfering RNAs (siRNAs) guide the *de novo* DNA methyltransferase DOMAINS REARRANGED METHYLTRANSFERASE2 (DRM2) to complementary DNA to initiate methylation [1,8]). For maintaining DNA methylation in the different sequence contexts, various pathways exist. Symmetrical methylation, methylation in CG and CHG contexts, is maintained through maintenance methylation pathways. CG methylation is maintained by a maintenance CG methyltransferase, METHYLTRANSFERASE 1 (MET 1) (DNA methyltransferase 1 (DNMT1) in animals). CHG methylation is maintained by CHROMOMETHYLASE 3 (CMT3), which binds to histone 3 lysine 9 dimethylation (H3K9me2) and methylates closeby CHG sites; a feedback loop is completed by the dimethylation of H3K9 by SUPPRESSOR OF VARIEGATION 3-9 HOMOLOGUE 4 (SUVH4, also called KRYPTONITE (KYP)), SUVH5 and SUVH6, which all recognize cytosine methylation. Non-symmetrical methylation (CHH) is maintained through two different mechanisms: the RdDM pathway and a CHROMOMETHYLASE 2 (CMT2)-mediated methylation pathway. SiRNAs guide the *de novo* DNA methyltransferase DRM2 to all cytosine sites in euchromatin, including CHH sites, while CHH sites in heterochromatin are methylated by CMT2, which recognises and binds to H3K9me2. The latter pathway is widely present in eudicots and monocots, though not in maize [14,15].

Another difference between DNA methylation in animals and plants appears the stability of DNA methylation in euchromatin. In animals, primarily at regulatory sequences, levels of DNA methylation are dynamically regulated between tissues and developmental stages [6,16–18]. Although there is evidence of dynamic regulation of DNA methylation in plants [2,19–26], studies in Arabidopsis indicate that for most tissues, there are only few changes in DNA methylation across different tissues and cell types [19,27,28]. Similar observations have been reported in maize [4,29,30], suggesting DNA methylation patterns are more stable in plants.

Studies in mouse and human cell lines showed that individual cytosines in mammalian genomes can not only be unmethylated or fully methylated, but also variably methylated [6,16]. All three types of cytosines are locally clustered, resulting in unmethylated regions (UMRs), fully methylated regions (FMRs), and low methylated regions (LMRs) [6,16]. UMRs and LMRs contained more than, or less than 30 CGs, respectively. Therefore, in mammals, by definition, UMRs represent regions with up to 10% median methylation and more than 30 CGs, LMRs regions with 10 to 50% methylation and less than 30 CGs, and FMRs regions with more than 50% median methylation [6,16]. UMRs are mostly stable across tissues, CG-rich and found at promoter proximal regulatory sites. LMRs, however, are more cell-type specific, CG-poor, located distantly from transcription start sites (TSSs), and enriched in active regulatory regions [6,16,31]. LMRs are indicated to be due to DNA demethylation, which is initiated by the binding of protein factors [6] To identify UMRs and LMRs based on percent methylation and number of CG dinucleotides in a genome-wide manner, Burger et al [16] developed the MethylSeekR package. We used this package to identify intergenic candidate enhancers in maize [4]. The candidate regions were characterized by high chromatin accessibility, H3K9 acetylation and at maximum 20% DNA methylation in CG and CHG context, and included known enhancers. In this study, we analyse DNA methylation data from several plant species with MethylSeekR, and show that UMRs, LMRs and FMRs can also be clearly distinguished in plant genomes. We explored the characteristics of plant UMRs and LMRs, and discovered that in plants, the majority of CG methylation is either present in UMRs or FMRs, while CHG methylation can be classified into UMRs, LMRs and FMRs. CHG LMRs consisted mostly of low methylated CCG sites. Reminiscent of mammalian CG UMRs, CG and CHG UMRs are mostly shared between tissues. Unlike in mammals, distal candidate regulatory sequences in plants appear to overlap with UMRs instead of LMRs.

## Results

### In plants LMRs are abundant in CHG context and not in CG context

To examine if distinct UMRs, LMRs and FMRs can be distinguished in plant genomes, we explored the genomes of four selected plant species, two dicots, *Arabidopsis thaliana*, *Solanum lycopersicum* (tomato), and two monocots, *Oryza sativa* (rice) and *Zea mays* (maize). These species differ greatly in genome size (ranging from about 135 Mb to 2,500 Mb), and transposable element (TE) content (ranging from about 20% to 85%), and hence the DNA fraction that is methylated [1,32–34]. We used MethylSeekR [16] to explore the existence of UMRs, LMRs and FMRs in all three DNA methylation sequence contexts observed in plants, CG, CHG and CHH. UMRs and LMRs were identified, and FMRs were defined as the methylated regions not being either UMR or LMR. On the basis of our analysis of the distribution of α-values for chromosomes 1 and 2 of all four species, we decided not to apply PMD filtering (see Methods).

MethylSeekR [16] first identifies the presence of consecutive CG dinucleotides with no or low DNA methylation, and then defines the identified regions as UMRs or LMRs if the number of CGs in the regions is at least, or less than 30, respectively. UMRs and LMRs from human tissues show two clear clusters (Additional file 1: Figure S1, reprinted from Burger et al. [16]); the cluster on the left was defined as containing LMRs and spanned from 0 to 30 CGs (x-axis, 1 to 5 in log scale), and 0 to 50% median methylation (y-axis). The cluster on the bottom right spanned from 30 to 1,024 CGs, and 0 to 10% median methylation, and was defined as containing UMRs. FMRs were regions with above 50% median methylation. Publicly available bisulphite (BS)-seq data sets of different tissues of the four plants were used: Arabidopsis wild-type Columbia seedlings [35], leaves from 3-week old plants [36], and immature fluorescence [37], tomato leaves and green fruits [2,38], rice leaves [39] and maize B73 coleoptile tissue [40]. The distribution in median methylation and number of CGs were compared between datasets using plots obtained from MethylSeekR [16].

In the plant species and tissues examined, for CG methylation, clusters corresponding to the UMR clusters in mammals were observed (Fig. 1, Additional file 1: Figure S2 and S3), and only relatively scarce regions with low DNA methylation (LMRs). There were 2 to 12 times more CG UMRs than CG LMRs (Table 1). On the other hand, for CHG methylation, clusters corresponding to UMRs, and clusters with LMRs were observed. Similar to mammalian data on CG LMRs and UMRs, for plants there are more CHG LMRs than UMRs (1.5 to 10 fold more, Table 1, Additional file 3: Table S2)[6,16]. As for CG sites, CHH methylation data showed mostly one cluster, corresponding to UMRs (Fig. 1, Additional file 1: Figure S2, S3). This work further only focused on CG and CHG methylation, as CG and CHG methylation comprise the majority of DNA methylation observed in plants, and CHH methylation only a minority [1,38,41,42]. Furthermore, sequence regions enriched for CHH are already elaborately studied by others (see e.g. [13,14,29,41,43,44].

**Figure 1.**
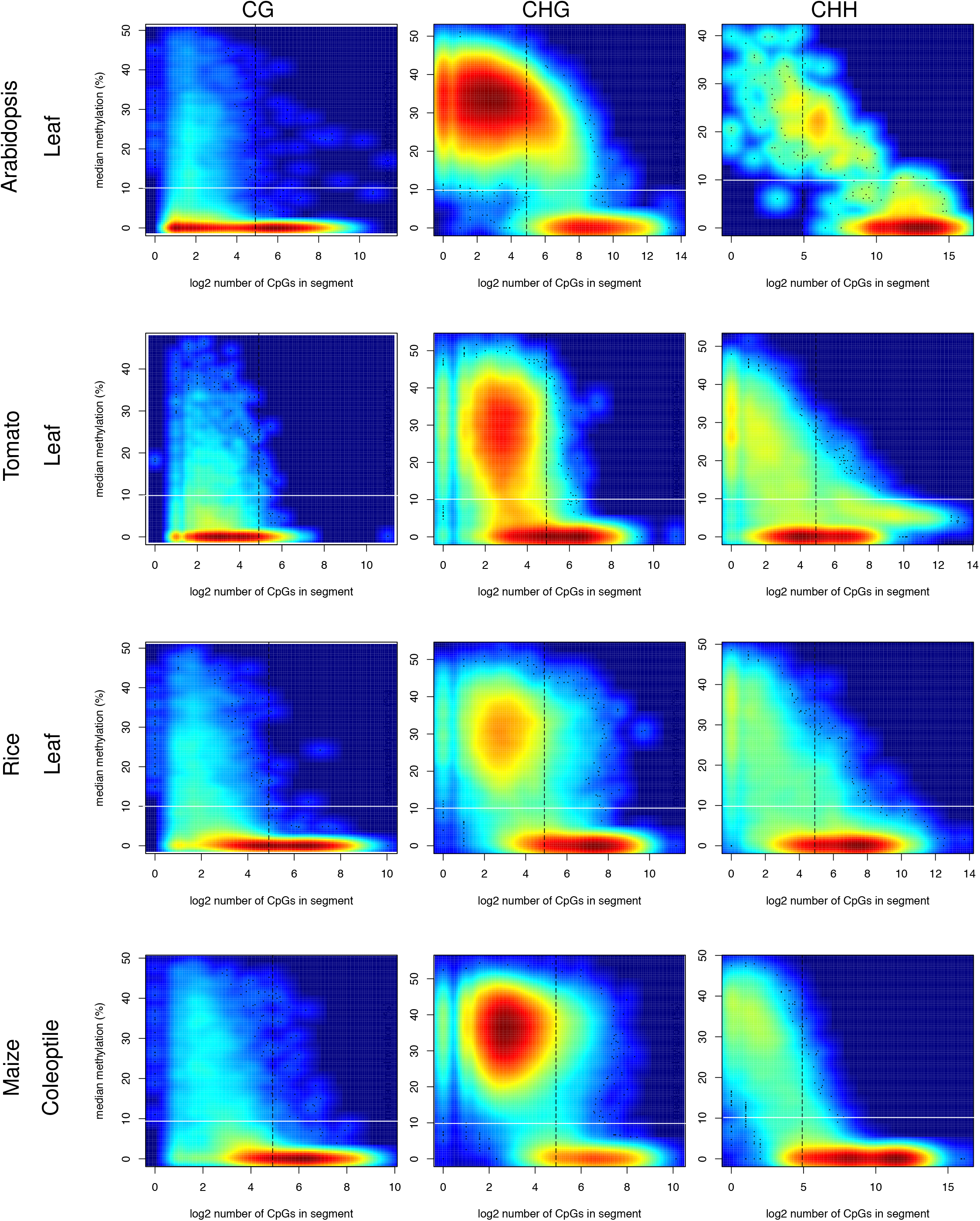
Number of CpGs against median methylation levels for CG, CHG and CHH methylation context up to 50% median methylation in Arabidopsis leaf (genome-wide), tomato leaf, rice leaf and maize coleoptile (chromosome 1) bisulfite data sets. The black dashed vertical lines indicate 30 CpGs, which were used to classify UMRs and LMRs in mammalian data sets using MethylSeekR [16]. The white solid horizontal lines indicate the 10% median methylation boundary chosen in this study to distinguish UMRs and LMRs in plants. Low to high density of data points is indicated by colour (blue to red, respectively); the colour scales are different for each dataset and not comparable between datasets.

**Table 1.** Number of LMRs and UMRs in the selected plant genomes.

| Plant tissue | CG |  |  | CHG |  |  |
| --- | --- | --- | --- | --- | --- | --- |
|  | UMRs | LMRs | ratio | UMRs | LMRs | ratio |
| Arabidopsis seedling | 16,162 | 2,254 | 7:1 | 2,046 | 19,630 | 1:10 |
| Arabidopsis immature inflorescence | 38,212 | 3,138 | 12:1 | 3,909 | 25,128 | 1:6 |
| Arabidopsis leaf | 33,985 | 2,947 | 11:1 | 2,894 | 19,545 | 1:7 |
| Tomato leaf | 75,758 | 13,336 | 6:1 | 48,422 | 76,682 | 2:3 |
| Tomato green fruit | 63,948 | 33,008 | 2:1 | 42,986 | 87,959 | 1:2 |
| Rice leaf | 115,966 | 15,245 | 8:1 | 61,200 | 90,176 | 2:3 |
| Maize coleoptile | 95,869 | 14,493 | 7:1 | 77,700 | 460,329 | 1:6 |

Consistent with the 2D density MethylSeekR plots, histograms displaying the frequency of regions with specific percentages CG and CHG median methylation showed a clear distribution of DNA methylation into UMRs, LMRs, but also FMRs (Additional file 1: Figure S4). In most tissues, a large part of the CG and CHG sites were present in FMRs. Also plotting of the MethylSeekR data in heatmaps using R indicated that plant genomes can be segmented in three distinct classes, UMRs, LMRs and FMRs (Additional file 1: Figure S5). The data in addition showed clear LMR peaks for the CHG, but not the CG data sets. In conclusion, plant genomes can be divided in UMRs, LMRs and FMRs; UMRs were present in CG and CHG DNA methylation contexts, while LMRs were mainly present in CHG context.

In MethylSeekR, UMRs and LMRs are classified based on the number of CG dinucleotides in the segments, as this criterion clearly identified two clusters in human cells [16] (Additional file 1: Figure S1). However, when compared to human data, in plants the UMR clusters generally showed a larger range in the number of CpGs, while the CHG LMR clusters showed a narrower distribution of median methylation, (Fig. 1, Additional file 1: Figure S1, S2, S3; [16]). This difference in distribution makes the classification to distinguish UMRs and LMRs by the number of CGs in segments less favourable. Therefore, as we observed a local minimum at 10% median methylation (Additional file 1: Figure S4), for plants we chose to define UMRs as regions with a median methylation below 10%, and LMRs as regions with 10% to 50% median methylation, and not take the number of CGs into account.

As also reported by Burger et al. [16], during the analyses we noticed that to observe clear clusters of UMRs and LMRs, sufficient coverage of sequencing reads (higher than 10x) was required. The clear separation of UMR and LMR clusters was not observed using data with an average coverage lower than 10-fold (Additional file 1: Figure S6, Additional file 3: Table S1).

### CG and CHG UMRs are preferentially located on chromosome arms, while CHG LMRs were more concentrated at repeated sequences

To examine the chromosomal distribution of CG and CHG UMRs, LMRs and FMRs, their chromosomal locations were studied. Chromosomes 1, in all species and tissues examined, were binned into 0.1 Mb (Arabidopsis and rice) or 1 Mb windows (tomato and maize), depending on the genome size. For each window, the coverages of C(H)G U/L/FMRs were calculated using bedtools [45] and plotted using R (Fig. 2, Additional file 2: Figure S7 and Figure S8). A coverage of 100% means the window was completely covered by the specific region investigated. The analysis revealed that for all samples examined, as expected CG and CHG UMRs were mainly located at chromosome arms, where most of the genes are located [3,32,38,46]. The coverage of CG LMRs was low throughout all analysed chromosomes 1, consistent with only a small portion of the genome consisting of CG LMRs (Fig. 2, 3b and Additional file 2: Figure S9). Intriguingly, CHG LMRs are mainly localized in pericentromeric regions in Arabidopsis and more throughout the genome for tomato, rice and maize (Fig. 2, Additional file 2: Figure S7). When examining the chromosomal distribution of CG and CHG FMRs, a high coincidence with the presence of transposable elements was observed (Additional file 2: Figure S8). This was expected since repeated sequences are generally hypermethylated [47,48].

**Figure 2.**
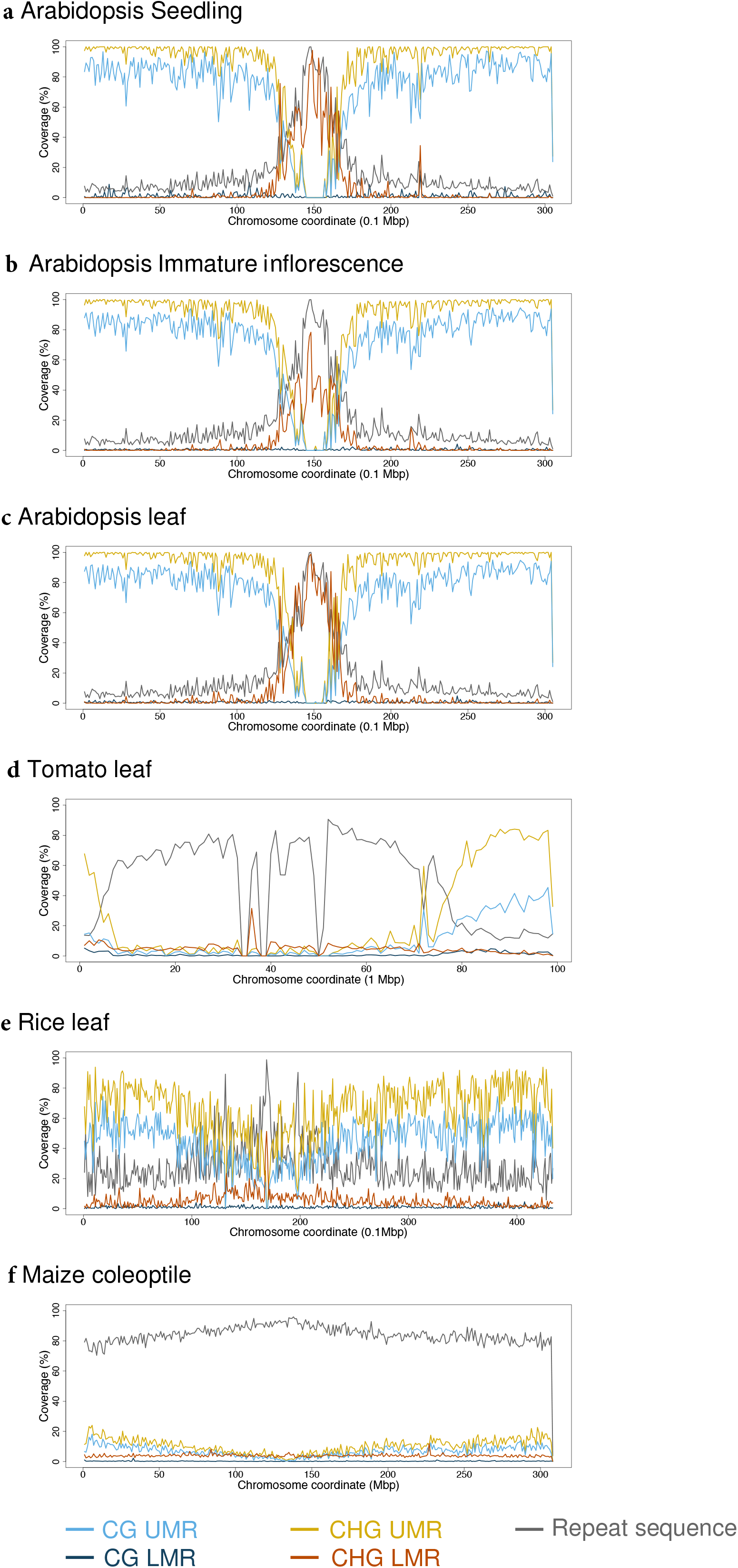
Coverage of LMRs and UMRs over windows in chromosome 1. Arabidopsis **a** seedling, **b** immature inflorescence, and **c** leaf, **d** rice leaf at 0.1 Mbp, **e** tomato leaf, **f** maize coleoptile data at 1 Mbp. The colour code shows UMRs and LMRs in different DNA methylation contexts and repeated sequences. A tomato CHG LMR peak around the centromere has been removed as the sequence at the peak was unknown (N’s).

### UMRs are enriched in genic regions, LMRs in intergenic regions

In mouse and human cells, UMRs are often located at proximal promoters and LMRs in intergenic regions distal from genes [6,16]. To examine the distribution of plant UMRs and LMRs, their enrichment in different types of genomic regions was examined. The genomic regions were defined as in Fig. 3a, and the total occupancy of UMRs and LMRs in each genomic region, in both the CG and CHG context, was calculated in kb (Fig. 3b and Additional file 2: Figure S9). In addition, to visualize the enrichment or depletion of UMRs and LMRs in particular genomic regions, the fraction of UMR and LMR occupancy in each genomic region was calculated compared to the total length of the respective genomic regions (Additional file 2: Figure S10). This analysis revealed that, as expected [3], CG and CHG UMRs were generally enriched in genic and immediately flanking regions, and depleted from distal regions and TEs. We in addition show that CG LMRs were enriched in genic sequences and depleted in TEs, while CHG LMRs were enriched in distal regions and TEs, and sometimes flanking regions. The observations for CG and CHG UMRs, and CHG LMRs are consistent with the observations in mammals [6,16]

**Figure 3.**
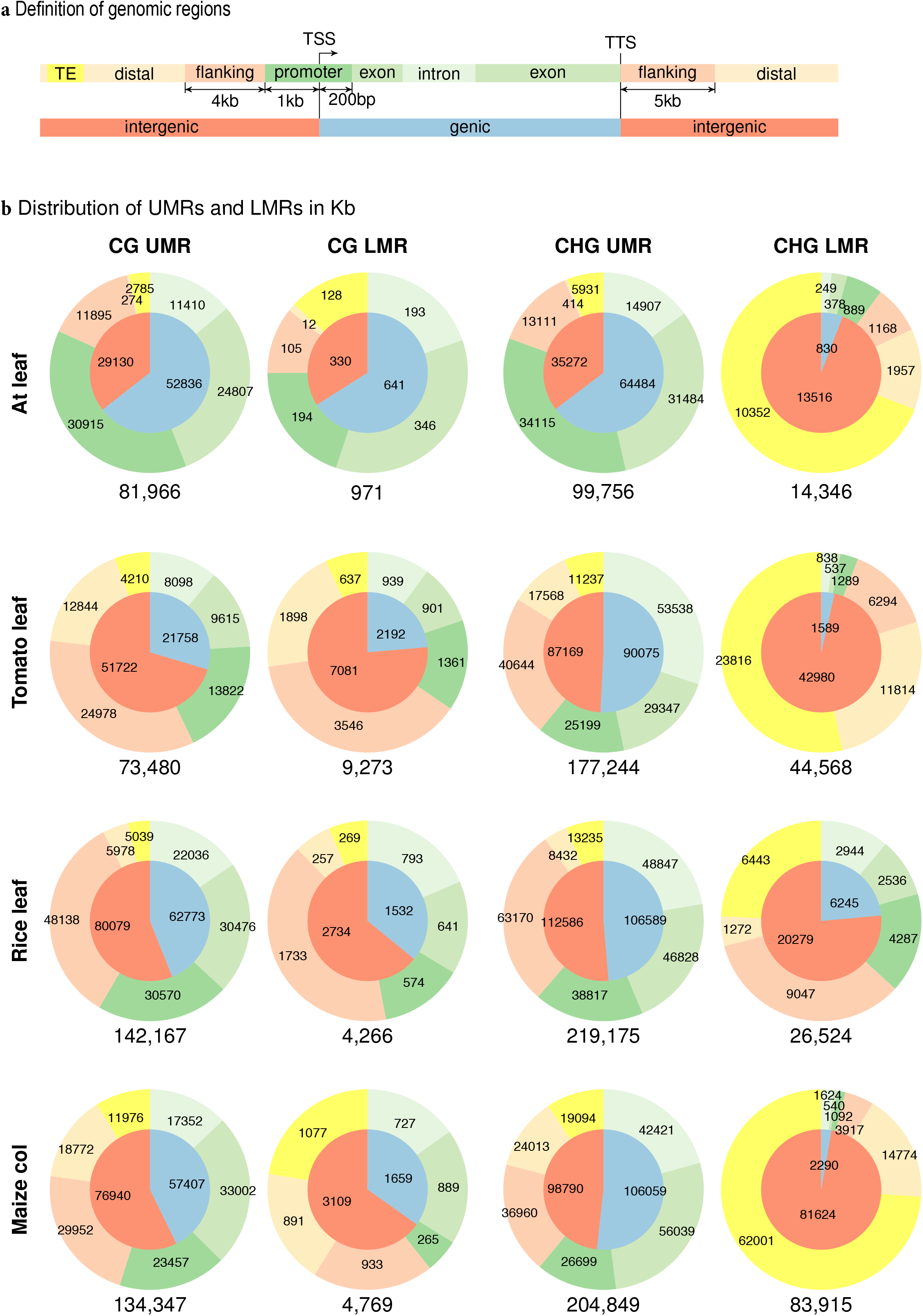
LMR and UMR distribution over genomic regions. **a** Definition and colour key for the genomic regions. **b** LMRs and UMRs among different genomic regions in the studied plant species. At stands for *Arabidopsis thaliana* and col for coleoptile. The numbers in and below the pie charts indicate occupation of each genomic region in kb.

To investigate if certain TE superfamilies were enriched for CHG LMRs in maize, TE superfamilies with more than 100 members in the maize genome were analysed in terms of the proportion of CHG LMRs covering the total length of given TE superfamilies. This analysis revealed a high enrichment of CHG LMRs at long terminal repeat retrotransposon (LTR) gypsy families (RLG), and at terminal inverted repeat (TIR) DTA families (Fig. 4a).

**Figure 4.**
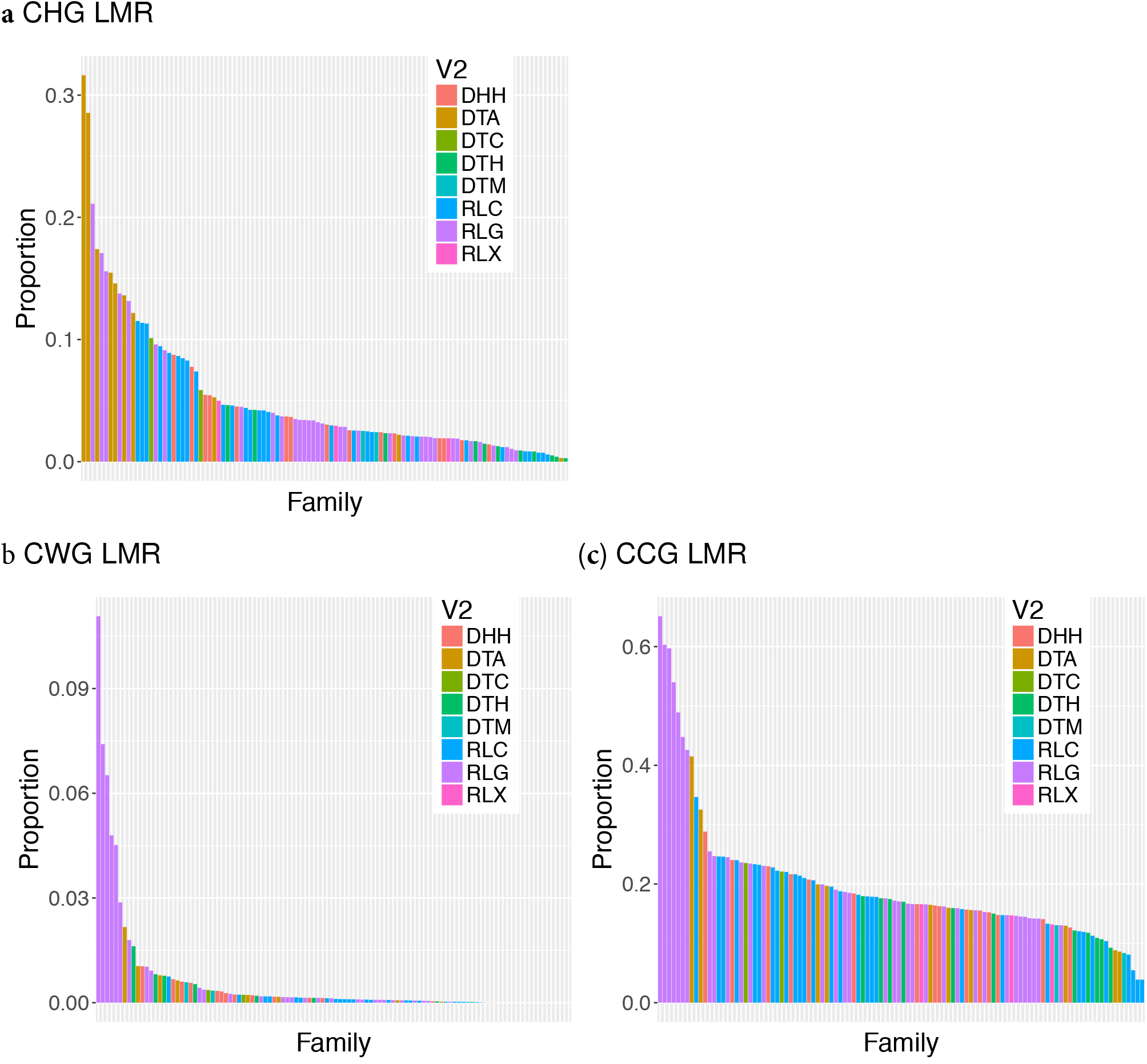
Enrichment of (A) CHG, (B) CWG and (C) CCG LMRs in TE families with more than 100 family members in the maize genome. The y-axis indicates the proportion of the total TE length of TE families covered by CHG/CWG/CCG LMRs. The colours indicate different superfamilies of TEs.

### UMRs are located at TSSs, LMRs distal from genes

To investigate the location of UMRs and LMRs in more detail, the distribution of distances between UMRs or LMRs and their closest TSS were studied by calculating the distances in base pairs and plotting them in histograms (Fig. 5 and Additional file 2: Figure S11). Relatively large fractions of both CG and CHG UMRs (20-66 % and 33-92 %, respectively) were overlapping with TSSs, whereby the fractions were highest for Arabidopsis and lowest for rice and maize. These data clearly indicate a preferential occupation of both CG and CHG UMRs at TSSs and is consistent with earlier observations that TSSs are generally unmethylated [3,40,49], and can be explained by the requirement of TSSs and promoter regions to be unmethylated to allow transcription initiation [50,51]. The percentages of LMRs overlapping TSSs were generally much lower (between 0.1 and 3%), with a few exceptions (CG LMRs of Arabidopsis seedling (9.35%) and tomato (7.60-9.92%), and CHG LMRs of tomato leaf (23.71%; Additional file 2: Figure S11). CHG LMRs were generally located further away from TSSs (Additional file 2: Figure S11), in distal regions and TEs (Additional file 2: Figure S10).

**Figure 5.**
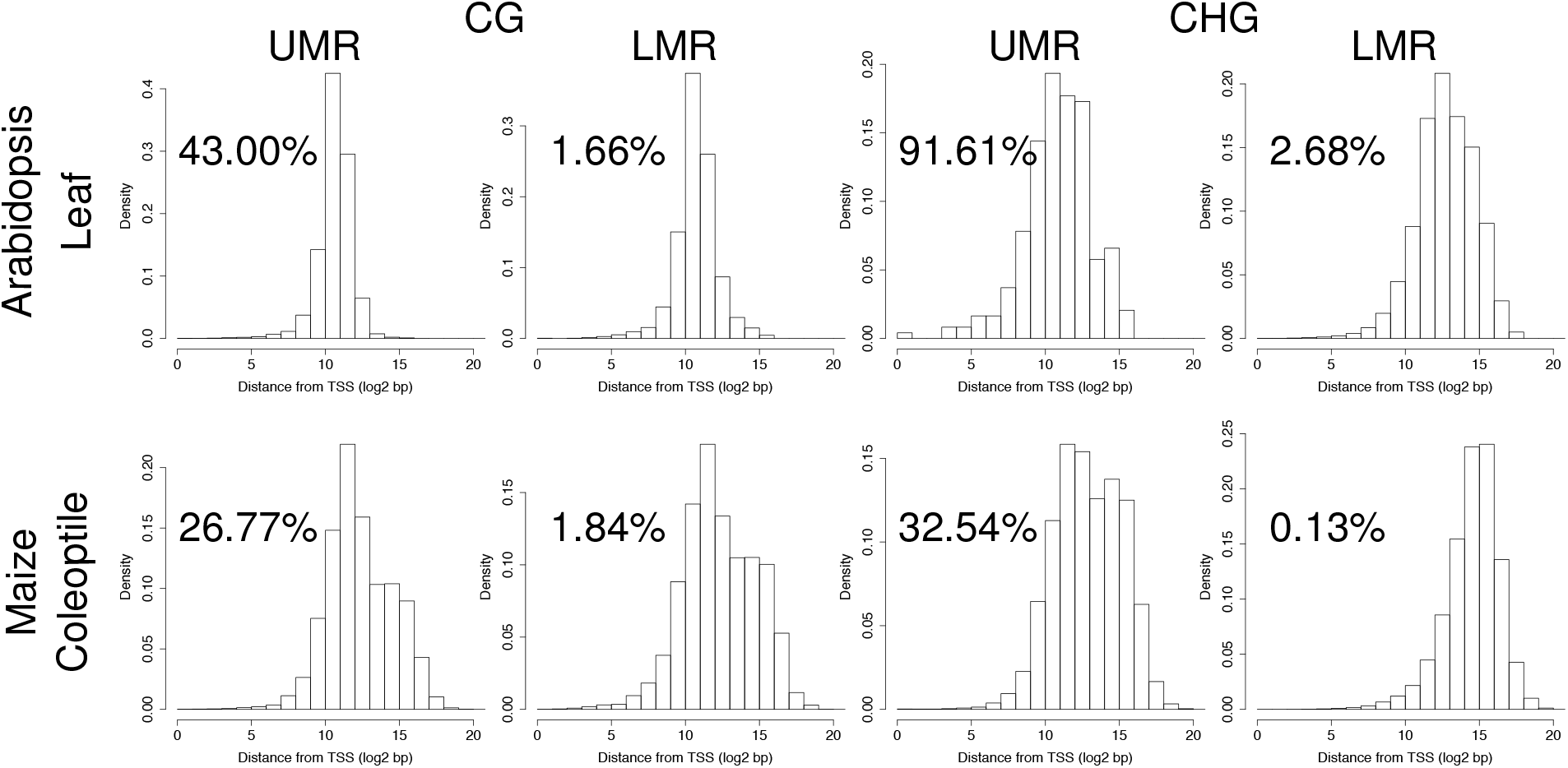
Distribution of distances between UMRs or LMRs and their closest TSSs in log2 scale for Arabidopsis leaf and maize coleoptile. The UMRs or LMRs with 0 distance from a TSS (i.e. overlapping with a TSS) were excluded from the figures, but their percentages are given instead.

### Plant UMRs are generally longer than LMRs

Next, the sizes of UMRs and LMRs were explored to see if there is a trend among the species. The comparison between median size values showed that UMRs were generally longer than LMRs, in both CG and CHG context (Additional file 2: Figure S12, Additional File 3: Table S3). For example, the median sizes of CHG UMRs in Arabidopsis leaf and maize coleoptiles were 16.4 kb and 1.7 kb, respectively, while those of CHG LMRs were 241 bp and 69 bp, respectively. The trend of UMRs being longer than LMRs is also true for mammalian cells (3- and 2-fold difference in human and mouse cells, respectively; Additional file 2: Figure S12), but the differences in size were generally larger in plants.

CHG UMRs were on average longer than CG UMRs. This was expected, as coding regions are usually depleted for CHG methylation, while a significant fraction of them are methylated in a CG context [3,40,49,52,53] The CHG UMRs appeared on average much larger in Arabidopsis than in tomato, rice and maize (12-21 kb versus about 2 kb). This is likely due to the Arabidopsis genome being most gene-dense, with several genes located close to each other [54,55]. As a result single UMRs can span several genes. The longest CHG UMR in Arabidopsis leaf is located on one of the arms of chromosome 4 (4:17,714,380-18,232,340), and contains 171 genes.

### UMRs are better conserved between tissues while LMRs were more tissue-specific

Previous studies showed that in mammals 90% or more of the UMRs are shared among various cell-types, while 20-50% of the LMRs are tissue-specific [6,16,31]. To examine the stability of UMRs and LMRs over various tissues in Arabidopsis and tomato, the numbers and fractions of CG and CHG UMRs or LMRs overlapping between tissues were calculated and plotted (Fig. 6 and Additional file 2: Figure S13). If an LMR or UMR in one tissue did not overlap with an LMR or UMR in another tissue, they were given a 0 for overlap. The percentages of LMRs or UMRs with 0% overlap were indicated in the figures.

**Figure 6.**
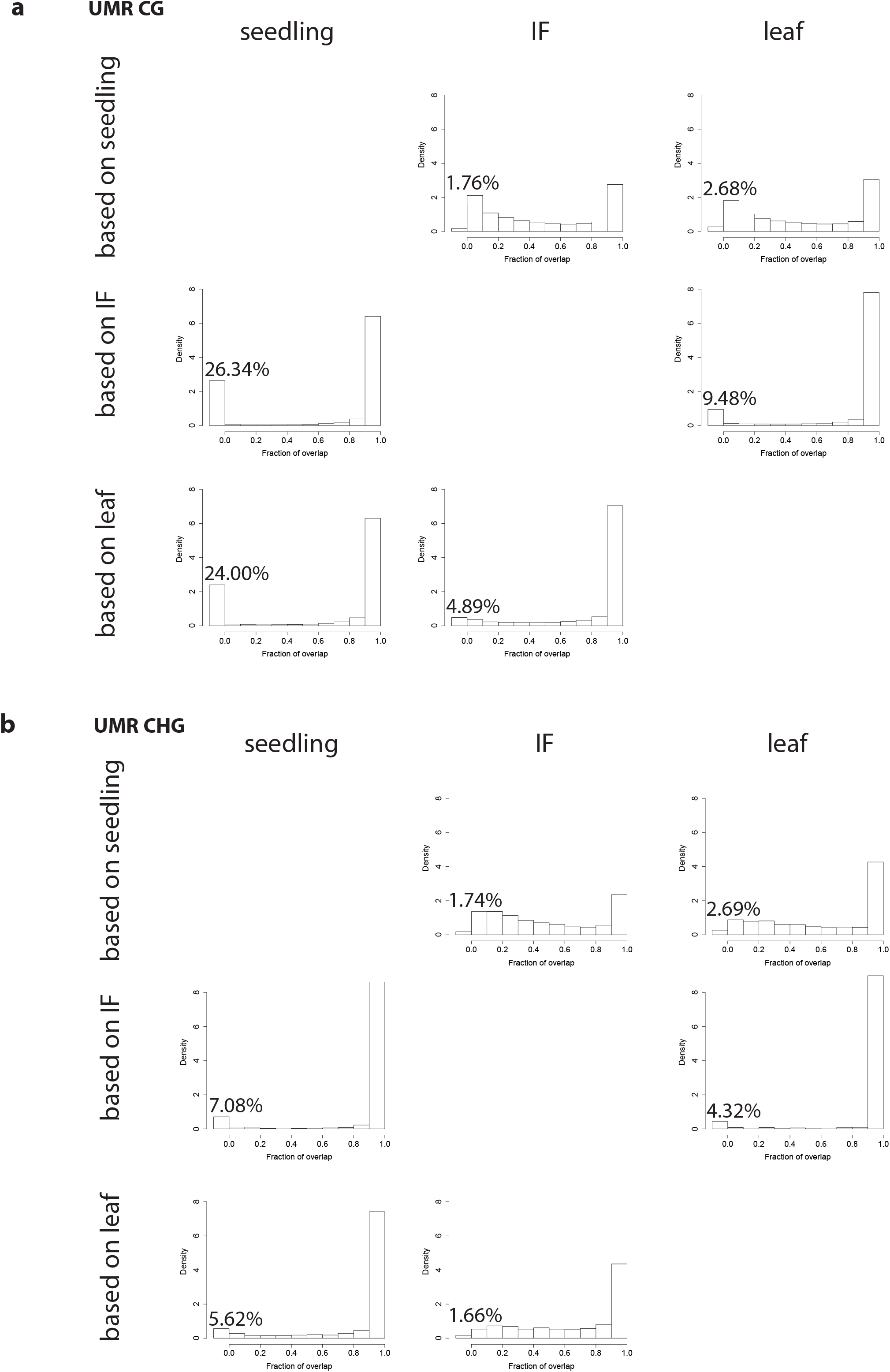
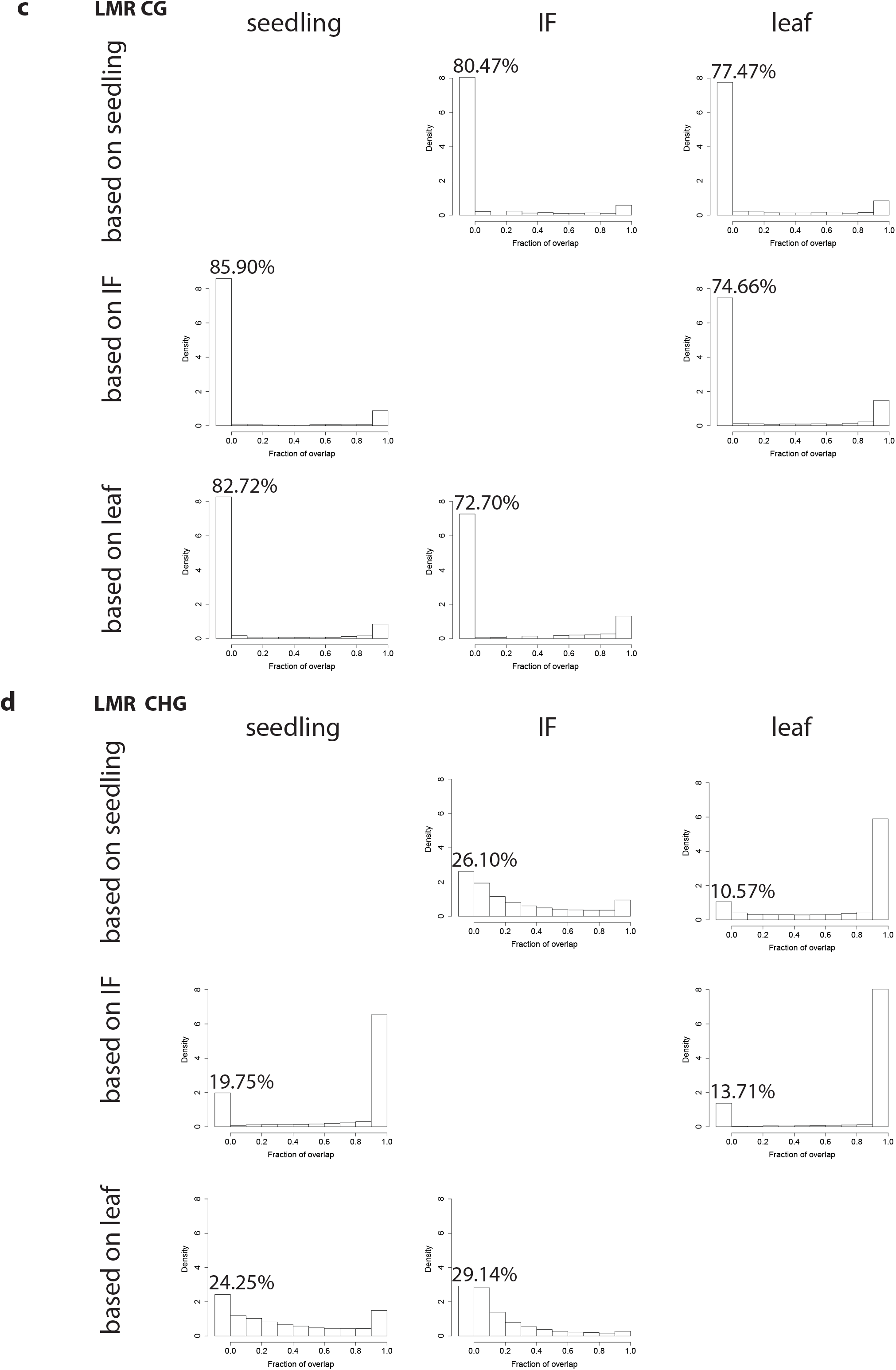
Fraction of overlapping Arabidopsis **a** CG and **b** CHG UMRs, and **c** CG and **d** CHG LMRs between tissues. The density on the y-axis indicates the fraction of UMRs or LMRs in each bin, hereby the areas of all the bars add up to 1. The percentage mentioned in each graph indicates the percentage of UMRs or LMRs without an overlap.

The 0.9-1.0 bin, corresponding to UMRs with 90 to 100% overlap, contained the highest number of data points in all comparisons, indicating that a large fraction of UMRs from different tissues overlap for more than 90% of their lengths. Relatively low percentages of the CG and CHG UMRs showed no overlap in Arabidopsis and tomato (Fig. 6 and Additional file 2: Figure S13). Strikingly, the percentages of LMRs without an overlap were clearly higher than the percentages of UMRs without an overlap. For both UMRs and LMRs, generally the percentages without an overlap were higher for CG than for CHG. Our results suggest that UMRs are generally conserved among various tissues, while LMRs are more tissue-specific. A similar trend is observed in mammalian systems [16].

### Plant CHG LMRs are associated with heterochromatic marks

To get a better insight in the possible functions of CHG LMRs, the enrichment of several chromatin features (H3K27me2, H3K9me2, H3K9ac and DNAse I accessibility) was studied by firstly calculating the average signal intensity in RPM for each UMR, LMR and FMR, followed by calculating the median values of the average signal intensities, and then plotting the median values together with the standard error using R [56]. This analysis was performed in maize using published [4,57] data sets.

UMRs in both CG and CHG context displayed high H3K9ac and DNase I accessibility, and relatively low H3K9me2 and high H3K27me2 (Fig. 7, Additional file 2: Figure S14, Additional file 3: Table S4, S5). These observations are consistent with UMRs mainly being located in genic regions, especially at TSSs (Fig. 3, 5, Additional file 2: Figures S11, S9 and S10). H3K9ac and DNase I accessibility marks active genes and regulatory regions, while H3K27me2 is among others associated with inactive genes [57–62]. FMRs, as expected, showed high H3K27me2 and H3K9me2 levels, and relatively low H3K9ac and DNase I accessibility. In plants, heterochromatin, composed of repeated sequences, is known to be associated with high levels of DNA methylation, H3K27me2 and H3K9me2 [57,63–65].

The chromatin marks at LMRs differed between CG and CHG LMRs. For CG LMRs, the levels of H3K9ac and DNase I accessibility were in between those at CG UMRs and FMRs, while the H3K9me2 and H3K27me2 levels were relatively high and more similar to those at CG FMRs. At CHG LMRs on the other hand, the levels of active marks (H3K9ac and DNase I accessibility) were as low as observed for CHG FMRs, while the levels of H3K27me2 and H3K9me2 were both relatively high, indicating inactive, inaccessible chromatin. The H3K27me2, and also H3K9me2 signal levels were, however, lower than at CG LMRs. The enrichment of CHG LMRs in TE regions (Fig. 3 and Additional file 2: Figure S10) is a possible explanation that they showed heterochromatic features and were depleted for active marks, but sharply contrasts with the observation in mouse that at least 80% of the LMRs contain DHSs and are enriched with H3K27ac, which marks active enhancers [6]. Thus, although the level of DNA methylation is similar between plant CHG LMRs and mammalian CG LMRs, mammalian CG LMRs are associated with active chromatin marks, and plant CHG LMRs are not.

**Figure 7.**
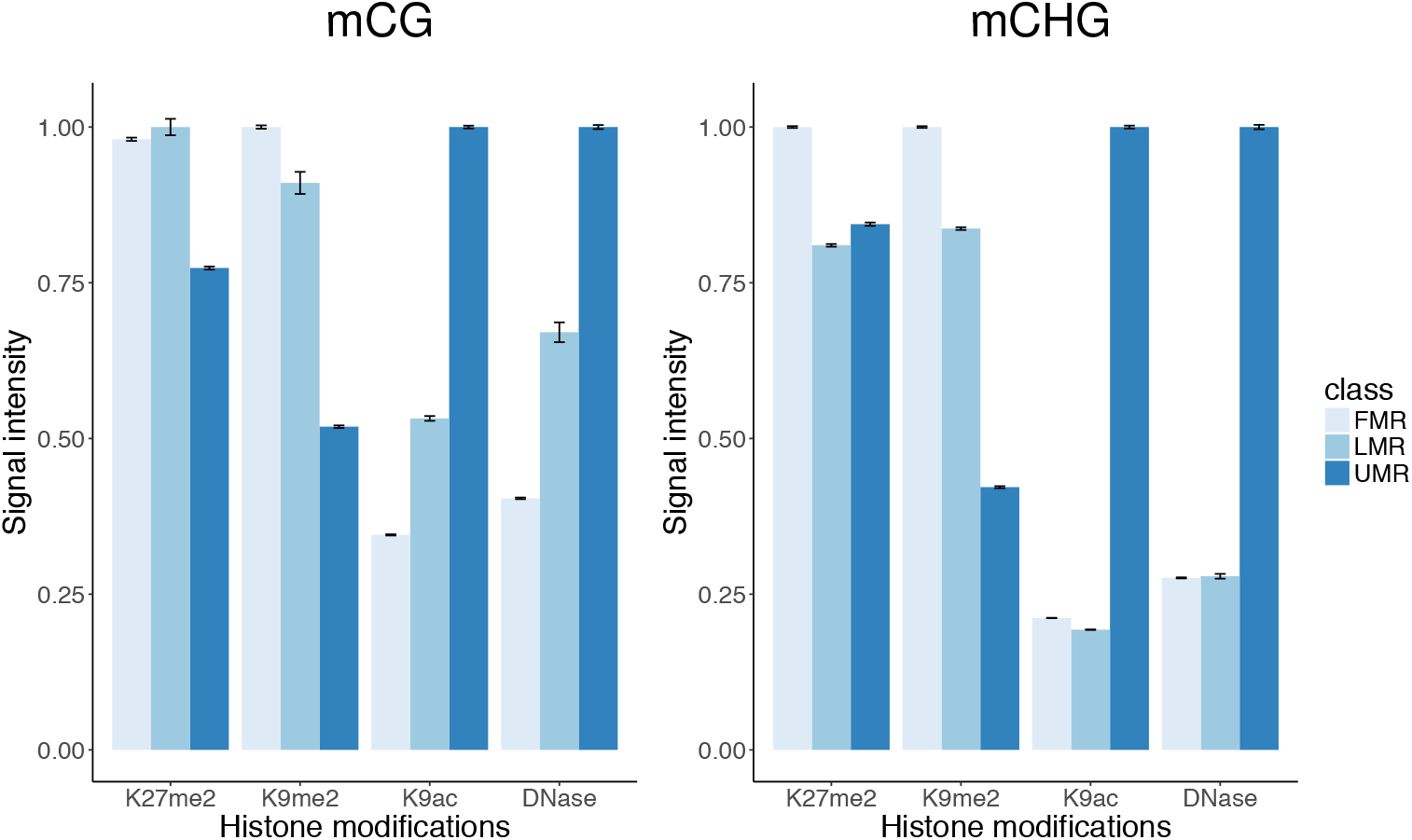
Enrichment of chromatin marks in FMRs, LMRs and UMRs in the maize genome. For each mark indicated, the average signal intensities of median signal intensities in each region were calculated, and normalised to the highest bar for each mark. H3K9me2 and H3K27me2 data is derived from inner stem tissue of 1-month-old plants (approximately V4 stage). H3K9ac and DNAse I accessibility data were derived from husk tissue.

In a previous study, we predicted approximately 1,500 intergenic candidate enhancers based on the simultaneous presence of DNAse I hypersensitive sites (DHSs), enrichment for H3K9ac and at maximum 20% DNA methylation in CG and CHG context [4]. Overlap of CHG UMRs and LMRs with DHSs and H3K9ac enriched regions indicated that the vast majority of DHSs and H3K9ac enriched regions overlapped with UMRs and only a few with LMRs (Additional file 3: Table S4). Importantly, all candidate enhancers overlapped with CHG UMRs and not with LMRs (Additional file 3: Table S4). Similar results were obtained for CG UMRs and LMRs (Additional file 3: Table S5). In addition to the ~1,500 intergenic candidate enhancers reported by Oka et al [4], recently another set of putative *cis*-regulatory elements, represented by accessible chromatin regions (ACRs), were reported by Ricci *et al*. (2019). Overlap of CHG and CG UMRs and LMRs with distal ACRs and all ACRs identified showed that also nearly all (d)ACRs overlapped with CHG and CG UMRs (Additional file 3: Table S6). Together, these results suggest that intergenic UMRs may represent most, if not all intergenic candidate regulatory sequences within a plant.

### CHG LMRs were mostly consisting of CCG LMRs

As the chromatin characteristics of plant CHG LMRs were clearly different from mammalian CG LMRs, we explored how the CHG context could be further classified and how the resulting regions were distributed over the genome. Previous studies in Arabidopsis, tomato, rice and maize showed that for CHG sites in heterochromatin, methylation of the outer C is 20-50% lower in CCG sites than in CAG/CTG (CWG) sites [38,66], suggesting that CHG LMRs may be due to low mCCG rather than mCWG methylation.

To investigate this hypothesis, the CCG and CWG methylation data were separated from each other and UMRs and LMRs were identified for each dataset using MethylSeekR [16]. Note that in this analysis, CCG methylation refers to methylation of the external C, independent of the methylation status of the inner C. The 2D distribution figures of median methylation and number of CpGs in UMRs and LMRs (Fig. 8a and Additional file 1: Figure S2) clearly show LMR clusters for CCG, but not CWG methylation, except for Arabidopsis, indicating CHG LMRs are mainly the result of low CCG methylation. The existence of CCG as well as CWG LMR clusters in Arabidopsis may be due to the generally relatively low DNA methylation level in Arabidopsis [38,67].

Analysis of the distribution of CCG/CWG UMRs, LMRs and FMRs over chromosomes and genomic regions showed that CCG and CWG UMRs were enriched at genic regions at chromosome arms and depleted at pericentromeric, heterochromatic regions, resembling the patterns of CHG, and also CG UMRs (Fig. 2, 8b, Additional file 2: Figure S15). Similar to CHG LMRs, CCG LMRs were enriched at heterochromatin, generally being highest around the centromere and declining towards the chromosome ends (Fig. 2; Additional file 2: Figure S15). In line with this, as for CHG LMRs, CCG LMRs, but also CWG LMRs, were enriched at intergenic regions, including TEs (Fig. 3, 8b, Additional file 2: Figures S9, S10). This is consistent with previously reported data that methylation of CHG, and CCG and CWG motifs is generally enriched at heterochromatin [38,40,41,67]. TE enrichment analysis showed that, in the maize genome, CCG LMRs were enriched at the Gypsy superfamily (RLG) of TEs (Fig. 4).

**Figure 8.**
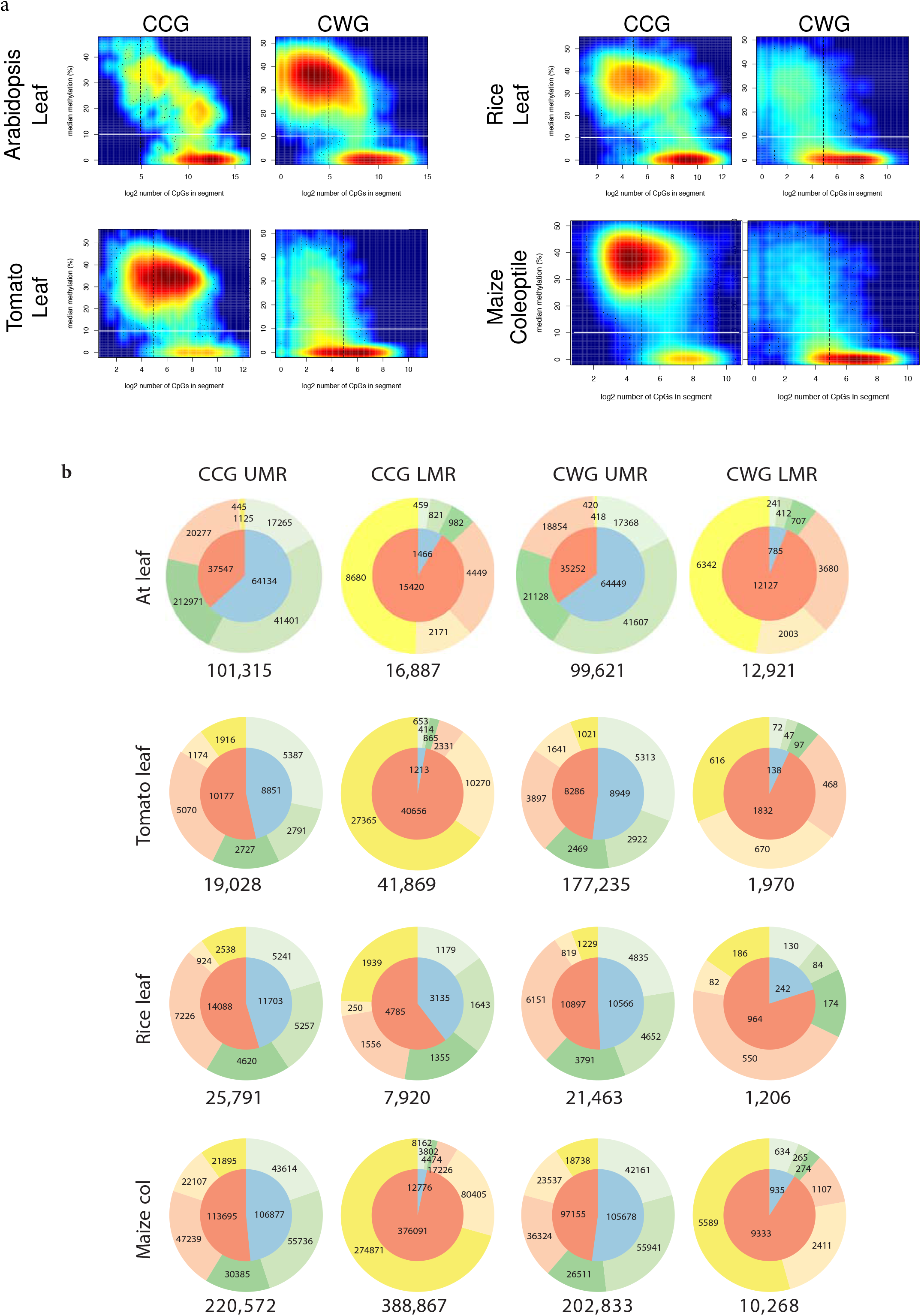
**a** Number of CpGs against median methylation levels for the CCG and CWG methylation contexts in Arabidopsis leaf (genome-wide), tomato and rice leaf, and maize coleoptile (chromosome 1) bisulfite data sets. The black dashed vertical lines indicate 30 CpGs, which were used to classify UMRs and LMRs in mammalian data sets using MethylSeekR [16]. Low to high density of data points is indicated by colour (blue to red, respectively); the colour scales are different for each dataset and not comparable between datasets. **b** UMR and LMR distribution in CCG and CWG contexts over genomic regions (see Figure 5a for definition and colour code) for Arabidopsis, tomato and rice leaf, and maize coleoptile. The numbers in and below the pie charts indicate occupation of each genomic region in kb.

CCG and CWG FMRs displayed the same coverage pattern as CCG LMRs, being enriched at heterochromatin, and decreasing in coverage towards the chromosome arms (Additional file 2: Figure S15). Furthermore, whereas the coverage across chromosomes of CWG LMRs was much lower than that of CCG LMRs, CWG FMRs showed a higher coverage than CCG FMRs.

### mCCG methylation depends on CmCG and mCWG methylation

In Arabidopsis, methylation of CWG requires CMT3, while that of the external cytosine within CCG requires both MET1 and CMT3 [38,68,69]. In a *met1* mutant, not only methylation of the internal (CmCG), but also the external C of CCG (mCCG) is depleted, consistent with methylation of the external C by CMT3 being dependent on methylation of the internal C in 5’-CCG-3’ trinucleotides by MET1. To further test this previously reported dependence, the mCCG and CmCG levels were extracted for every 5’-CCG-3’ and the data plotted in 2D density plots in log scale (Fig. 9, Additional file 4: Table S7). In general the density plots for all four species show that most of the data points are above the main diagonal (from (0,0) to (1,1)). This indicates that for most locations the level of methylation of the internal C (CmCG) is higher than that of the external C. This holds true for all analysed species, Arabidopsis, tomato, rice and maize, and is consistent with reported data [38,68] showing that the methylation of the external C is dependent on the methylation of the internal C.

**Figure 9.**
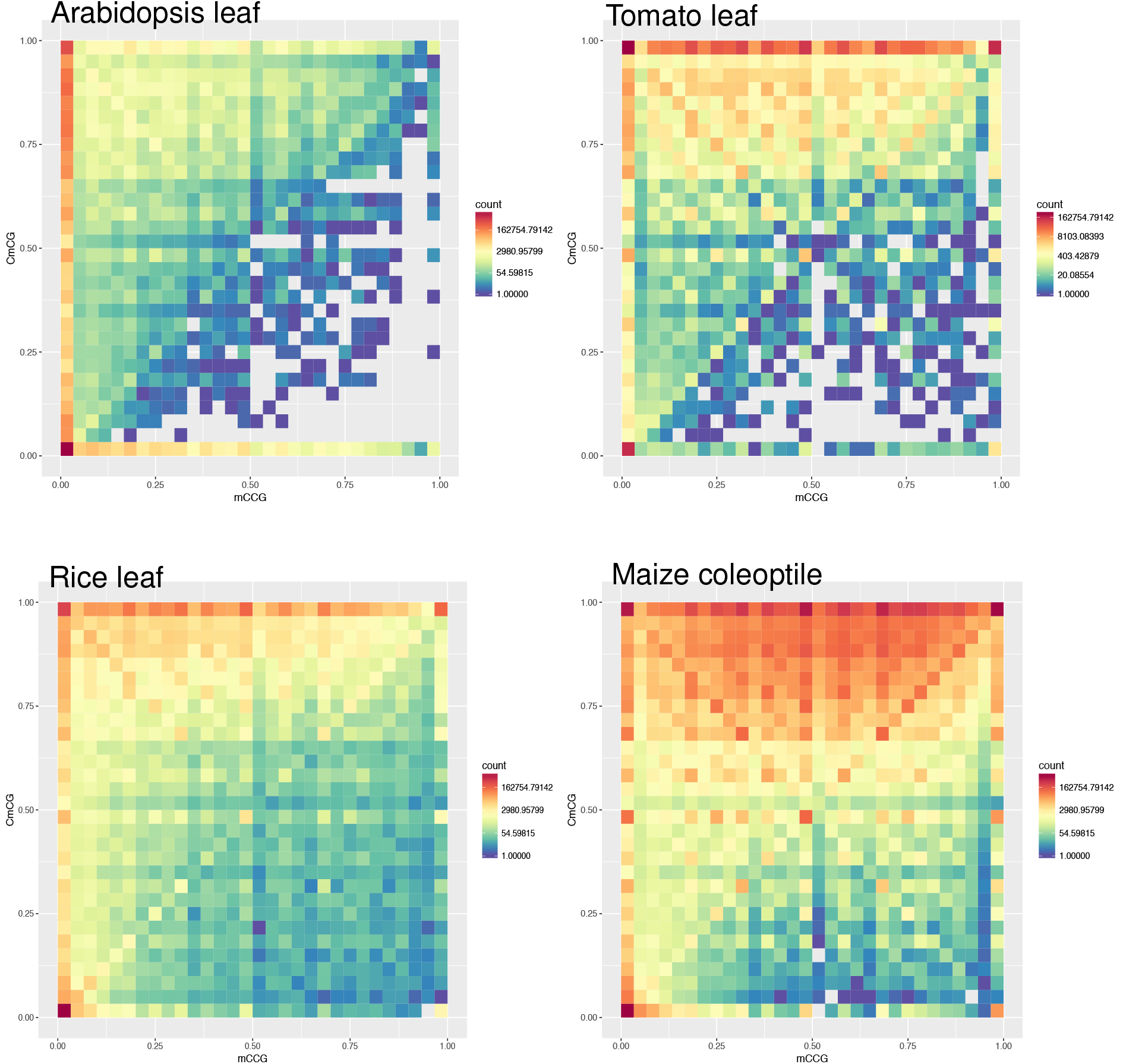
Heatmap of mCCG and CmCG counts in Arabidopsis, tomato, rice and maize. X-axis: the methylation level of the outer C in CCG (mCCG); y-axis: the methylation level of the inner C in CCG (CmCG). The colour indicates the number of occurrences.

Remarkable observations for the different species are that in Arabidopsis and rice a high fraction of external cytosines in 5’-CCG-3’ trinucleotides were fully unmethylated (96.37 and 61.27% respectively), whereby a relatively high fraction of CCG sites had both cytosines fully unmethylated (66.35 and 45.91% respectively); In tomato and maize, species with bigger genomes, these percentages are much lower. In tomato a high fraction of the internal cytosines, 70.91%, were fully methylated, while in maize, with an about 2.5 times bigger genome than tomato, about 43.20% of the internal cytosines within CCG sites were fully methylated. In both species in about 8% of the CCG sites both cytosines were fully methylated.

Considering the molecular mechanism of CCG methylation in heterochromatin, we hypothesized that the CCG methylation level may be positively correlated with the nearby concentration of CWG sites. If true, the nearby concentration of CWG sites in FMRs should be higher than in LMRs. To test this hypothesis, the concentration of CWG sites in every CCG LMR, FMR, but also UMR were determined for maize (Additional file 2: Figure S16). This analysis revealed that the median CWG concentrations in CCG UMRs, LMRs or FMRs were similar among each other, whereby the distributions were narrow for CCG UMRs and broad for CCG LMRs and FMRs. No dependency of the mCCG methylation level on the nearby CWG concentration was observed.

It was postulated that H3K9 dimethylation, by the binding of H3K9 methyltransferases to methylated CWG sites, would enhance the recruitment of CMT3 to neighboring CCG sites [38]. If true, one would expect a relation between the methylation levels of overlapping CWG and CCG regions. To test this hypothesis, the overlap in the number of base pairs for CCG and CWG UMRs, LMRs and FMRs were calculated for maize. Results are presented in Additional file 3: Table S7A. Calculating the percentages of basepairs in the three different CWG regions indicated that 9.64%, 0.49% and 89.87% of the genome is covered by CWG UMRs, LMRs and FMRs, respectively (Additional file 3: Table S7B, second column). If the methylation levels of the CWG and CCG regions have no relation with each other, the expected percentages for the overlapping regions would be the same as indicated for the CWG regions. However, although only 9.64% of the base pairs are contained in CWG UMRs, 80.63% of the 221 million base pair in CCG UMR regions is overlapping with the 203 million base pairs in CWG UMRs (8.4 fold enrichment), indicating a positive relation between the lack of methylation in CWG and CCG regions. The number of base pairs in CCG LMRs overlapping with CWG LMR regions are 3.6 fold higher than expected (1.75% instead of 0.49%), indicating a positive correlation. The same is true for CCG FMRs and CWG FMRs. These results indicate a positive correlation between the methylation level of a CWG region and that of a CCG region, which is in line with the hypothesis stated by Gouil et al [38].

## Discussion

In this study we show that MethylSeekR is a useful tool to identify unmethylated DNA regions (UMRs), which for large, highly methylated genomes greatly facilitates the identification of the *cis*-regulatory genome. The vast majority of the active genome and candidate enhancers in maize identified in previous studies [4,5] are contained in the unmethylated part of the genome.

Previous studies in mammalian systems revealed three types of DNA methylated regions, UMRs, LMRs and FMRs, and showed that UMRs and LMRs generally represent proximal regulatory elements and active distal regulatory elements, respectively [6,16]. In mammals, UMRs are mostly shared among tissues, whereas LMRs are more cell-type specific. Here, we studied if in plants similar DNA methylation categories can be distinguished. We found that plant CG methylation mainly classified in two clusters, corresponding to UMRs and FMRs; the LMR cluster is lacking (Fig. 1, Additional file 1: Figure S2, S3 and S5). CHG methylation, however, did show the three type of clusters found in mammals, UMRs, LMRs and FMRs. To our knowledge, LMRs have not been described in plants before. CHH methylation, representing a minor fraction of the DNA methylation within plant genomes [38,40,42], behaved mostly like CG methylation, displaying mainly UMR and FMR cluster. Further context classification of CHG into CCG and CWG showed that the main contributor to the LMR clusters was CCG and not CWG methylation (Figure 8A), consistent with the 20-50% lower methylation levels of CCG motifs compared to CWG motifs observed by Gouil and Baulcombe [38].

Our MethylSeekR and density plots showed that, to identify plant UMRs and LMRs, drawing a line at 10% median methylation was best, as there was a clear local minimum in the number of regions at 10% (Additional file 1: Figure S4). To distinguish UMRs and LMRs in mammalian data, the presence of 30 CpGs within a region was selected as a threshold [6,16], with LMRs and UMRs being characterized by less than, or at least 30 CpGs, respectively. Although also in plants LMRs on average contain less CpGs than UMRs, the distributions of LMRs and UMRs appeared to overlap to some extent, hampering the use of a specific number of CpGs as a threshold to identify the two types of regions.

Although the sequence context of LMRs differs between mammals and plants, CG versus CHG, plant LMRs, but also UMRs, still have a lot of features in common with those of mammals [6,16]: i) LMRs are enriched at intergenic sequences, UMRs in genic and immediate flanking regions, ii) LMRs are on average shorter than UMRs, iii) LMRs are on average further away from TSSs than UMRs, and iv) A significant fraction of LMRs is tissue-specific, while UMRs are mostly shared between tissues. As pointed at above, plant LMRs do however also show differences with mammals: i) plants contain CHG LMRs instead of CG LMRs, and ii) CHG LMRs are associated with heterochromatic marks instead of accessible chromatin. CHG methylation is maintained by CMT3, which is a plant-specific enzyme [1,8], which could indicate a different function of plant LMRs compared to mammalian LMRs. At the same time, it could not be excluded that plant and mammalian LMRs, although mediated by diversified enzymes have similar functions.

CG and CHG UMRs were generally enriched at genic and immediate flanking regions (Fig. 3 and Figure S9), and compared to LMRs more frequently located at TSSs (Fig. 5 and Additional file 2: Figure S11). TSSs and their flanking regions have indeed previously been shown to be hypomethylated [3,40,49,52]. Hypomethylation at these regions is in line with the role of proximal regulatory sequences, as transcription initiation is hindered by DNA methylation [50,51] (62, 63). We further showed that a significant fraction of UMRs is associated with H3K9ac and DNAseI accessibility, and that the other way around, nearly all accessible and H3K9ac enriched regions overlapped with UMRs instead of LMRs (Additional file 3: Table S4, S5). Moreover, all intergenic candidate enhancers we reported before [4] overlapped with CG and CHG UMRs and not with LMRs (Additional file 3: Table S4, S5). Similarly, the vast majority of the recently reported distal ACRs, a fraction of which is indicated as putative *cis*-regulatory elements [5] overlapped with UMRs. Importantly, there are many more intergenic UMRs than the ones overlapping with intergenic candidate enhancers. We expect these remaining intergenic UMRs to represent regulatory sequences in other tissues than examined [4] Consistent with this idea, UMRs are highly conserved among various tissues (Fig. 6 and Additional file 2: Figure S13). We therefore propose that, as also postulated by Crisp et al. [28], the genome-wide identification of UMRs will be a very useful approach to find the vast majority of the *cis*-regulatory sequences, especially in plants with large, highly methylated genomes. MethylSeekR is an excellent tool to find these UMRs.

As also observed in mammals [6,16], in all plant species studied, UMRs were generally longer than LMRs, whereby CHG UMRs were longer than CG UMRs. The latter can be explained as, although TSSs are generally depleted of both CG and CHG methylation, coding regions are generally only depleted of CHG methylation; they often carry gene body methylation in the CG context [3,11,40,49,52]. Thus CHG UMRs often span both the TSS and entire coding region, while CG UMRs do not. The median sizes of CHG UMRs in Arabidopsis were much larger than those in the other plant species (~12-21 kb versus ~2 kb) (Additional file 2: Figure S12). This is likely due to the high gene density in Arabidopsis compared to the other species [54,55,70,71], resulting in CHG UMRs spanning several closely located genes. In conclusion, our observations on plant UMRs indicate that in general they have a very comparable function as mammalian UMRs.

As observed for mammalian LMRs, CHG LMRs showed relatively high levels of tissue-specificity, and were enriched at intergenic regions, primarily distal and TE regions (Figs. 3 and 6, Additional file 2: Figure S9, S10, S13). In mammals, CG LMRs are associated with active distal regulatory sequences and result from the binding of transcription factors, which directly or indirectly trigger active DNA demethylation by TET enzymes, activating the regulatory elements [6,17,18]. Although the enrichment of LMRs at distal regions fits the observations in mammals [6], the enrichment at TE regions may be less expected, as TEs usually display high DNA methylation and transcriptional repression [72,73]. TE sequences have, however, been indicated as a source of regulatory activity [4,74–78]. In accordance, about 33% and at least 80% of mouse CG LMRs overlap with repeated DNA, and accessible chromatin, respectively [6]. We, however, showed that CHG LMRs in maize plants are enriched for inactive chromatin marks, H3K9me2 and H3K27me2, and depleted for active chromatin marks such as H3K9ac and DNase I accessibility (Fig. 7 and Additional file 2: Figure S14). In line with these results there is also no significant overlap with candidate enhancers identified in maize [4] (Additional file 3: Table S4, S5, S6). Instead, the candidate enhancers in Oka et al [4] overlap for 100% with UMRs. About 60% of these candidate enhancers is further away than 5 kb to the closest flanking gene [4], and indeed, although UMRs are enriched at genic regions, a fraction of the UMRs are located at distal regions.

Our data indicate a difference in the processes involved in activating regulatory sequences in plants versus mammals In mammals distal regulatory elements are demethylated upon activation, generating LMRs [6]. Our results indicate that in plants, generally no active DNA demethylation is required to activate regulatory sequences. Still, a small fraction of the UMRs is tissue-specific (Fig. 6 and Additional file 2: Figure S13), in line with studies showing a role for active DNA demethylation during a.o. specific developmental processes or plant responses to pathogens [2,79–83]. For example, in tomato, active cytosine demethylation in the promoter of particular genes plays a crucial role in fruit ripening [2,79], whereas in Arabidopsis cytosine demethylation at TEs/repeats-associated defense-related genes is involved in upregulation of gene expression upon pathogen infection [80,81].

To conclude, CHG LMRs are to a large extent tissue-specific, enriched in distal regions and TEs, but unlike LMRs in mammals, highly associated with H3K9me2 and H3K27me2, and not significantly overlapping with maize candidate enhancers [4,5], suggesting that, despite DNA hypomethylation, CHG LMRs may be located in heterochromatic regions [57,84] (46, 85). A potential function of the largely tissue-specific CHG LMRs is, however, unknown. It may in part be regions of relatively recently inserted TEs that are waiting to be stably inactivated through DNA hypermethylation.

Methylation of the external cytosine in both CCG and CWG sites is catalysed by CMT3 [8,12], whereby the binding of CMT3 to H3K9 methylated histone tails stimulates CMT3 to methylate CHG [85]. As reported previously, and also observed in this study, the methylation levels of the outer cytosine in the CCG context (mCCG) are lower than those in the CWG context [38] (Fig. 9; Additional file 4: Table S7). It has been shown that methylation of the internal cytosine (CmCG), is required for that of the external cytosine (mCCG; [38,68], and that the lower DNA methylation level of CCG sites is unlikely due to specific DNA demethylation of these sites [38]. The H3K9 methyltransferases SUVH4, and especially SUVH5 and SUVH6, are indicated to be redundantly required for mCCG [38]. Gouil and Baulcombe postulated that the external cytosine in CCG sites is methylated with a lower affinity than those in CWG sites [38]. Lower H3K9me2 levels at CCG sites than CWG sites could contribute to a lower efficiency of CMT3 to methylate the external cytosine in CCG sites. In line with this, CHG LMRs Data on the binding preferences of the H3K9 methyltransferases SUVH4, SUVH5 and SUVH6 support such hypothesis [66].

Gouil and Baulcombe [38] postulated that the binding of SUVH5/6 to methylated CWG sites would result in H3K9me2, recruiting CMT3, and methylation of not only the CWG sites, but also neighboring CCG sites. If that would be true, one would expect that the more methylated CWGs present in a region, the higher the methylation level of CCG sites. We did indeed observe a positive correlation between the methylation level of CWG regions and that of CCG regions. We did not observe a significant difference in the CWG concentration between CCG UMRs, LMRs and FMRs (Additional file 2: Figure S16), suggesting that the concentration of CWG as such does not affect the CCG methylation level. Future experiments will have to further unravel the mechanisms underlying the DNA methylation difference between CCG and CWG sites.

## Conclusions

To our knowledge, this is the first study in plants classifying DNA methylated regions into UMRs, LMRs and FMRs, and comparing their features with those in mammals. Unlike mammalian LMRs, plant LMRs are abundant in the CHG, but not CG context, and they do not overlap with candidate enhancers, but are enriched for heterochromatic marks, in line with having a yet unknown function.

We provide evidence that plant UMRs have a lot of features in common with mammalian UMRs, except that plant UMRs contain not only proximal regulatory regions like mammalian UMRs, but also distal regulatory sequences. Like mammalian UMRs, plant UMRs are mostly shared between tissues, supporting a model in which plant regulatory sequences are unmethylated in a tissue-independent manner, independent of their regulatory activity. We propose that for plants with large, highly methylated genomes, the genome-wide identification of UMRs is a useful approach to find distal *cis*-regulatory sequences. MethylSeekR is an excellent tool to find these UMRs.

## Methods

### Genome data

The reference genome sequences and associated annotations (TAIR10 [86]) for *Arabidopsis thaliana*, SL2.5 [87] for *Solanum lycopersicum*, RGAP7 [88] for *Oryza sativa Japonica*, AGPv4 [89] for *Zea mays* and RefBeet 1.2.2 [90] for *Beta vulgaris* were downloaded from Plants Ensembl [91]. TE annotations were obtained from Gramene [92] for *Zea mays*.

Raw data of genome-wide bisulphite sequencing experiments on Arabidopsis Columbia wild-type Aerial rosette tissue at 21/22 DAS (days after sowing) (GSE99404 [35]), wild-type tomato leaf and green fruit (SRX151217, SRX098309 [2] and SRX2008738 [38]), wild-type rice leaf tissue (GSM946552 [39]), wildtype B73 coleoptile shoot tissue (harvested 5 days after the start of germination) (GSE39232 [40]) and beet leaf (GSM2096947 [42]) were obtained from the NCBI database. FastX toolkit [93] was used to filter artefacts introduced by library construction such as linker and/or adaptor sequences, and to filter reads of which the qualities of more than 80 % of the bases were lower than a threshold of Q20. The reads were trimmed based on their per base sequence qualities and reads shorter than 70 bases after trimming were removed using PRINSEQ [94]. The read mapping to the reference genomes and methylation base calling was performed using BS-seeker2 [95,96].

Genome-wide bisulphite sequencing data on Arabidopsis columbia wild-type immature inflorescence (GSM1848703 [37]) and leaf (GSM207842 [36]) was obtained from the NCBI database.

After extracting i) the number of total reads covering a position, and ii) the number of reads without conversion at a given cytosine position in CpG context, the presence of partially methylated domains (PMDs; ref) was investigated. In mammals, PMDs have been observed in a small number of cell types and such domains need to be identified and masked prior to the segmentation of the genome into UMRs, LMRs and FMRs. To do so, the distribution of α-values for chromosome one and two of the four plant species was calculated, for CG and CHG methylation (Additional file 2: Figure S17). In most species and tissues examined, CG and CHG methylation clearly displayed a unimodal distribution, in all of them the vast majority of the α-values were below 1, indicating the absence of PMDs. This was also illustrated by plots (generated using the plotPMDSegmentation function) of randomly chosen regions of the maize genome (Additional file 2: Figure S18). Given there were no indications of PMDs, for subsequent analysis, no separate identification and masking of PMDs was performed.

The low and un-methylated regions were identified and median percent methylation against log2 number of CG in segments was plotted for CG, CHG and CHH data using MethylSeekR [16] with parameters m=0.5 and n=4 using public BSgenome packages for Arabidopsis [97] and rice [98]. For the other species, the BSgenome packages were compiled manually using the BSgenome packge [99] in R [56]. All the genomic regions that were not defined as UMRs nor LMRs, were defined as FMRs.

For maize H3K9me2 and H3K27me2 [57] data, the quality control was performed in the same manner as for BS-seq reads described above. After quality control of the reads, the reads were mapped to AGPv4 using BWA [100]. The signal intensity data from MACS2 [101] peak calling output was converted to bigWig file using the wigToBigWig tool [102]. The H3K9ac and DNase-seq data generation and analysis were described in Oka et al [4]. H3K9me2 and H3K27me2 data is derived from inner stem tissue of 1-month-old plants approximately in the V4 stage with roots and exposed leaf blades removed.

Genomic regions were defined as follows: genic regions, exons and transposable elements (TEs) were annotated according to the reference annotation. The annotated exons include the untranslated regions (UTRs). The entire genome, except for the genic regions, were called intergenic regions. Introns were genic regions excluding exons. Promoters were defined as the sequence 1 kb upstream and 200 bp downstream of transcription start sites (TSSs). Flanking regions were defined as sequences 4 kb upstream from promoter regions and 5 kb downstream from the transcription termination sites (TTSs; Fig. 5a). Distal regions were intergenic regions that were not classified above. The parts in each genomic region were identified using BEDtools [45]; total lengths in Mb for the genomic composition, in kb for UMRs and LMRs were calculated and plotted using plotrix package [103] in R [56]. The fraction was calculated by the UMR or LMR occupancy in kb divided by the total lengths of each genomic region in kb.

The distances between any regions and TSSs was calculated from TSS to the closest end of regions. When the TSS locates within a region, the reported distance was 0. The tissue specificities of UMRs and LMRs were measured by the overlap of CHG UMRs and CHG LMRs between tissues. To observe the UMR, LMR or FMR distribution trends over chromosome 1, the chromosome 1 from all the selected species were binned for calculating the window coverage to see the trend over the chromosome better. The average signal intensities of chromatin marks were extracted for each UMRs or LMRs using bwtools [104]. Their median and standard error were calculated using R [56]. The enrichment of CHG, CCG and CWG LMRs in TEs were calculated by taking the sum of the CHG LMR overlapping TE length divided by the sum of the TE lengths for each TE family. The levels of methylation at the first and the second C in CCG context were extracted using bwtools [104] and plotted in heatmaps using ggplot2 package [105] in R [56]. The distance calculation, overlap measurement and coverage calculation was performed using BEDtools [45].

### Additional files

**Additional file 1: Figure S1.** Number of CpGs against median CG methylation levels for human genomewide bisulfite data sets re-printed from Burger et al. (2013). **Figure S2.** Number of CpGs against median methylation levels for CG, CHG, CHH, CCG and CWG methylation context in Arabidopsis (genome-wide), tomato, rice and maize (chromosome 1) bisulfite data sets. **Figure S3.** Number of CpGs against median methylation levels for CG, CHG, CHH, CCG and CWG methylation context in tomato, rice and maize bisulfite data sets for all chromosomes except chromosome 1. **Figure S4.** Distributions of median percent methylation of U/L/FMRs for CG and CHG context. **Figure S5.** Number of CpGs against median methylation of UMRs, LMRs and FMRs in CG and CHG contexts for Arabidopsis leaf, tomato leaf, rice leaf and maize coleoptile. **Figure S6.** Number of CpGs against median methylation of UMRs and LMRs in CG and CHG contexts for low coverage data in beet and tomato leaf. (PDF 32.6 Mb)

**Additional file 2: Figure S7.** Coverage of CG/CHG UMRs and LMRs over 0.1 and 1 Mb windows in chromosome 1 for Tomato leaf, green fruit and Maize coleoptile. **Figure S8.** Coverage of FMRs in CG and CHG context over 0.1 Mb and 1 Mb windows in chromosome 1 for Arabidopsis seedling, leaf and immature florescence, tomato leaf and green fruit, rice leaf and maize coleoptile. **Figure S9.** Genomic composition and distributions of UMRs and LMRs over defined genomic regions of Arabidopsis, tomato, rice and maize. **Figure S10.** The fractions CG and CHG un- and low methylated regions occupy at the various genomic regions. **Figure S11.** Distribution of distances between UMRs or LMRs and their closest TSSs in log2 scale. **Figure S12.** Sizes of UMRs and LMRs in plants and mammals in base pairs. **Figure S13.** Fraction of overlapping CG and CHG UMRs and LMRs between tomato leaf and green fruit. **Figure S14.** Enrichment of chromatin marks in FMRs, LMRs and UMRs in the maize genome. **Figure S15.** Coverage of UMRs, LMRs or FMRs in CCG and CWG context over 0.1 and 1 Mb windows in chromosome 1 for Arabidopsis leaf, tomato leaf, rice leaf and Maize coleoptile. **Figure S16.** Concentration of CWGs in every CCG UMR, LMR or FMR in maize per kb. **Figure S17.** The distribution of α-values for chromosome 1 and 2 of the indicated species and tissues, for CG and CHG. **Figure S18.** DNA methylation levels at randomly chosen regions of the maize genome are plotted using the plotPMDSegmentation function. (PDF 1.9 Mb)

**Additional file 3: Tables S1.** Summary of datasets used in this study. **Table S2.** Number of CG LMRs and UMRs in different human cell types and mouse ES cells. **Table S3.** Median and maximum sizes of LMRs and UMRs in bp. **Table S4.** Overlap between CHG UMRs/LMRs and DHSs, H3K9ac enriched regions, and enhancer candidates. **Table S5.** Overlap between CG UMRs/LMRs and DHSs, H3K9ac enriched regions, and enhancer candidates. **Table S6.** Number of CHG/CG UMRs/LMRs that overlap with ACRs and vice versa (DOCX 127 kb). **Table S7.** Overlap between CCG and CWG UMRs, LMRs and FMRs. (docx 128 kb)

**Additional file 4: Table S7.** Counts used in the heatmaps of Figure 9. (XLSX 29 kb)

## Supporting information

additional tabels

## Declarations

### Ethics approval and consent to participate

Not Applicable

### Consent for publication

Not Applicable

### Availability of data and material

The raw and processed BS-seq datasets were obtained from the GEO and SRA databases (GSE99404, GSM1848703, GSM2078421, SRX151217, SRX098309, SRX2008738, GSM946552, GSE39232).

### Authors’ contributions

HCJH and MS designed the project. RO and MB conducted the computational data analysis and generated the Figures and Tables. All authors were involved in data interpretation and discussion. RO, HCJH and MS wrote the manuscript. All authors read and approved the final manuscript.

## Acknowledgements

We are grateful to Nathan M. Springer and Sarah N. Anderson for their assistance and advice regarding maize TE annotations, and for helpful discussions at the first phase of this study. We thank Blaise Weber, Mariliis Tark-Dame, Till Bey, Jihed Chouaref and Aimer Gutierrez Diaz for helpful discussions on the study, Jonathan Gent for valuable discussions and critical review of an earlier version of the manuscript, and Hans de Jong for advice on the tomato genome. We thank Roman Lakerveld for his help with the reference list.

