## additional figures 1 to 6 for "In plants distal regulatory sequences overlap with unmethylated rather than low-methylated regions, in contrast to mammals"

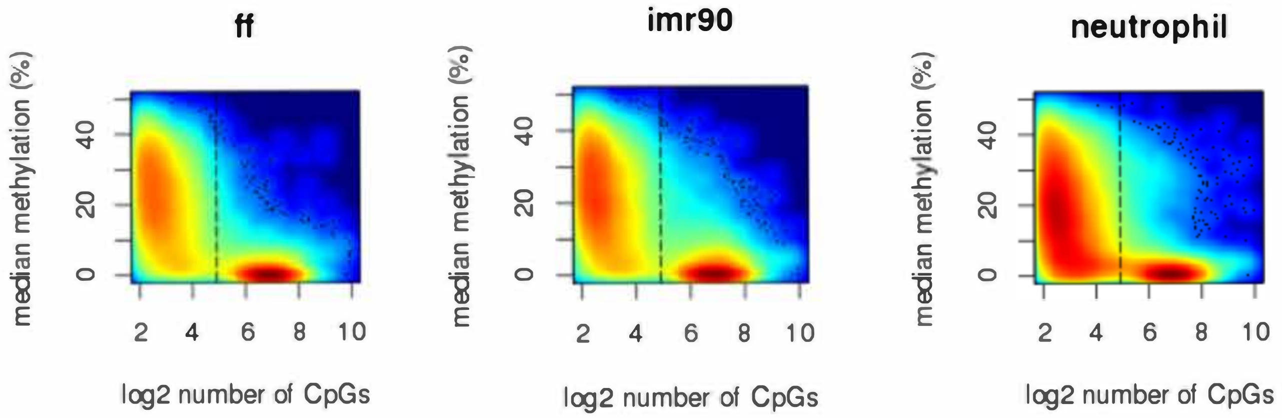


Supplementary Figure S1. Number of CpGs against median CG methylation levels for human genome-wide bisulfite data sets re-printed from Burger et al. (2013). The black dashed vertical lines indicate 30 CpGs, which were used to classify UMRs and LMRs in mammalian data sets using MethylSeekR (Burger et al 2013). Low to high density of data points is indicated by colour (blue to red, respectively); the colour scales are different for each dataset and not comparable between datasets.

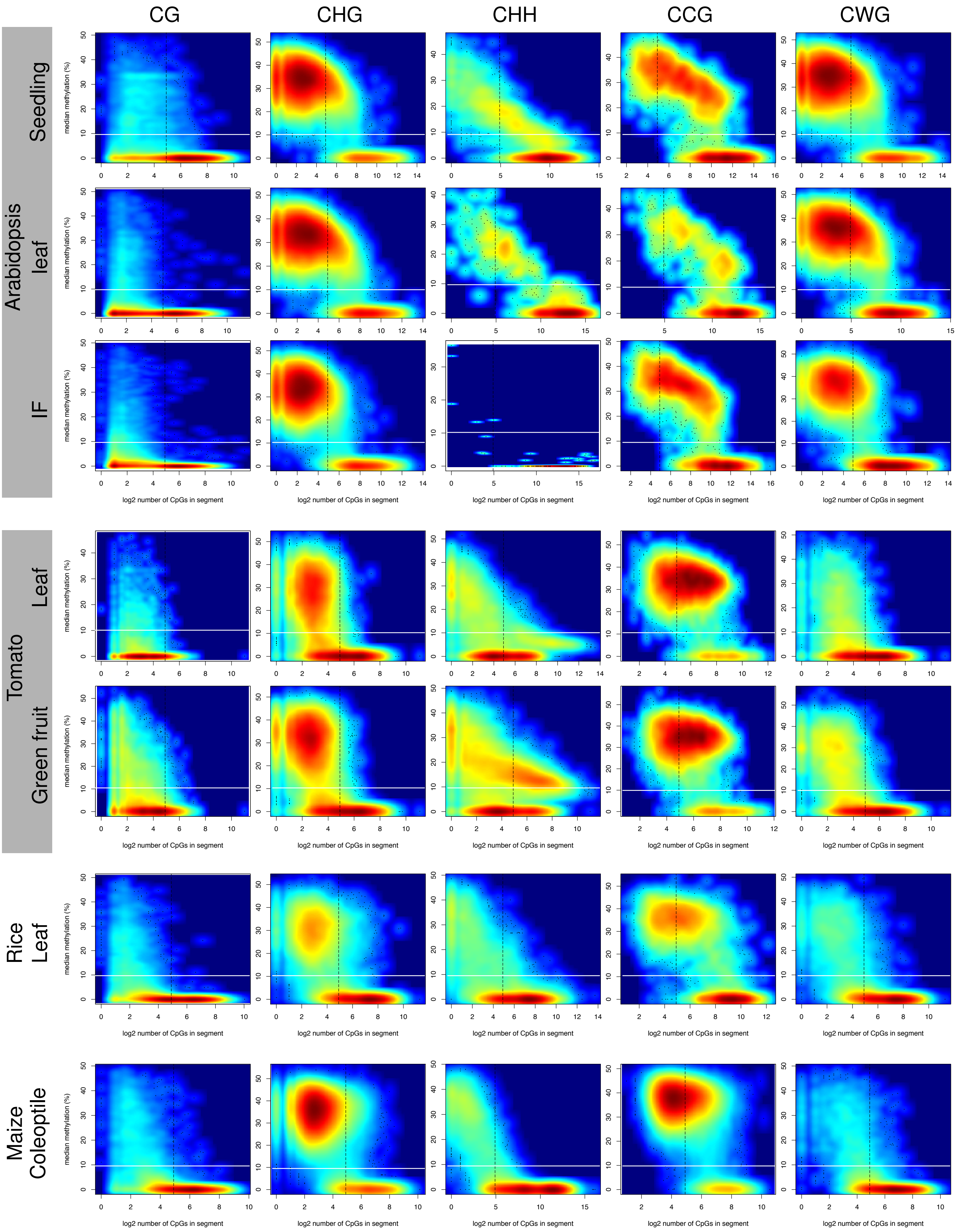

CG
CHG
CHH
CCG
CWG
Seedling
Arabidopsis leaf
IF
Tomato
Leaf
Green fruit
Rice Leaf
Maize Coleoptile
median methylation (%)
log2 number of CpGs in segment

Supplementary Figure S2. Number of CpGs against median methylation levels for CG, CHG, CHH, CCG and CWG methylation context in Arabidopsis (genome-wide), tomato, rice and maize (chromosome 1) bisulfite data sets. The black dashed vertical lines indicate 30 CpGs, which were used to classify UMRs and LMRs in mammalian data sets using MethylSeekR (Burger et al 2013). The white solid horizontal lines indicate the 10% median methylation boundary chosen in this study to distinguish UMRs and LMRs in plants. Low to high density of data points is indicated by colour (blue to red, respectively); the colour scales are different for each dataset and not comparable between datasets.

(A) Tomato leaf CG

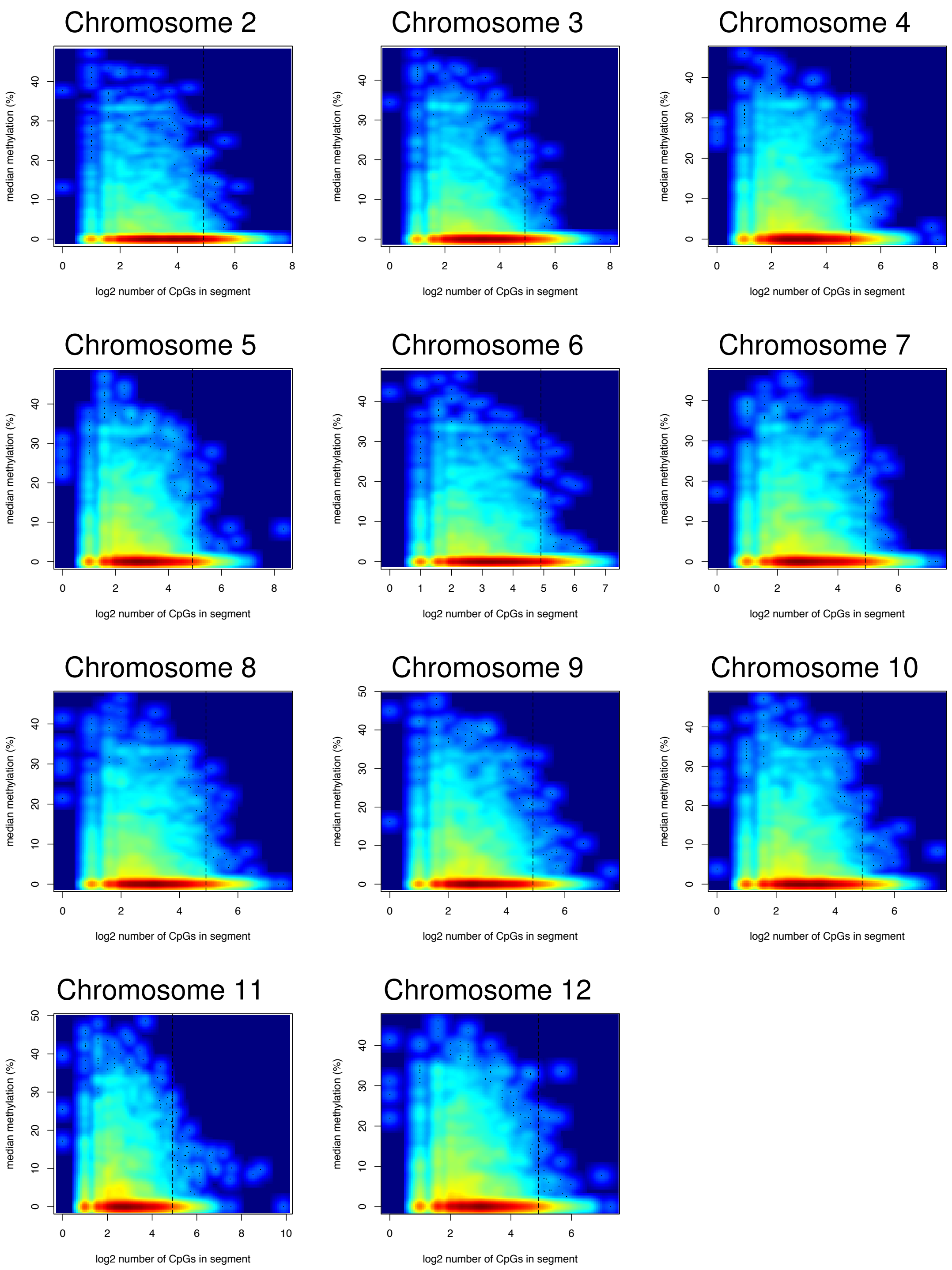


Supplementary Figure S3. Number of CpGs against median methylation levels for CG, CHG, CHH, CCG and CWG methylation context in tomato, rice and maize bisulfite data sets for all chromosomes except chromosome 1. (A) tomato leaf CG, (B) CHG, (C) CHH, (D) CCG, and (E) CWG; (F) tomato green fruit CG, (G) CHG, (H) CHH, (I) CCG, and (J) CWG; (K) rice leaf CG, (L) CHG, (M) CHH, (N) CCG, and (O) CWG; (P) maize coleoptile CG, (Q) CHG, (R) CHH, (S) CCG, and (T) CWG. The black dashed vertical lines indicate 30 CpGs, which were used to classify UMRs and LMRs in mammalian data sets using MethylSeekR (Burger et al 2013). Low to high density of data points is indicated by colour (blue to red, respectively); the colour scales are different for each dataset and not comparable between datasets.

(B) Tomato leaf CHG

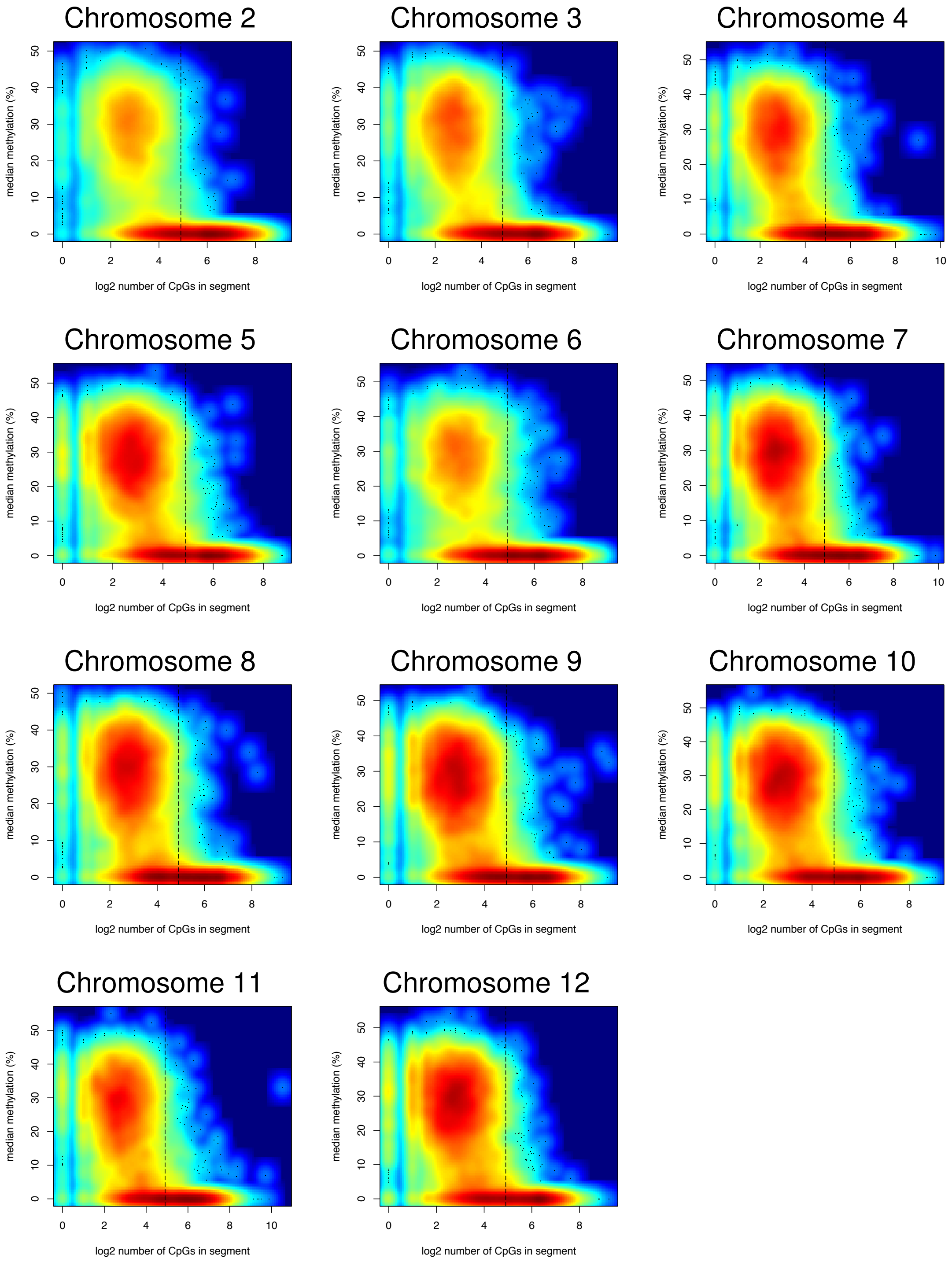

(C) Tomato leaf CHH

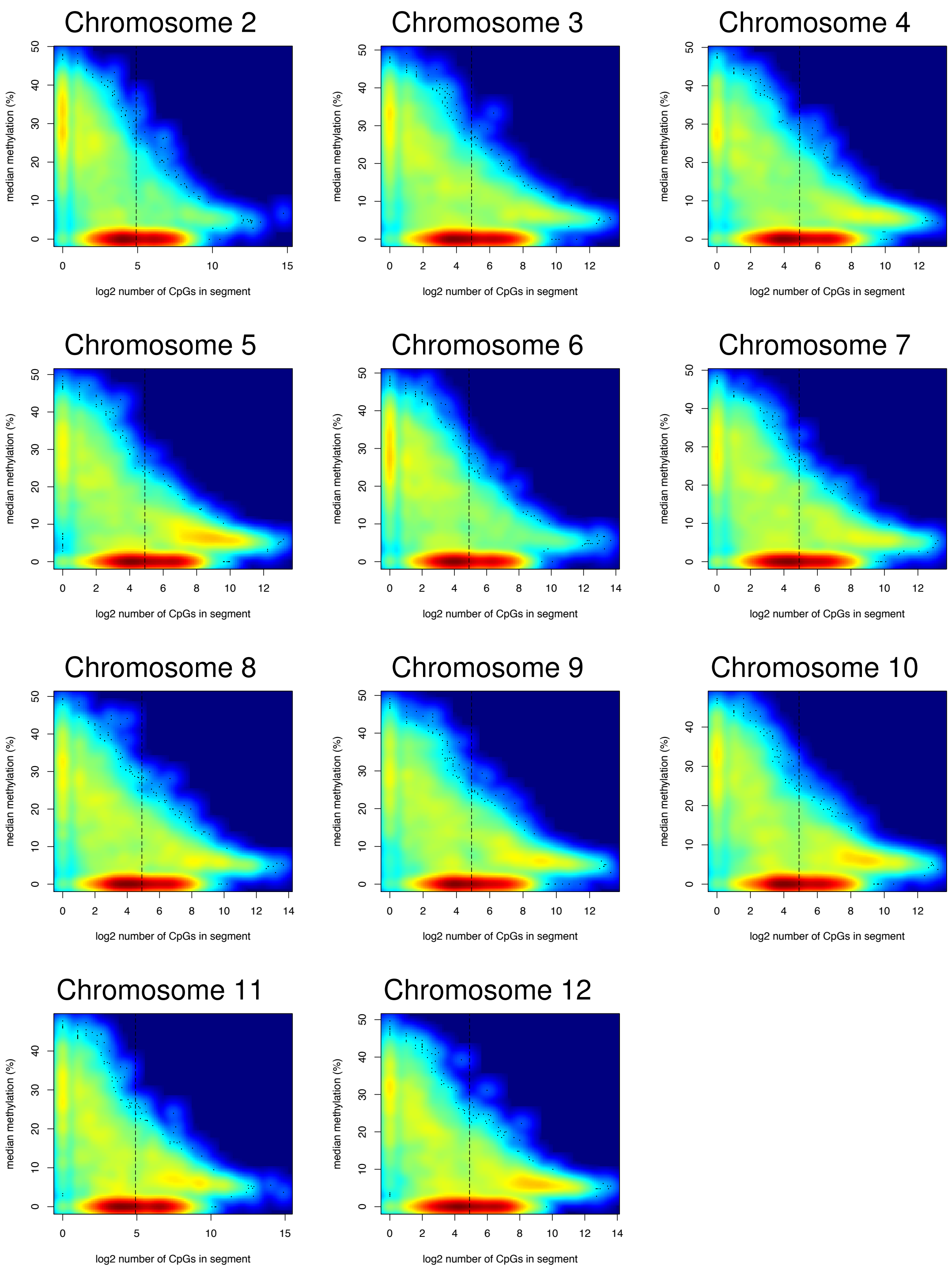

(D) Tomato leaf CCG

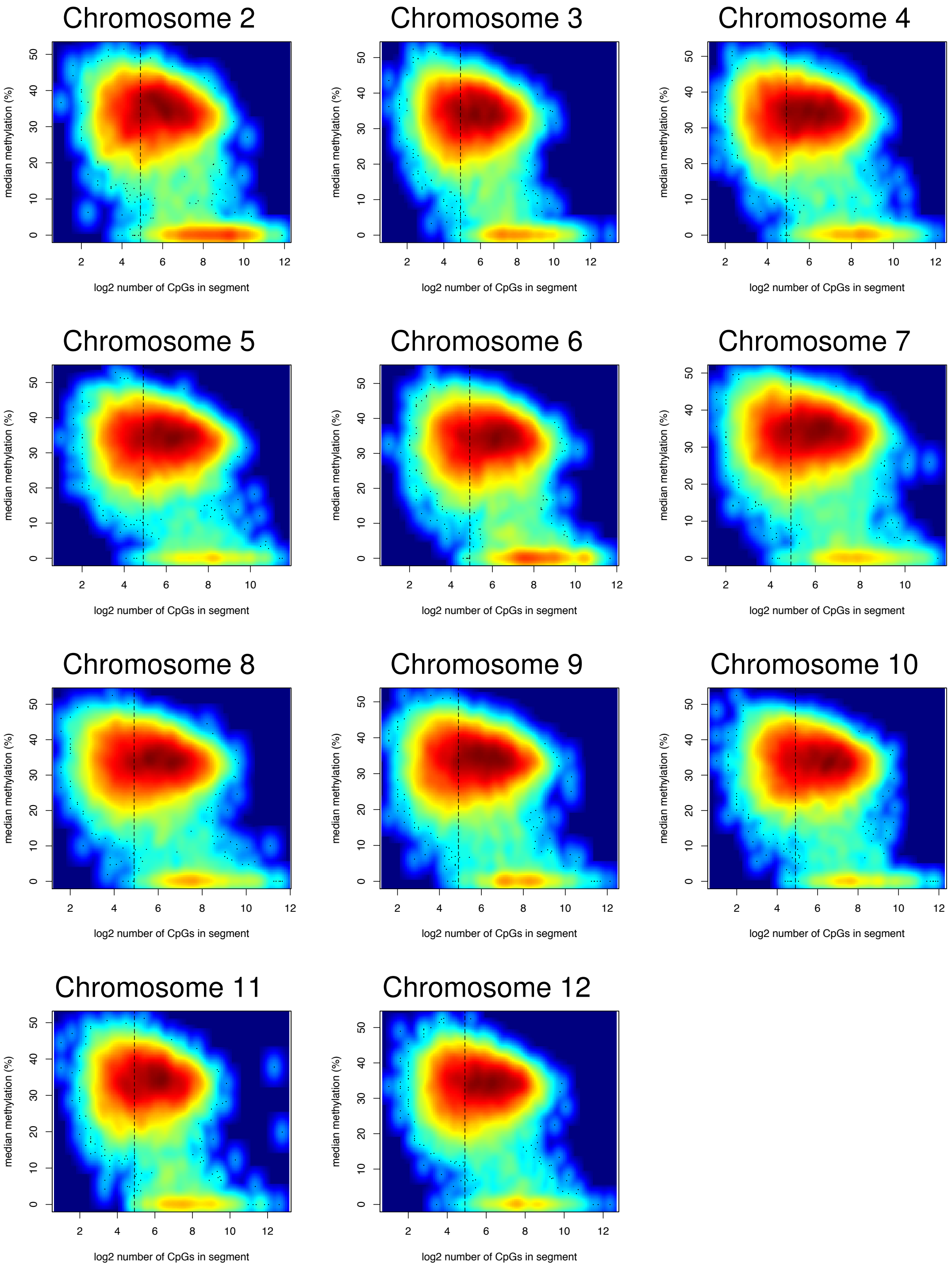

(E) Tomato leaf CWG

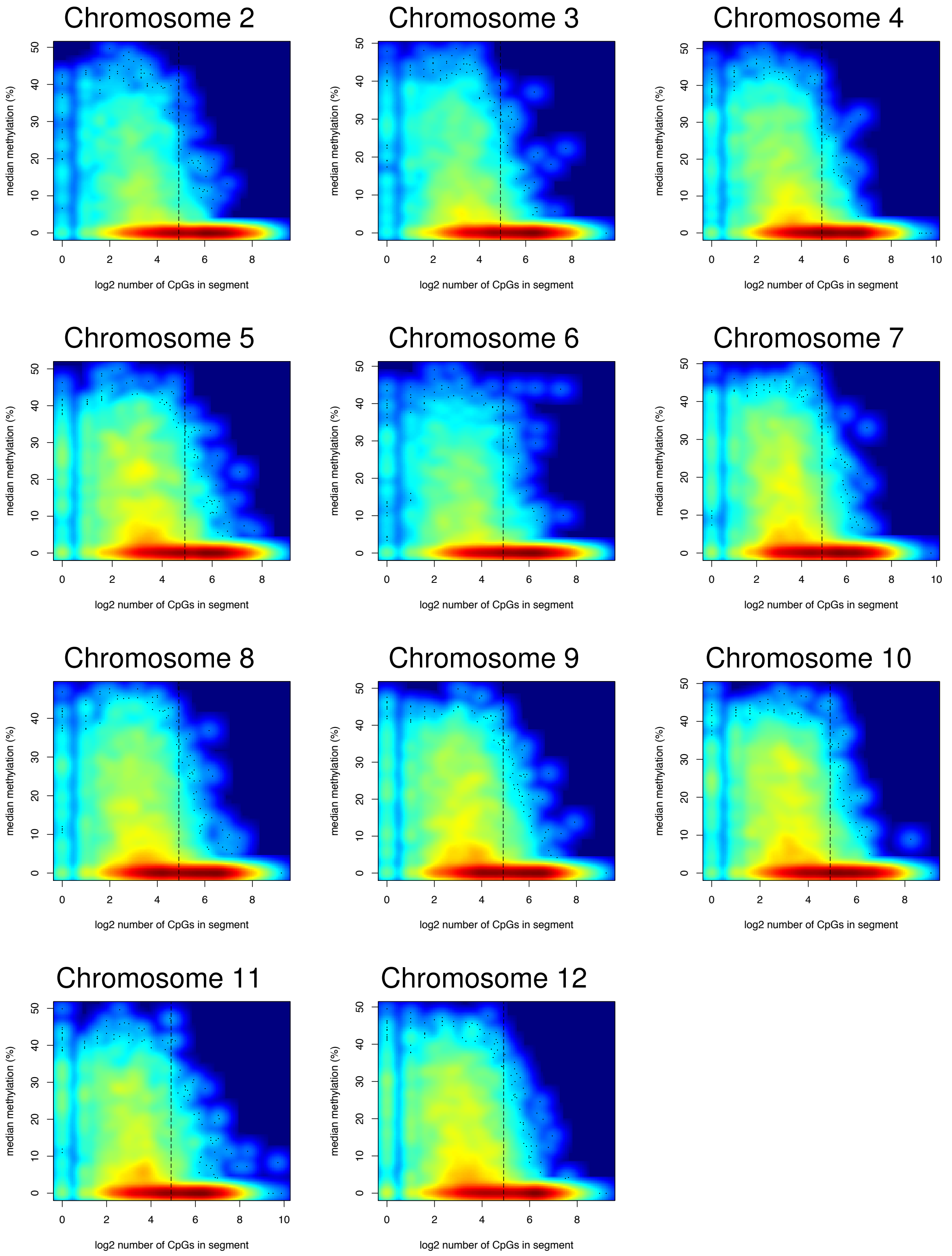

(F) Tomato green fruit CG

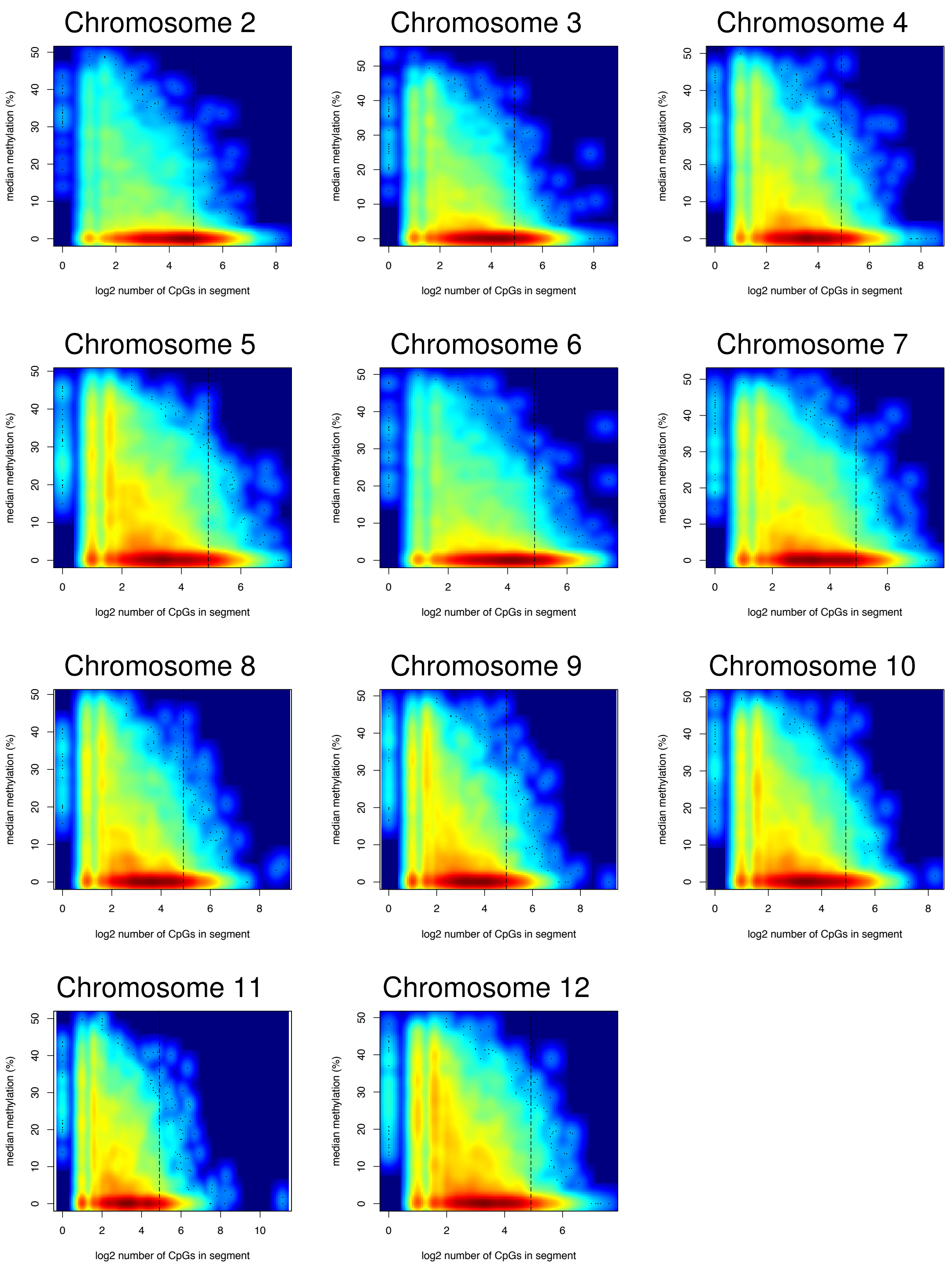

(G) Tomato green fruit CHG

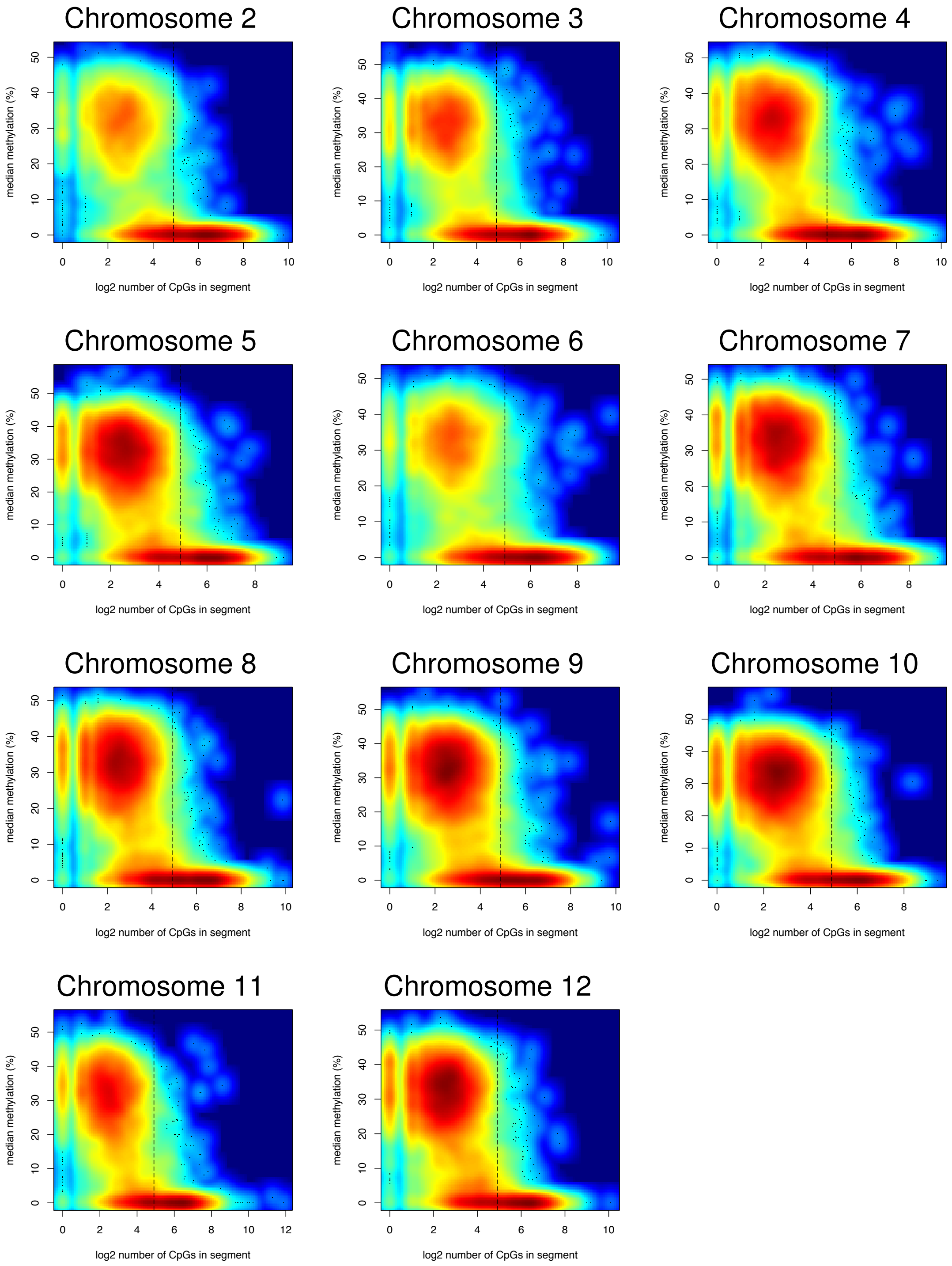

### (H) Tomato green fruit CHH

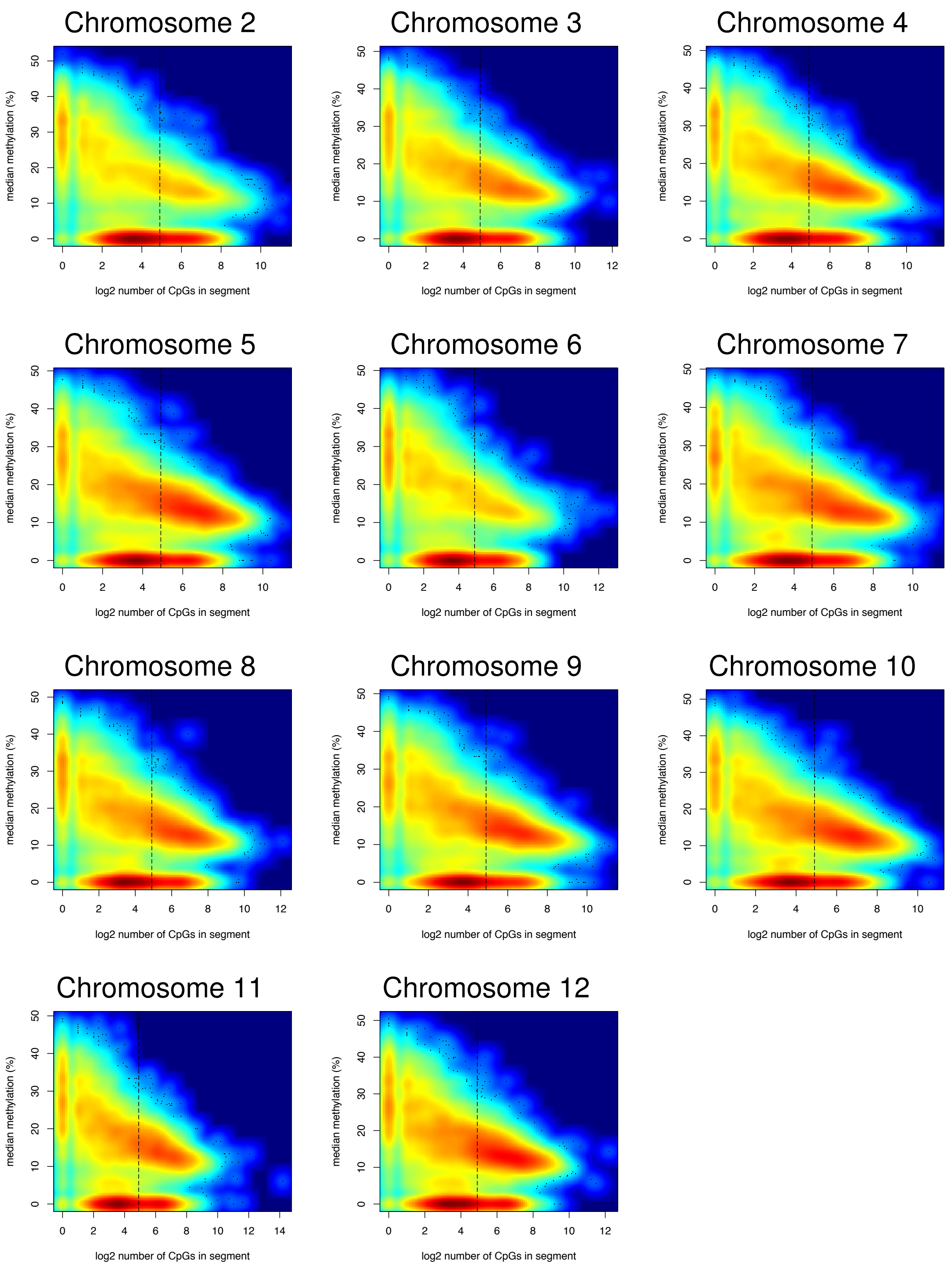

(I) Tomato green fruit CCG

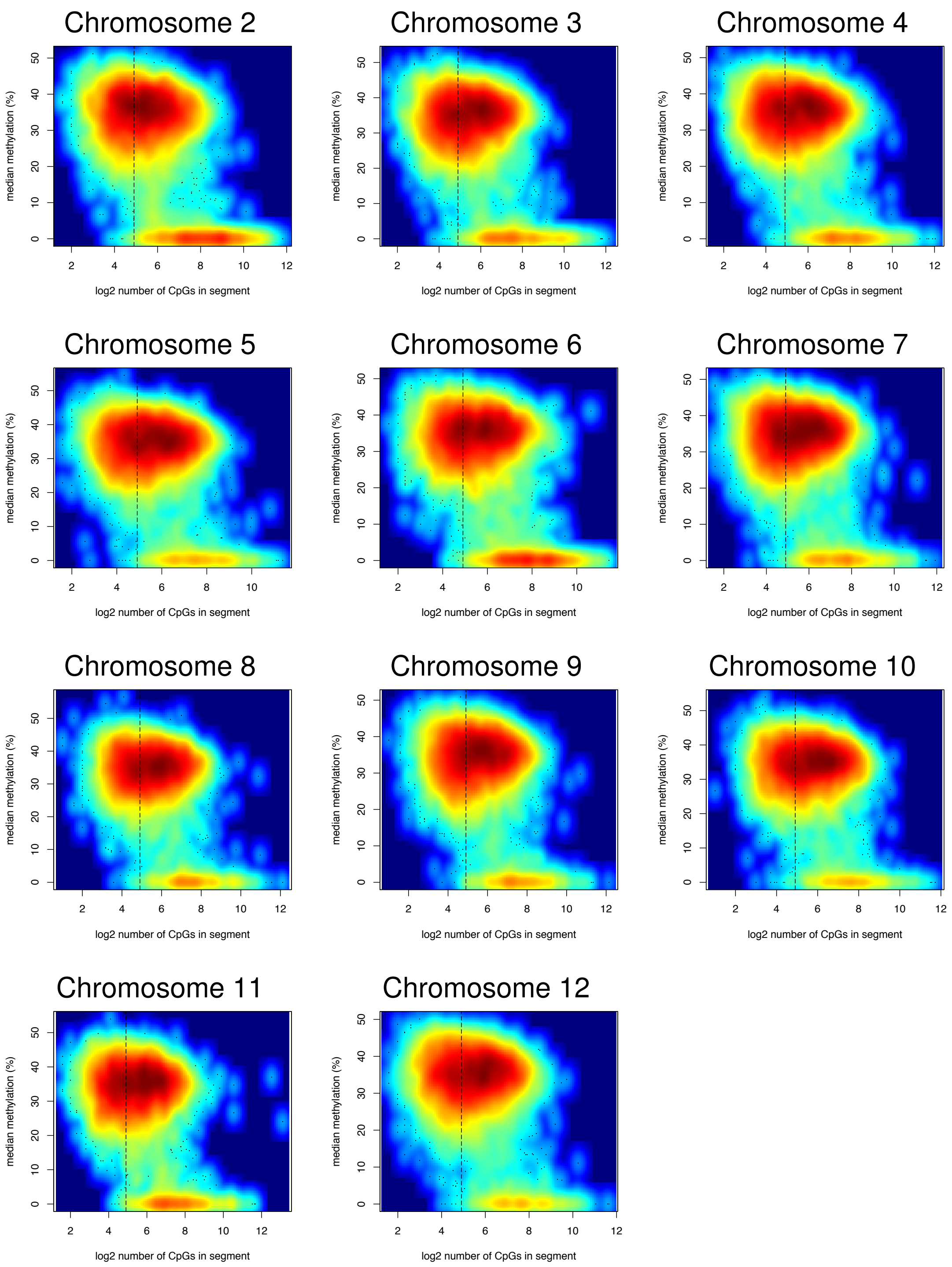

(J) Tomato green fruit CWG

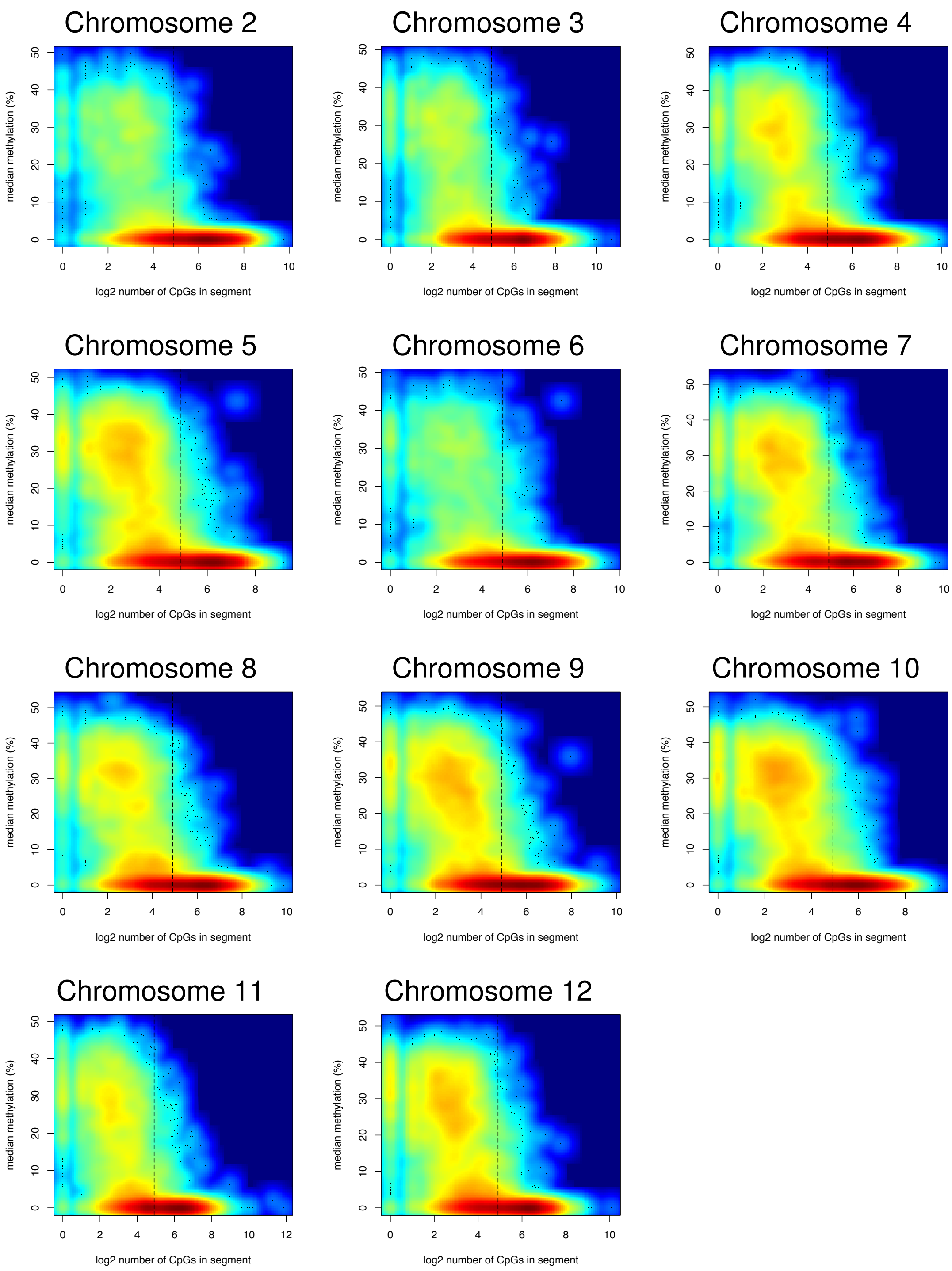

(K) Rice CG

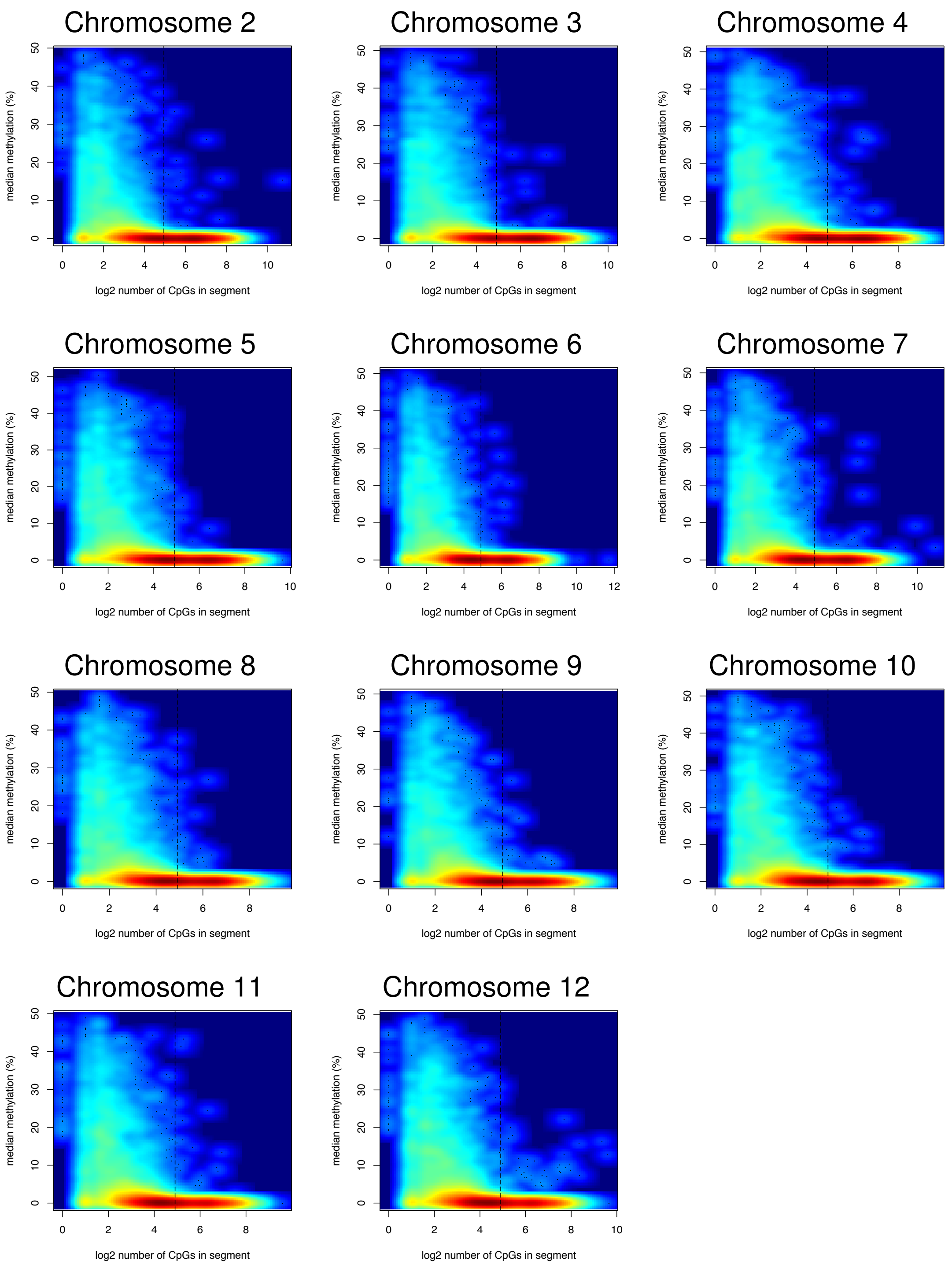
Chromosome 2
Chromosome 3
Chromosome 4
Chromosome 5
Chromosome 6
Chromosome 7
Chromosome 8
Chromosome 9
Chromosome 10
Chromosome 11
Chromosome 12
median methylation (%)
log2 number of CpGs in segment

(L) Rice CHG

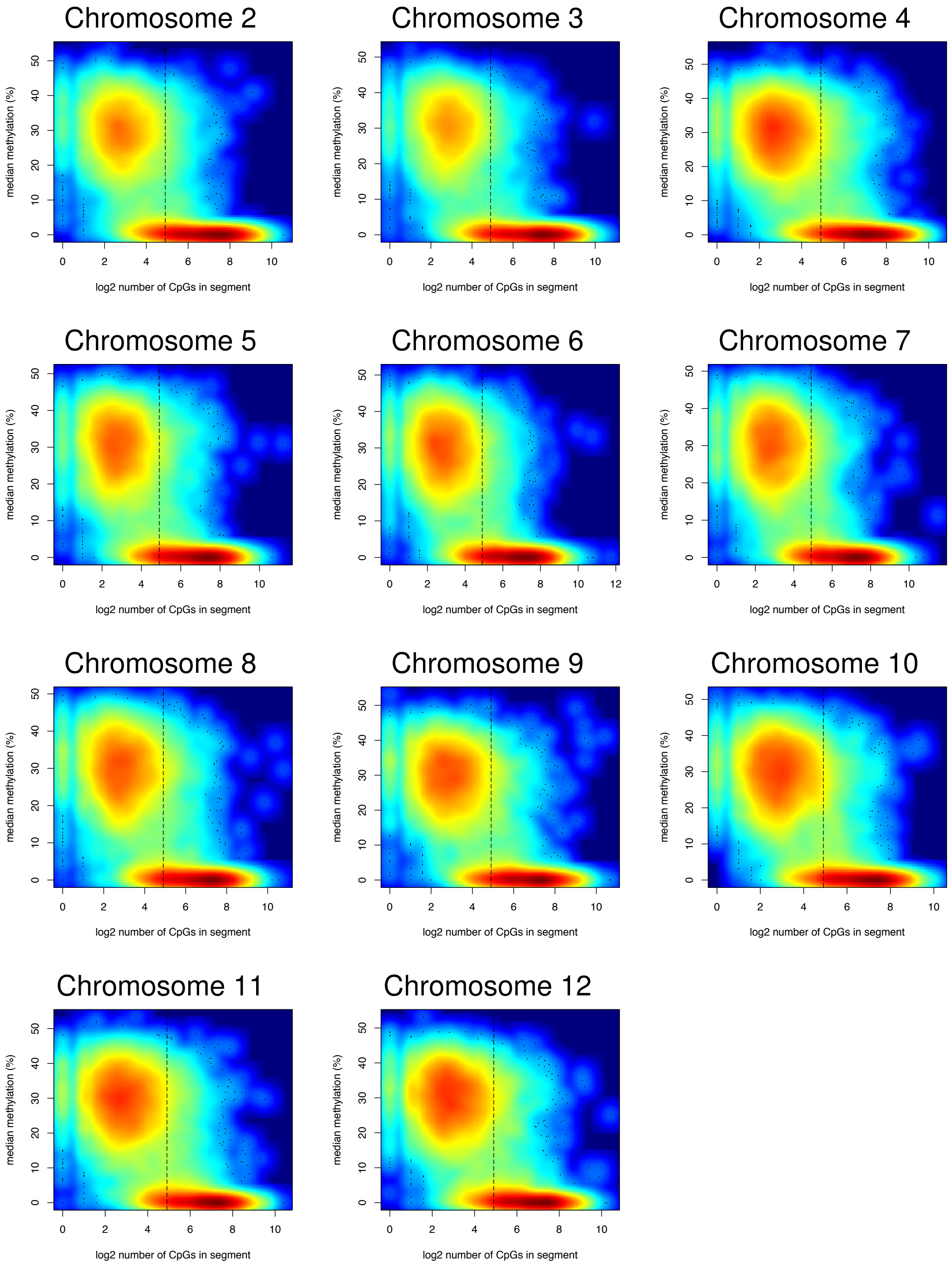
Chromosome 2
Chromosome 3
Chromosome 4
Chromosome 5
Chromosome 6
Chromosome 7
Chromosome 8
Chromosome 9
Chromosome 10
Chromosome 11
Chromosome 12
median methylation (%)
log2 number of CpGs in segment

(M) Rice CHH

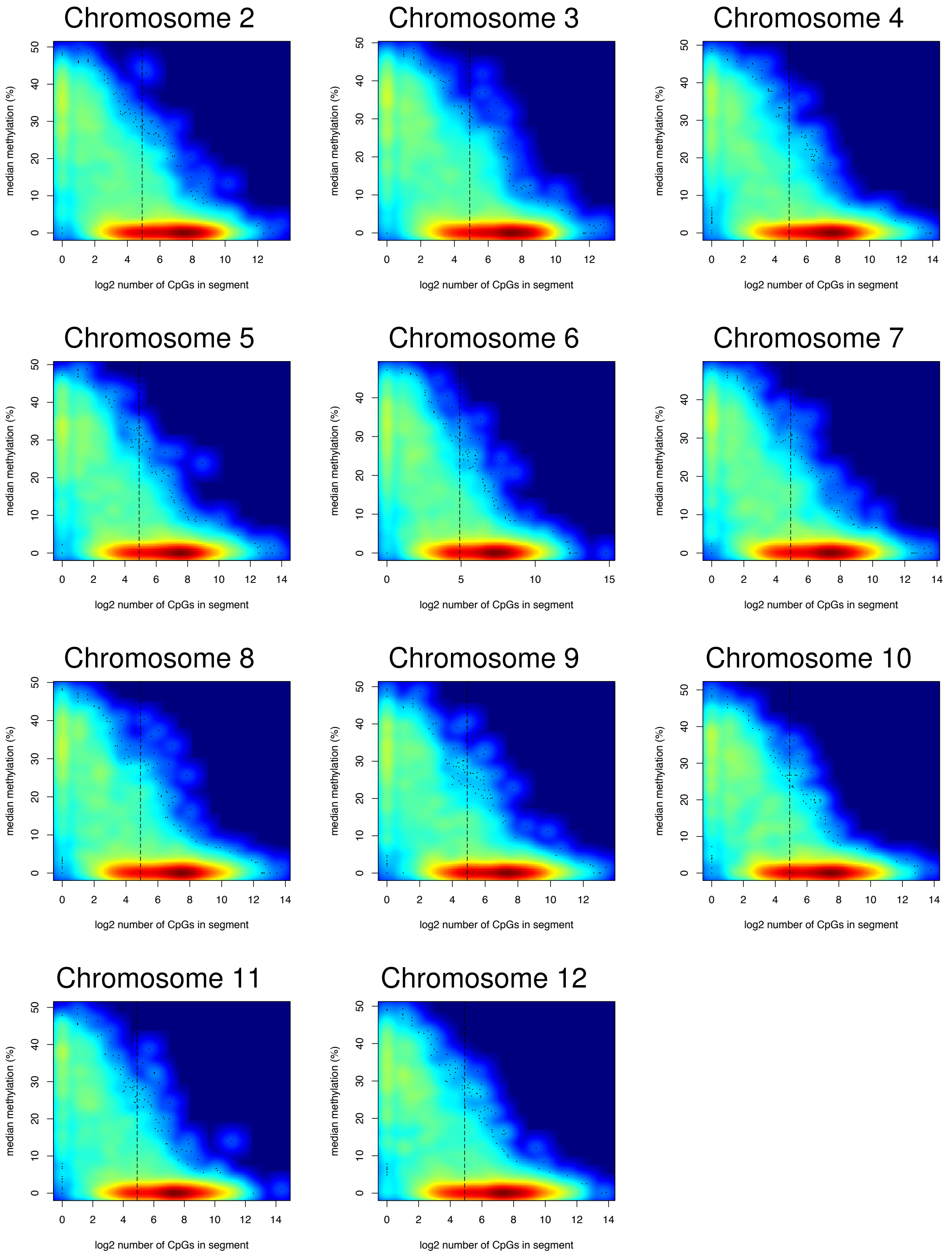
Chromosome 2
Chromosome 3
Chromosome 4
Chromosome 5
Chromosome 6
Chromosome 7
Chromosome 8
Chromosome 9
Chromosome 10
Chromosome 11
Chromosome 12
median methylation (%)
log2 number of CpGs in segment

(N) Rice CCG

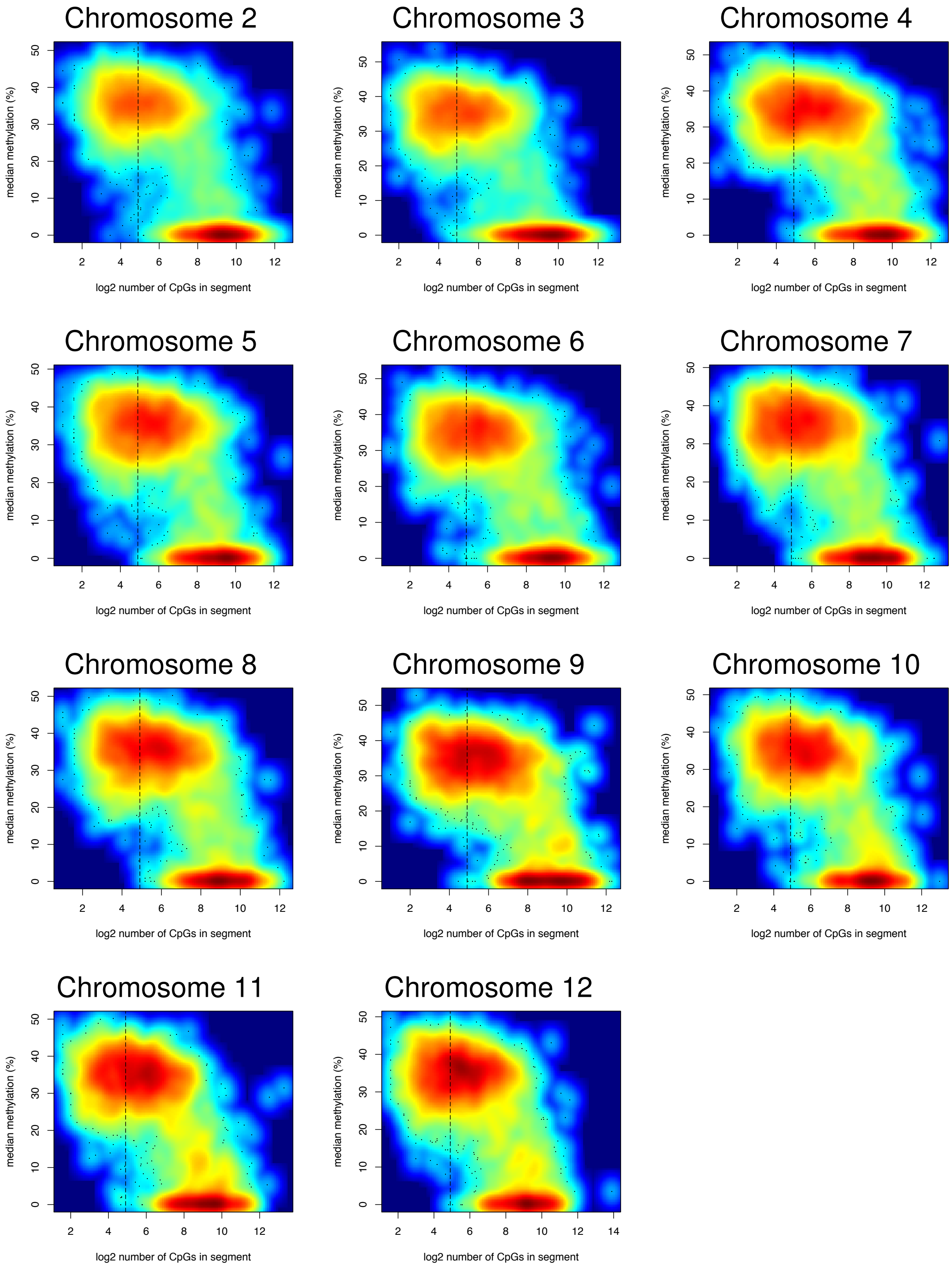
Chromosome 2
Chromosome 3
Chromosome 4
Chromosome 5
Chromosome 6
Chromosome 7
Chromosome 8
Chromosome 9
Chromosome 10
Chromosome 11
Chromosome 12
median methylation (%)
log2 number of CpGs in segment

#### (O) Rice CWG

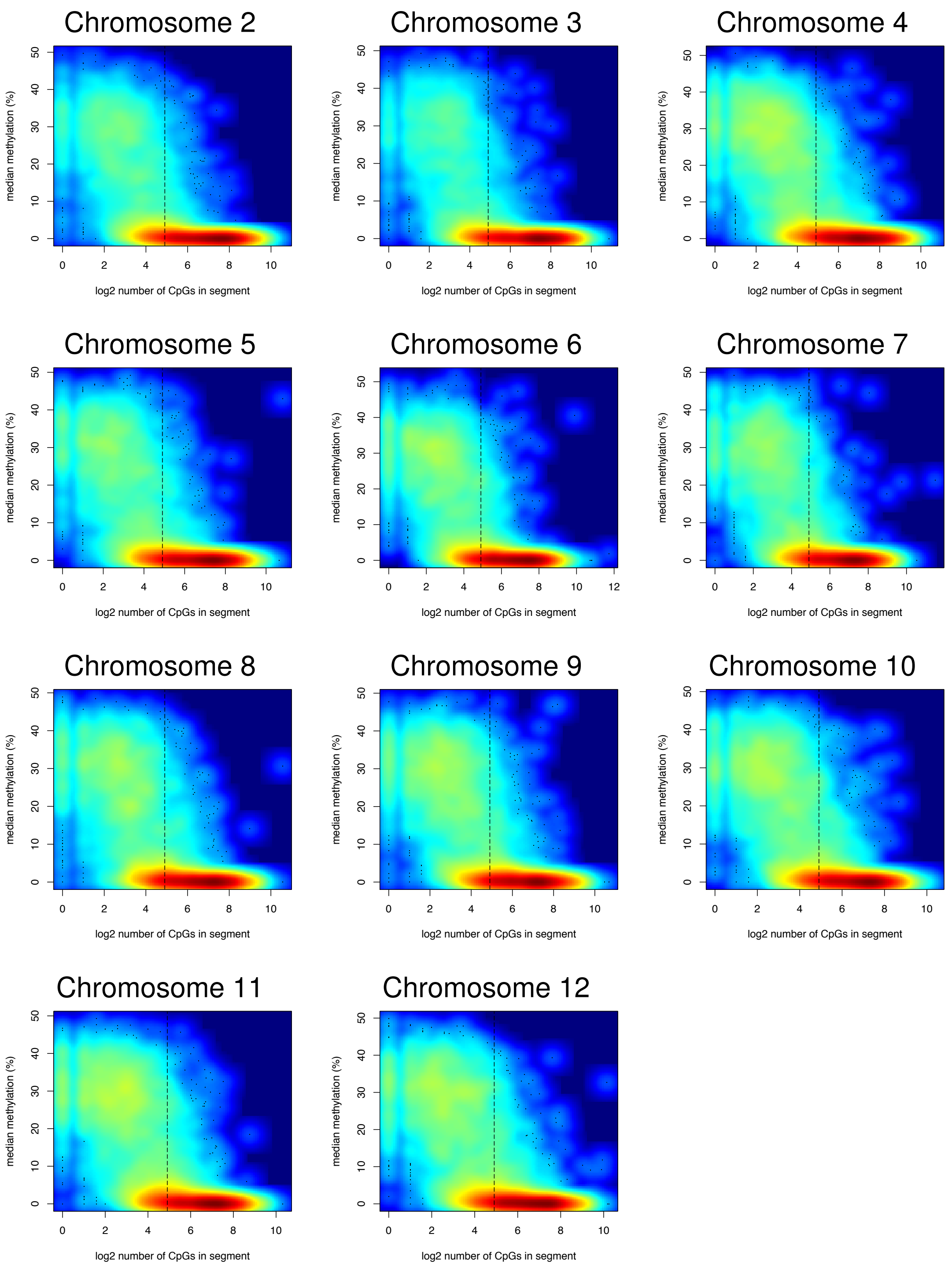

Chromosome 2
Chromosome 3
Chromosome 4
Chromosome 5
Chromosome 6
Chromosome 7
Chromosome 8
Chromosome 9
Chromosome 10
Chromosome 11
Chromosome 12
median methylation (%)
log2 number of CpGs in segment

(P) Maize CG

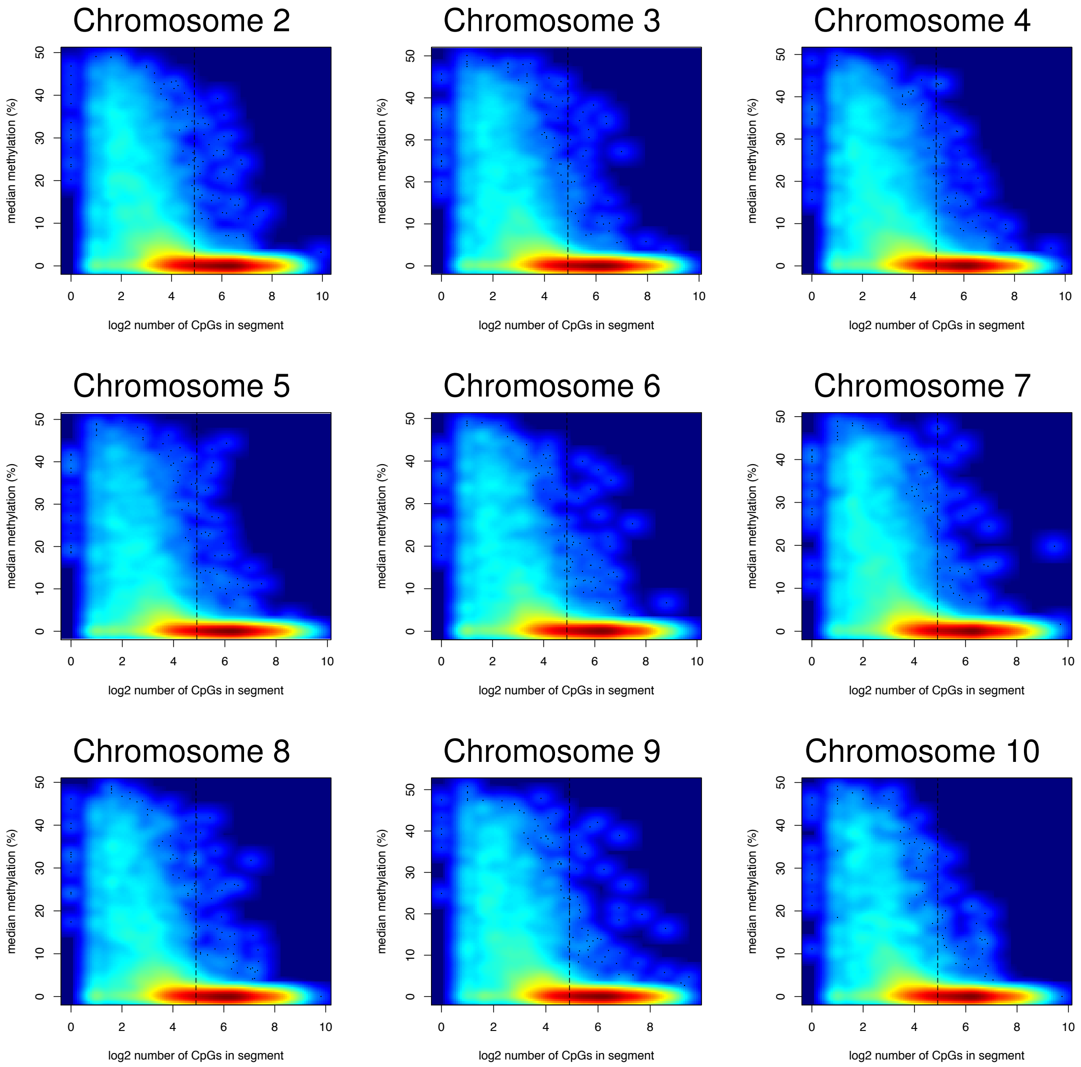

Chromosome 2
Chromosome 3
Chromosome 4
Chromosome 5
Chromosome 6
Chromosome 7
Chromosome 8
Chromosome 9
Chromosome 10
median methylation (%)
log2 number of CpGs in segment

(Q) Maize CHG

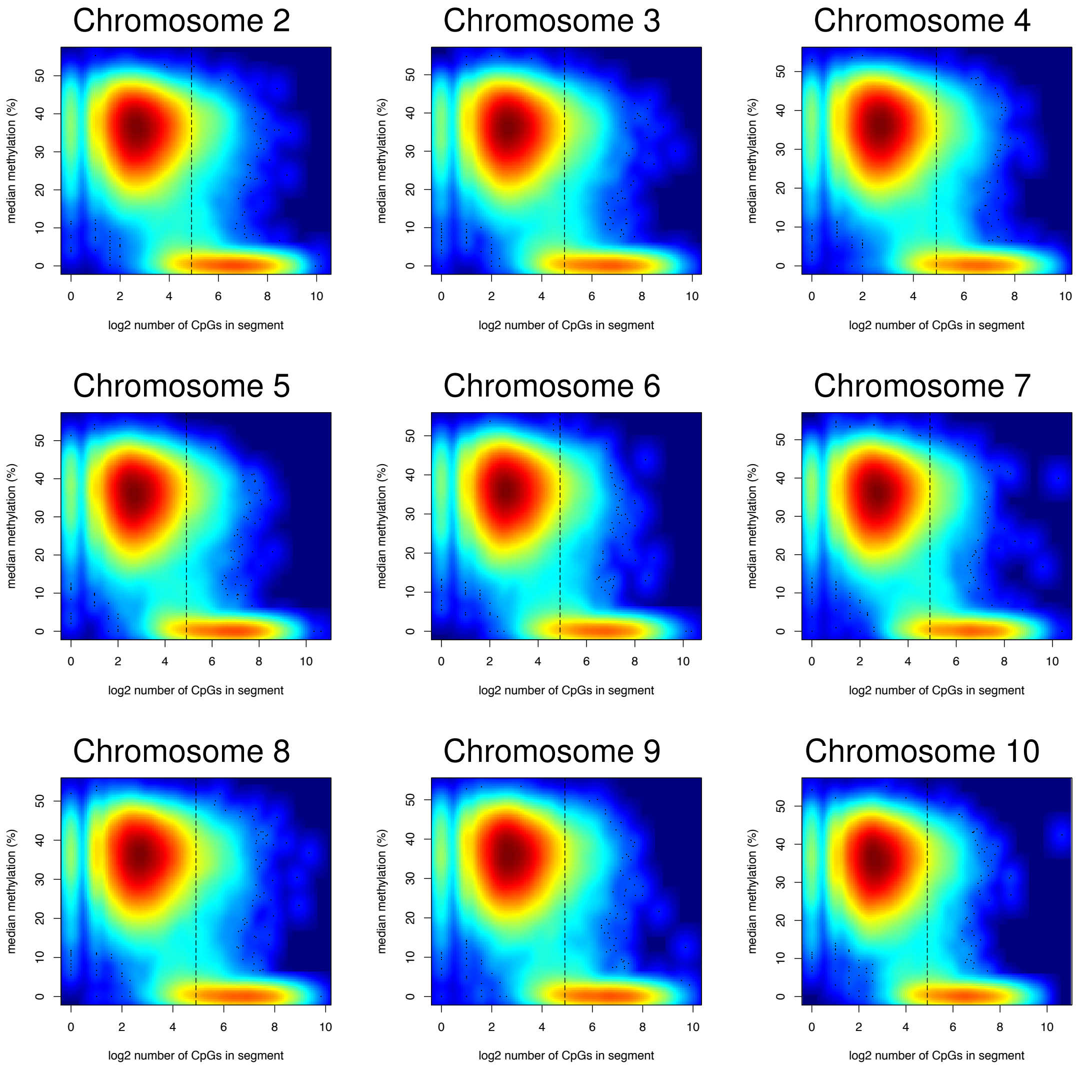

Chromosome 2
Chromosome 3
Chromosome 4
Chromosome 5
Chromosome 6
Chromosome 7
Chromosome 8
Chromosome 9
Chromosome 10
median methylation (%)
log2 number of CpGs in segment

(R) Maize CHH

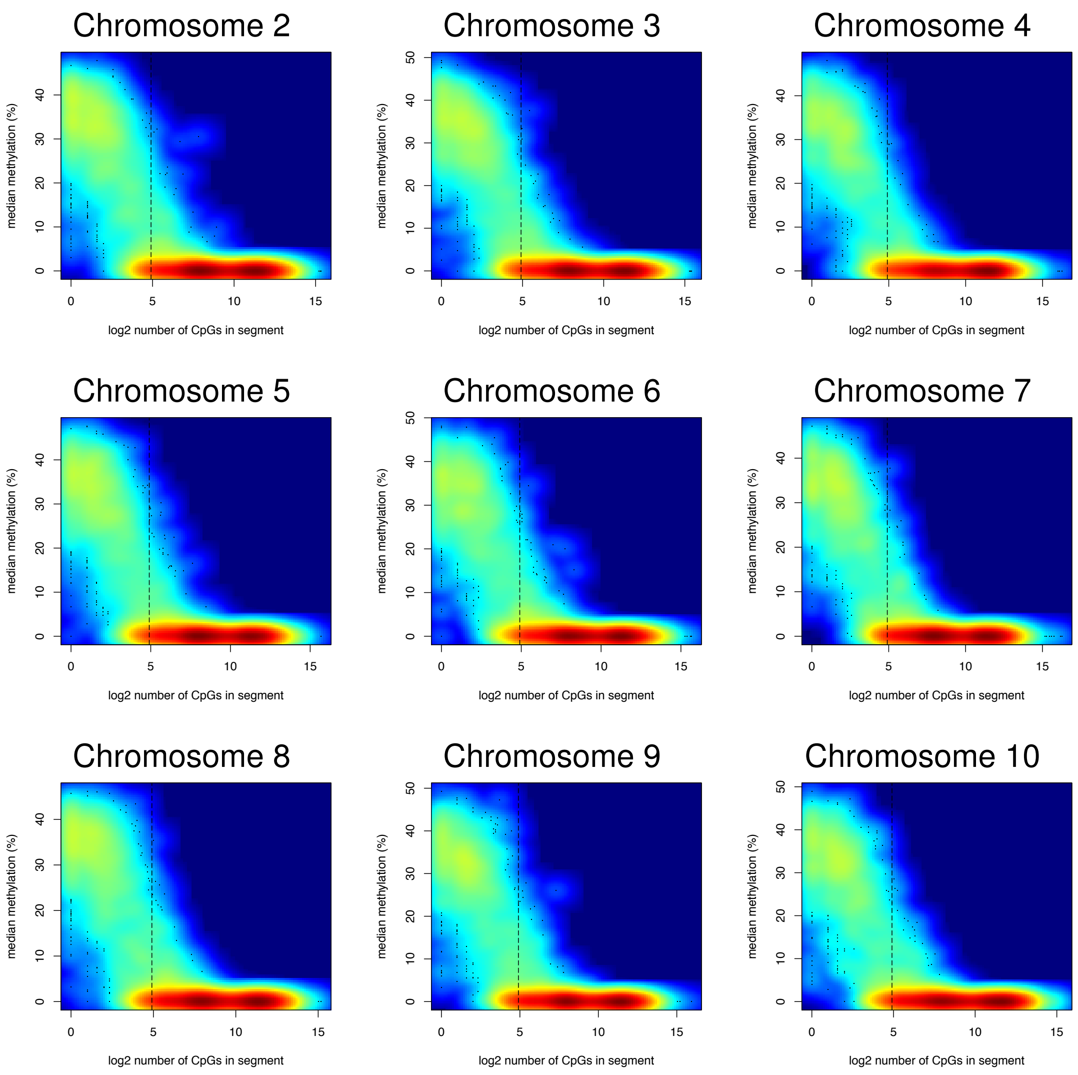

Chromosome 2
Chromosome 3
Chromosome 4
Chromosome 5
Chromosome 6
Chromosome 7
Chromosome 8
Chromosome 9
Chromosome 10
median methylation (%)
log2 number of CpGs in segment
0
5
10
15
20
30
40
50

(S) Maize CCG

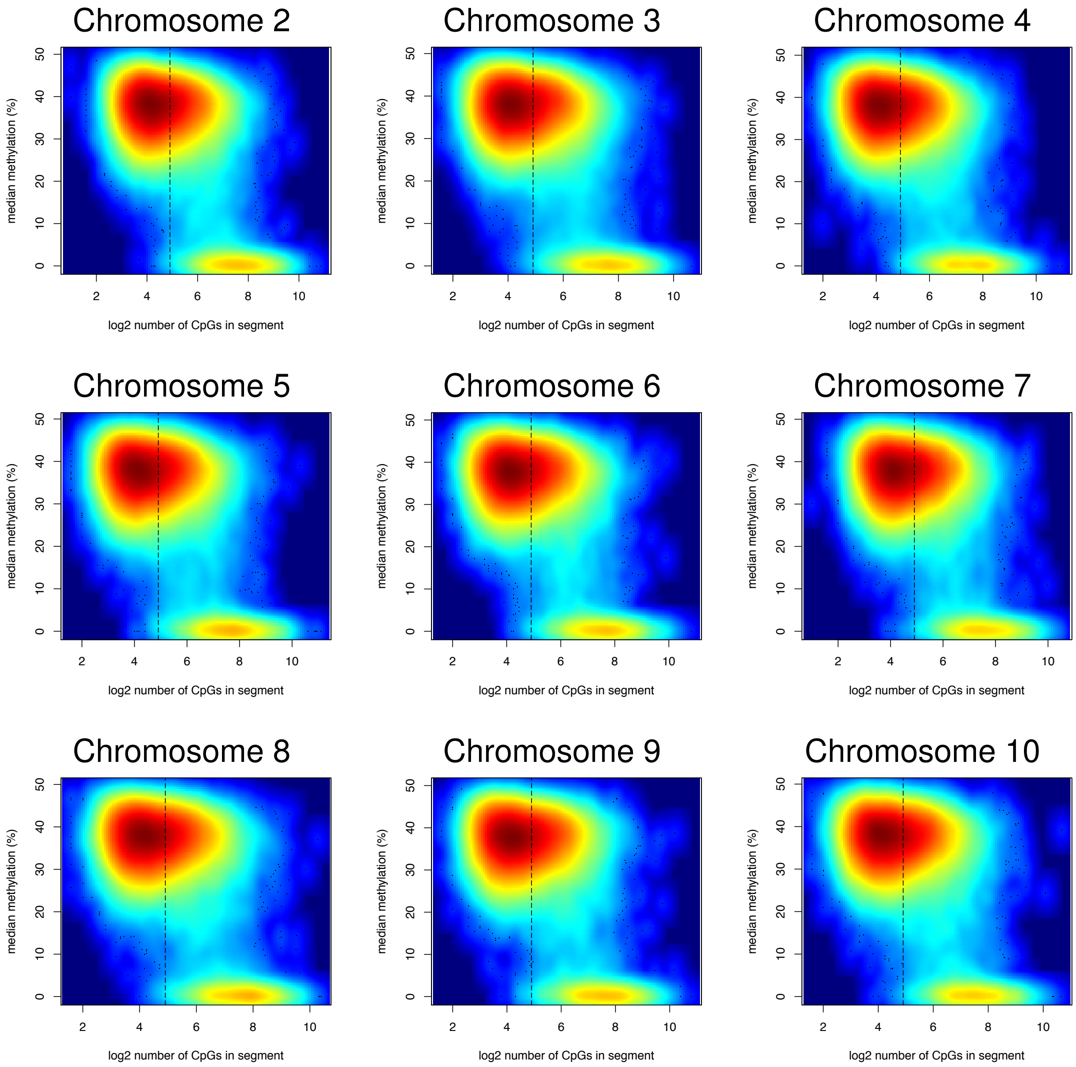
Chromosome 2
Chromosome 3
Chromosome 4
Chromosome 5
Chromosome 6
Chromosome 7
Chromosome 8
Chromosome 9
Chromosome 10
median methylation (%)
log2 number of CpGs in segment
0
10
20
30
40
50
2
4
6
8
10

(T) Maize CWG

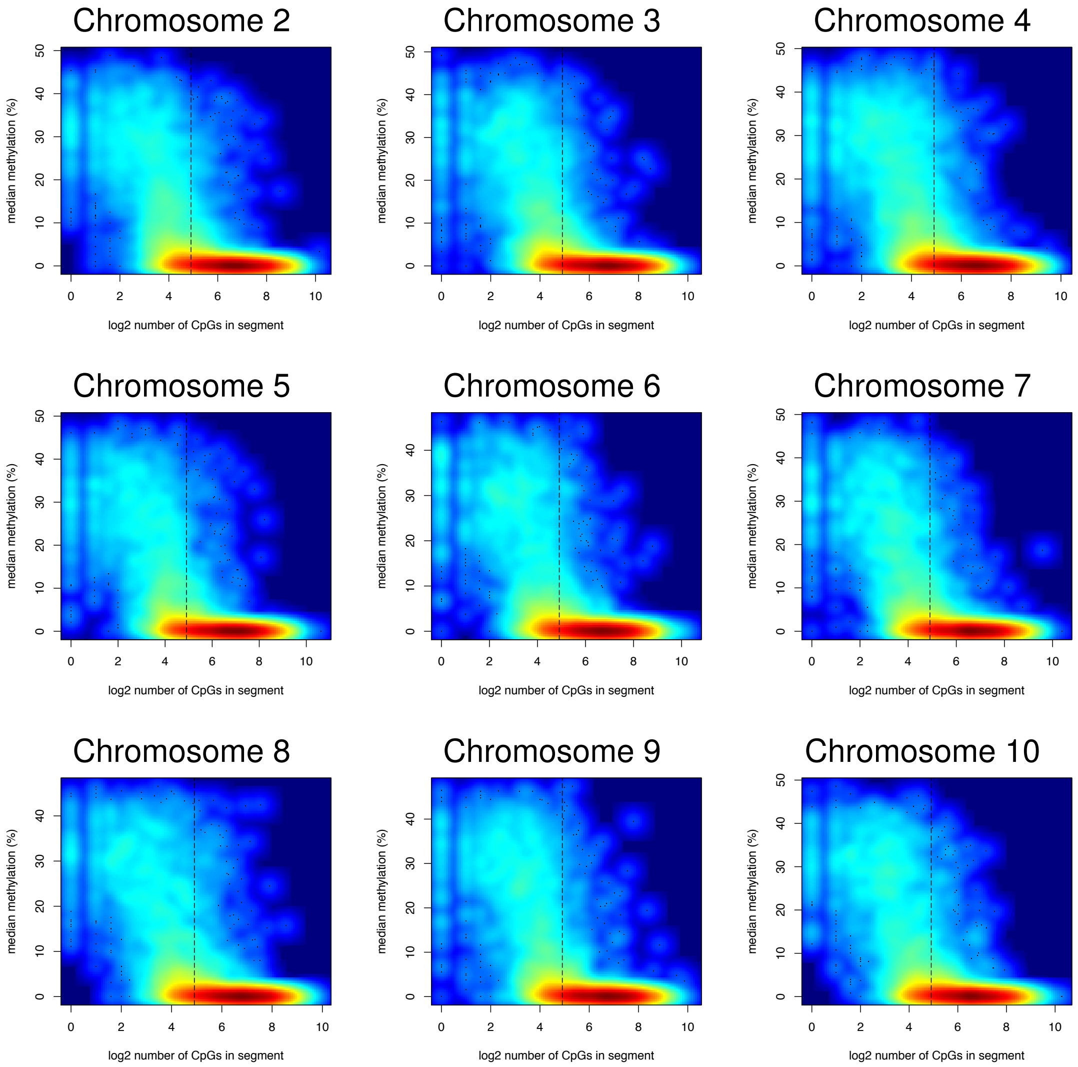
Chromosome 2
Chromosome 3
Chromosome 4
Chromosome 5
Chromosome 6
Chromosome 7
Chromosome 8
Chromosome 9
Chromosome 10
median methylation (%)
log2 number of CpGs in segment

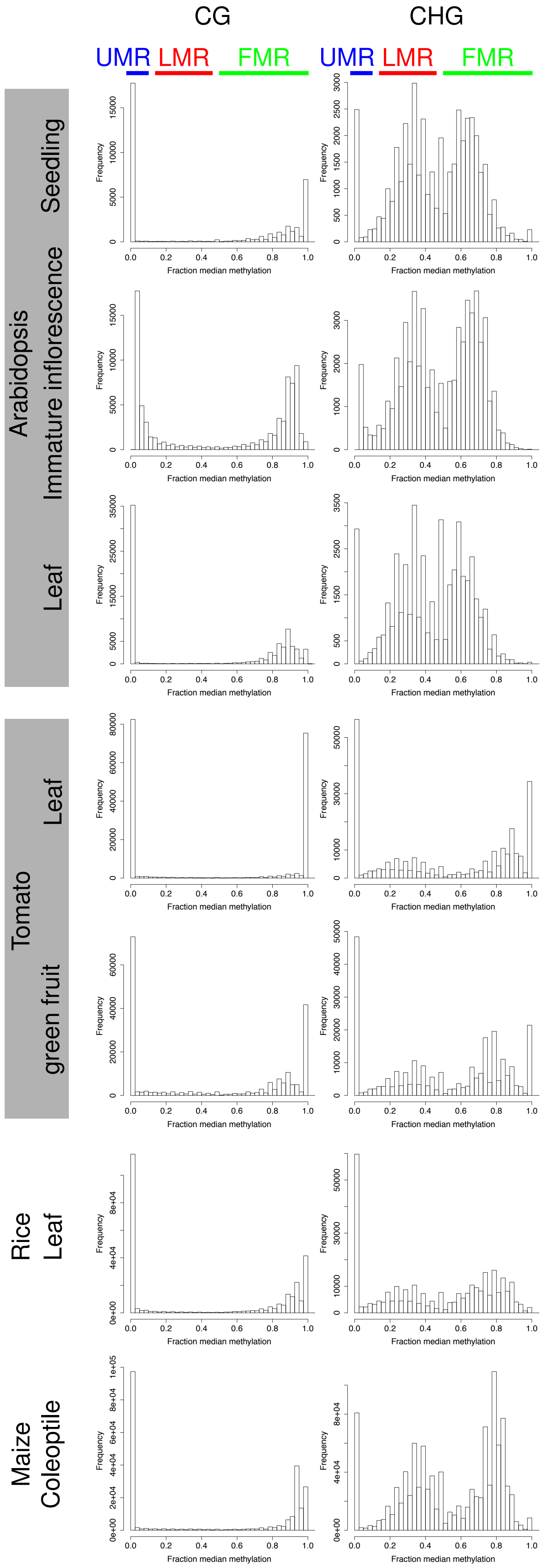


Supplementary Figure S4. Distributions of median percent methylation of U/L/FMRs for CG and CHG context. The bin size is 2.5 %, the frequency shows the number of regions with the median DNA methylation levels indicated.

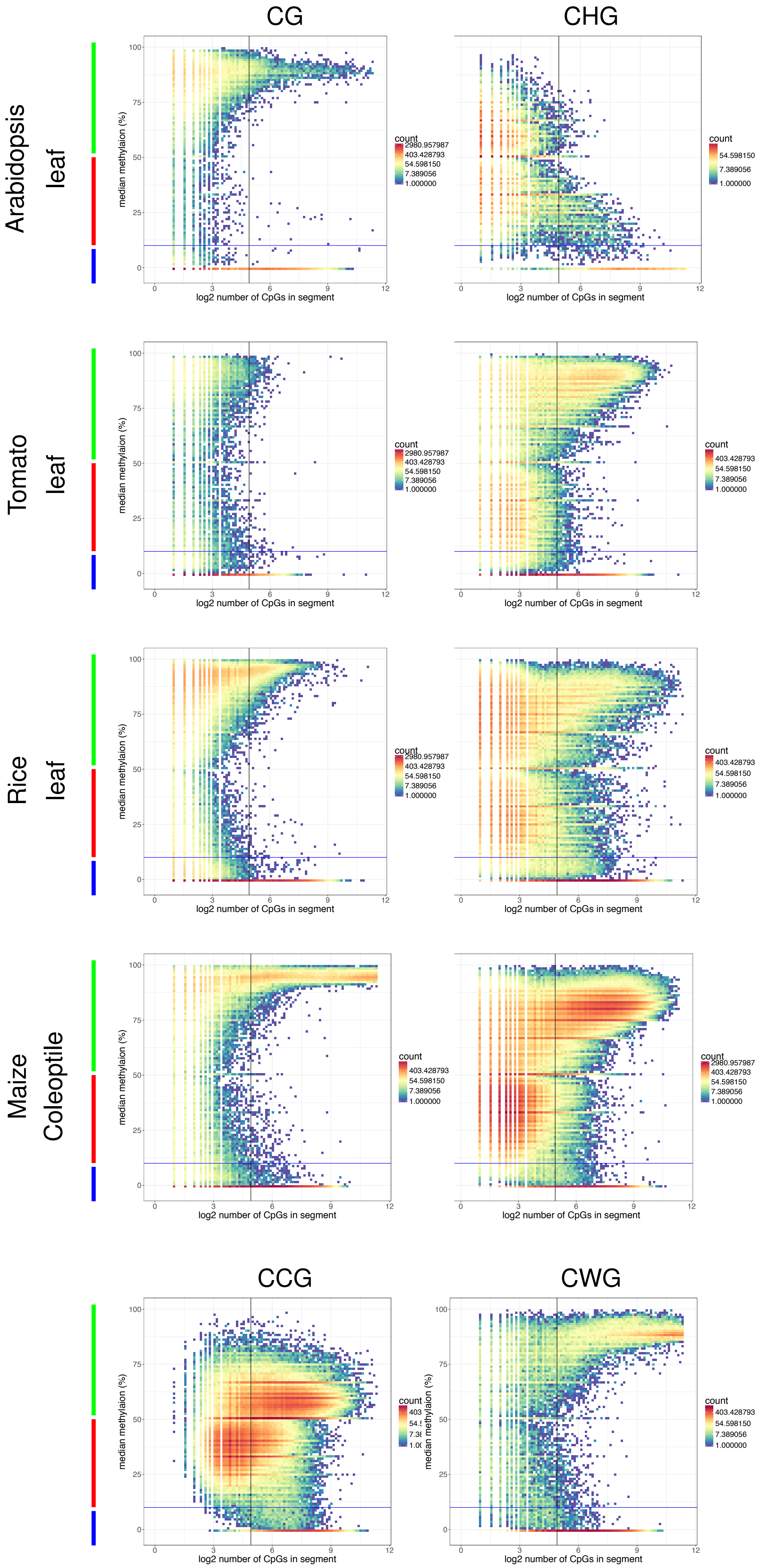

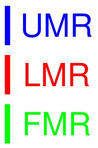


Supplementary Figure S5. Number of CpGs against median methylation of UMRs, LMRs and FMRs in CG and CHG contexts for Arabidopsis leaf, tomato leaf, rice leaf and maize coleoptile. For maize coleoptile, CCG and CWG data is show in addition. The colour indicates the number of data points at positions indicated in log scale.

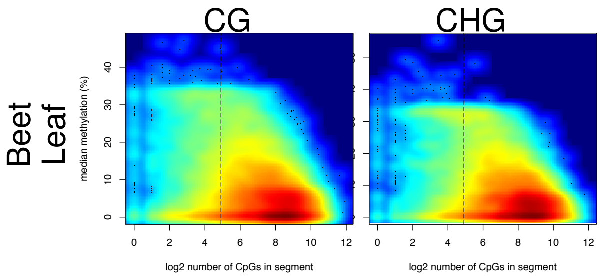


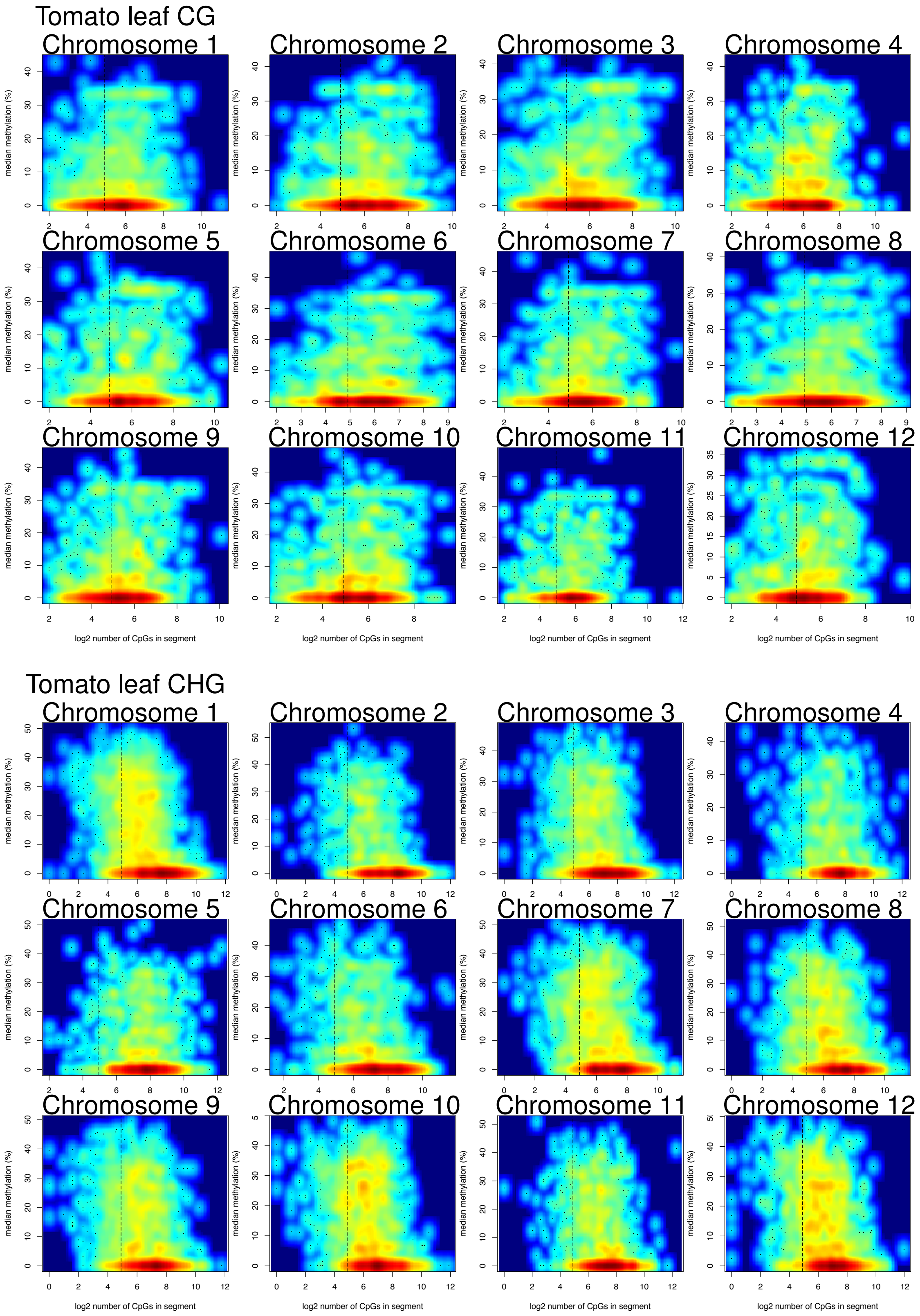


Supplementary Figure S6. Number of CpGs against median methylation of UMRs and LMRs in CG and CHG contexts for low coverage data in beet and tomato leaf.
