## additional figures 7 to 18 for "In plants distal regulatory sequences overlap with unmethylated rather than low-methylated regions, in contrast to mammals"

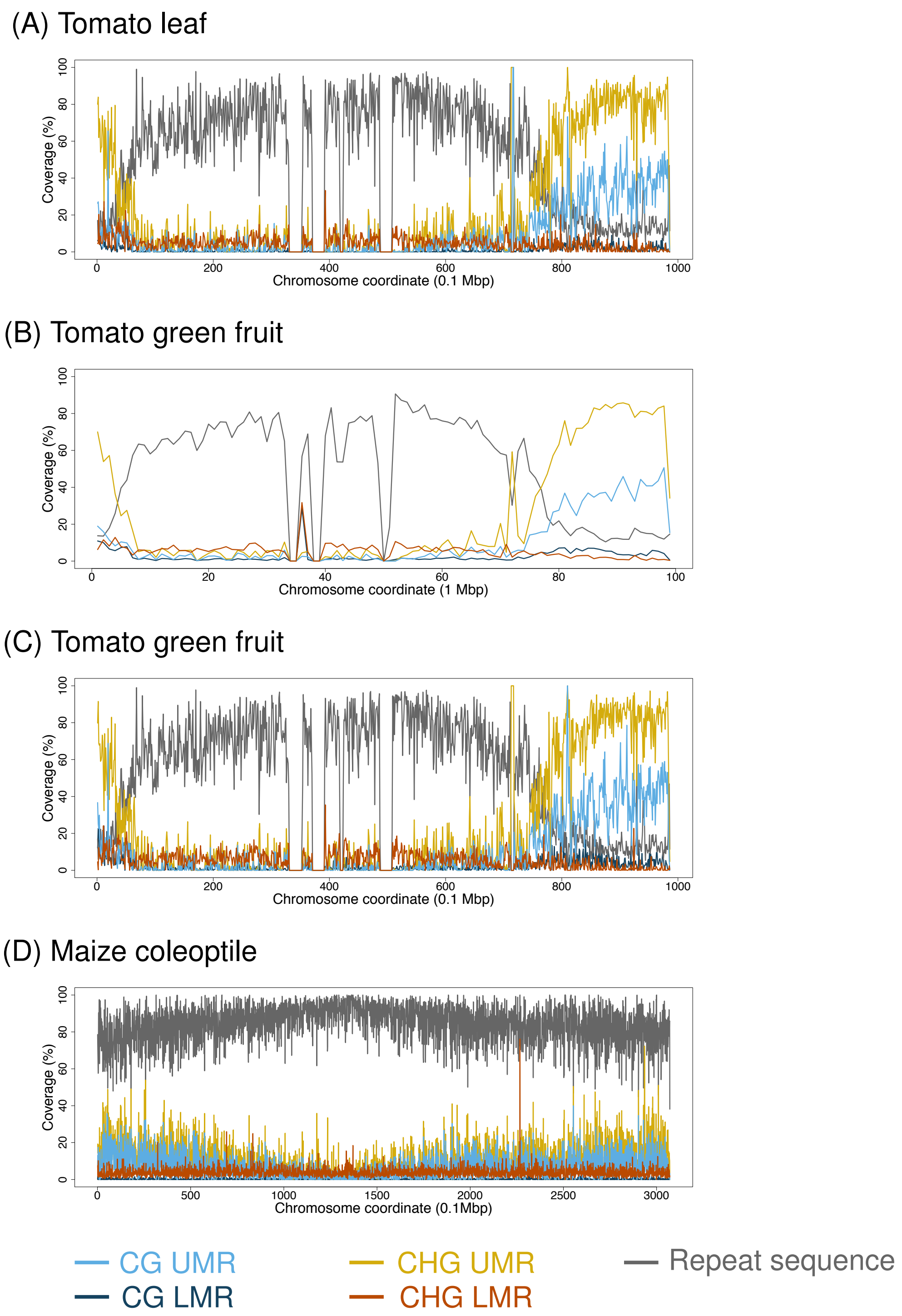


Supplementary Figure S7. Coverage of CG/CHG UMRs and LMRs over windows in chromosome 1 for (A) Tomato leaf (0.1 Mb windows), (B) Tomato green fruit (1 Mb windows), (C) Tomato green fruit (0.1 Mb windows), and (D) Maize coleoptile (0.1 Mb windows). In the tomato CHG LMR data set, a peak around the centromere has been removed as the sequence at the peak was unknown (N's).

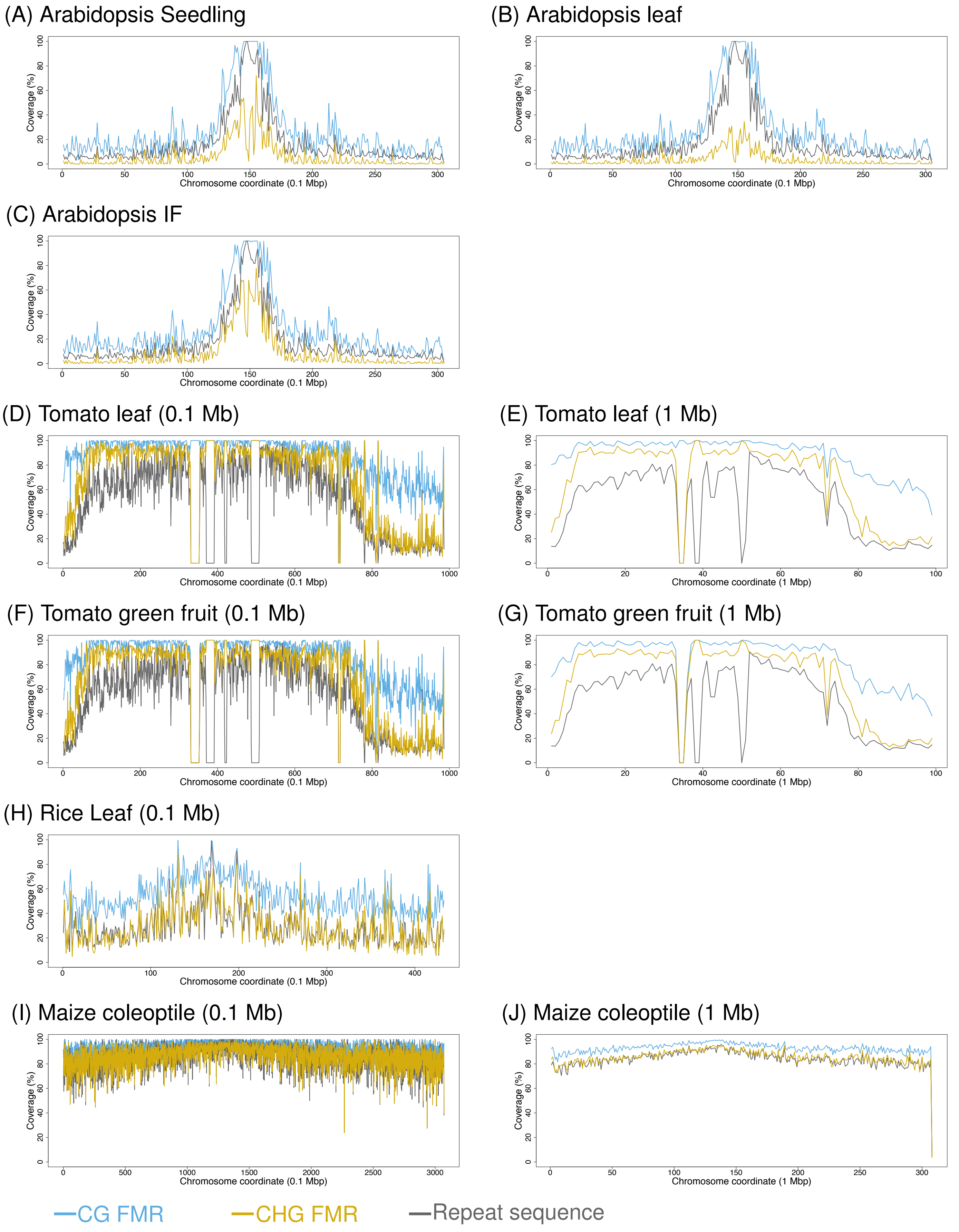


Supplementary Figure S8. Coverage of FMRs in CG and CHG context over 0.1 and 1 Mb windows in chromosome 1 for: (A-C) Arabidopsis (0.1 Mb windows), (A) seedling, (B) leaf, (C) immature florescence (IF); (D-E) Tomato leaf, (F-G) Tomato green fruit; (H) rice leaf; (I-J) Maize coleoptile.

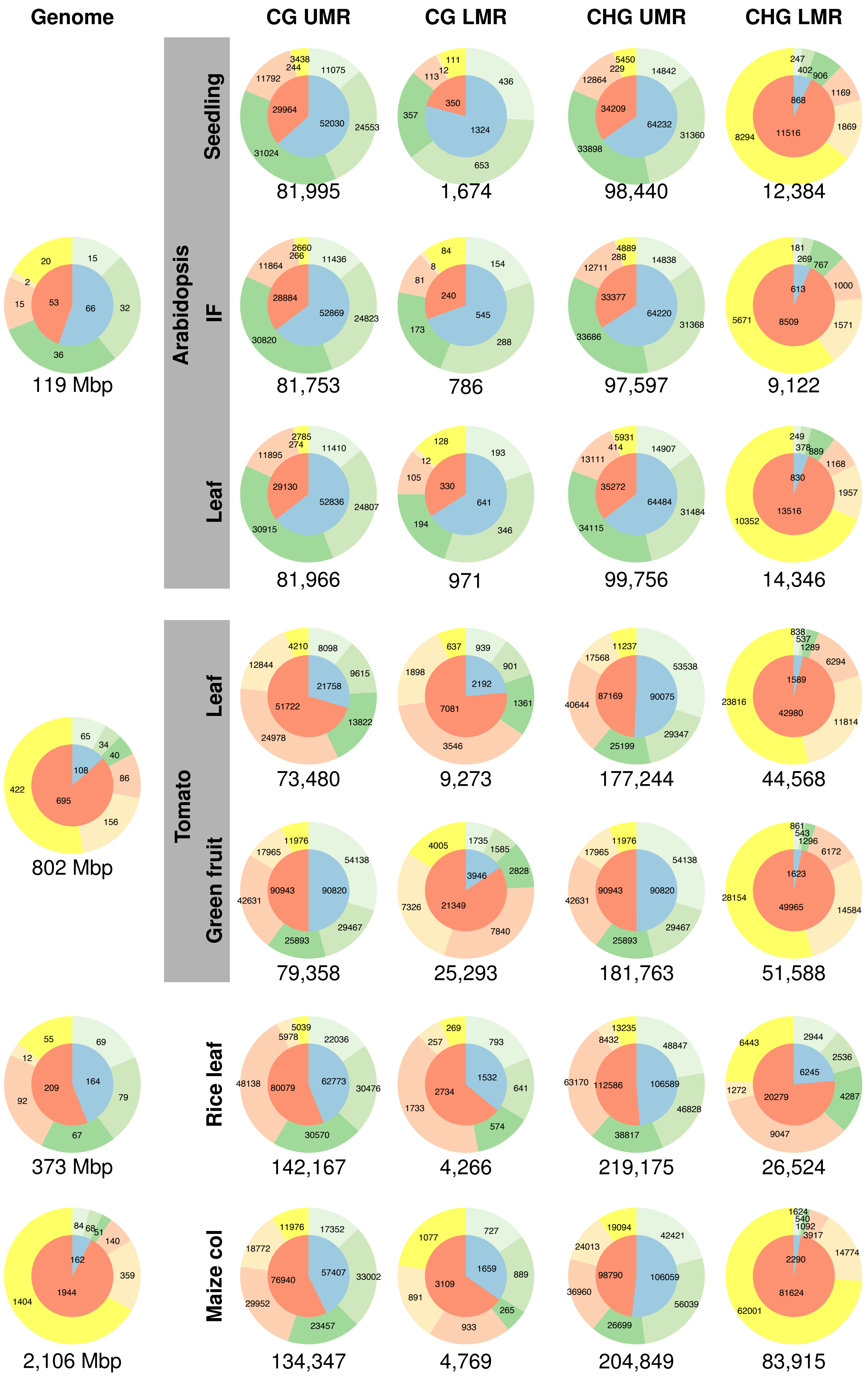

Genome
CG UMR
CG LMR
CHG UMR
CHG LMR
Arabidopsis
Seedling
IF
Leaf
Tomato
Leaf
Green fruit
Rice leaf
Maize col
20
15
2
15
53
66
32
36
119 Mbp
3438
244
11792
11075
29964
52030
24553
31024
81,995
111
12
113
436
350
357
1324
653
1,674
5450
229
12864
14842
34209
64232
31360
33898
98,440
247
402
906
1169
868
1869
8294
11516
12,384
2660
266
11864
11436
28884
52869
24823
30820
81,753
84
8
81
154
240
545
173
288
786
4889
288
12711
14838
33377
64220
31368
33686
97,597
181
269
767
1000
613
5671
8509
1571
9,122
2785
274
11895
11410
29130
52836
24807
30915
81,966
128
12
105
193
330
641
194
346
971
5931
414
13111
14907
35272
64484
31484
34115
99,756
249
378
889
1168
830
1957
10352
13516
14,346
65
34
40
108
86
422
695
156
802 Mbp
4210
8098
12844
9615
21758
51722
13822
24978
73,480
637
939
901
1898
2192
7081
1361
3546
9,273
11237
17568
53538
40644
87169
90075
29347
25199
177,244
838
537
1289
6294
1589
23816
42980
11814
44,568
11976
17965
54138
42631
90943
90820
29467
25893
79,358
1735
4005
1585
2828
3946
7326
21349
7840
25,293
11976
17965
54138
42631
90943
90820
29467
25893
181,763
861
543
1296
6172
1623
28154
49965
14584
51,588
55
69
12
209
164
79
92
67
373 Mbp
5039
5978
22036
48138
80079
62773
30476
30570
142,167
269
257
793
1532
2734
641
1733
574
4,266
13235
8432
48847
63170
112586
106589
46828
38817
219,175
6443
2944
2536
6245
1272
20279
4287
9047
26,524
84
68
51
140
162
359
1944
1404
2,106 Mbp
11976
17352
18772
57407
33002
76940
29952
23457
134,347
1077
727
1659
889
3109
891
265
933
4,769
19094
42421
24013
98790
106059
36960
56039
26699
204,849
1624
540
1092
3917
14774
2290
81624
62001
83,915

Supplementary Figure S9. Genomic composition (left) and distributions of UMRs and LMRs over the defined genomic regions of Arabidopsis (TAIR10), tomato (SL2.50), rice (RAGP7) and maize (AGPv4). IF stands for immature inflorescence and col for coleoptile. The colour code follows that in Figure 3a. The numbers in and below the pie charts indicate the occupation of each genomic region in kb for the genomic composition, and in kb for the distributions.

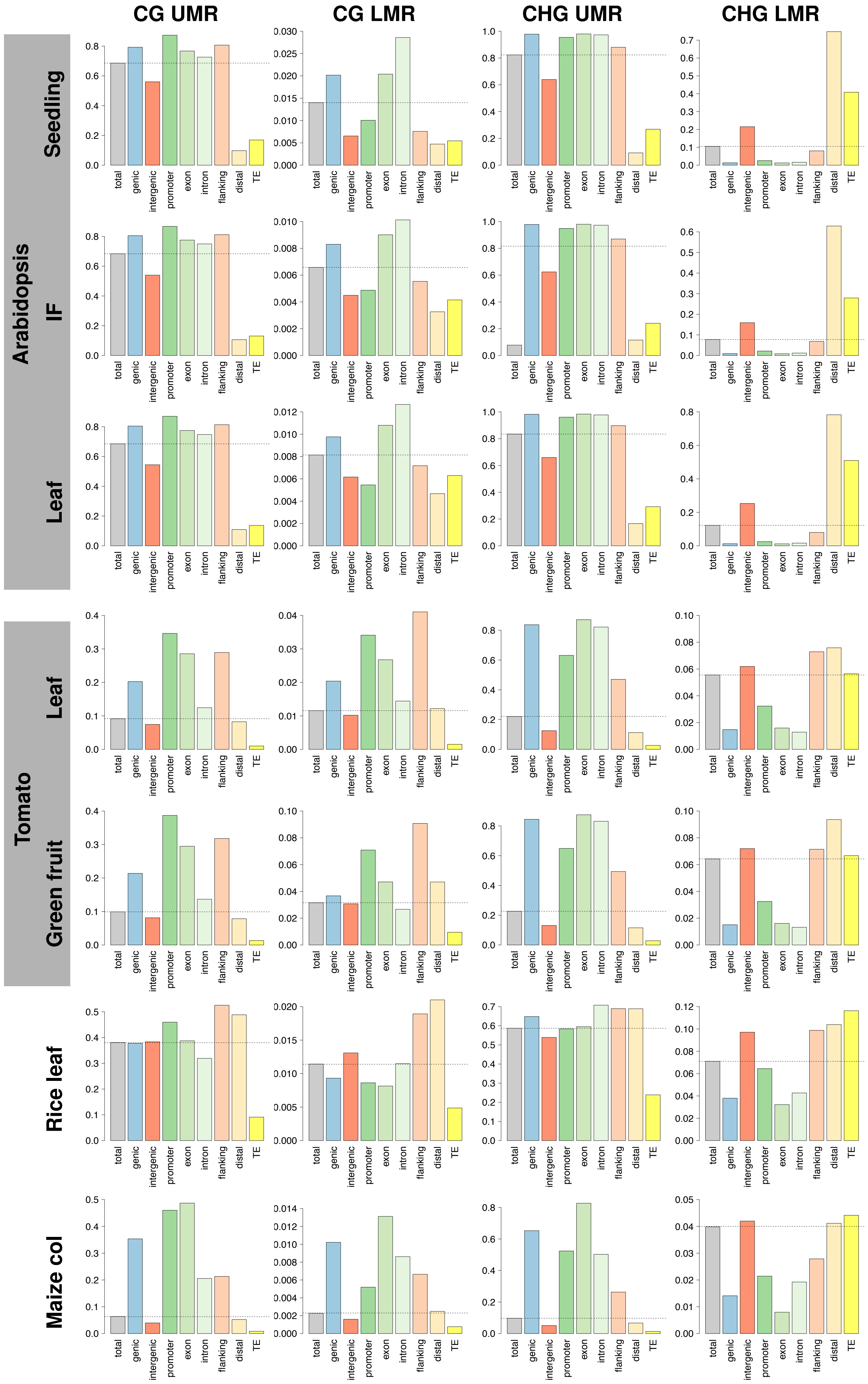

CG UMR
CG LMR
CHG UMR
CHG LMR
Arabidopsis
Seedling
IF
Leaf
Tomato
Leaf
Green fruit
Rice leaf
Maize col
total
genic
intergenic
promoter
exon
intron
flanking
distal
TE

Supplementary Figure S10. The fractions (y-axes: kbp/kbp, from 0 to 1) the CG and CHG un- and low methylated regions occupy at the various genomic regions. The gray bars indicate the fraction of overall occupancy in the genome. The same colour coding is used as defined in Figure 3a. IF: Immature inflorescence.

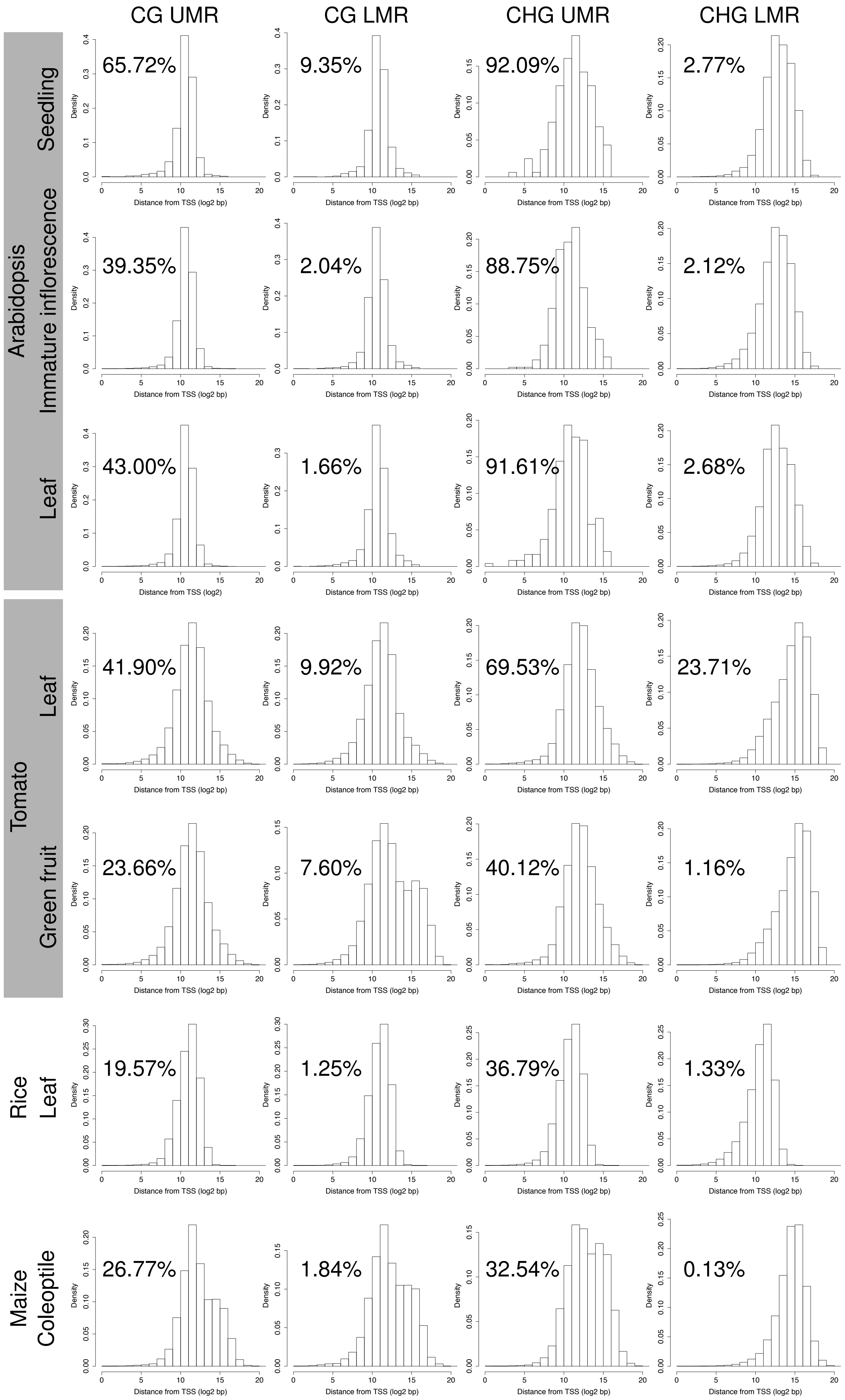
CG UMR
CG LMR
CHG UMR
CHG LMR
Arabidopsis
Seedling
65.72%
9.35%
92.09%
2.77%
Immature inflorescence
39.35%
2.04%
88.75%
2.12%
Leaf
43.00%
1.66%
91.61%
2.68%
Tomato
Leaf
41.90%
9.92%
69.53%
23.71%
Green fruit
23.66%
7.60%
40.12%
1.16%
Rice
Leaf
19.57%
1.25%
36.79%
1.33%
Maize
Coleoptile
26.77%
1.84%
32.54%
0.13%
Density
Distance from TSS (log2 bp)
Distance from TSS (log2)

Supplementary Figure S11. Distribution of distances between UMRs or LMRs and their closest TSSs in log2 scale. The UMRs or LMRs with 0 distance from a TSS (i.e. overlapping with a TSS) were excluded from the figures, but their percentages are given instead.

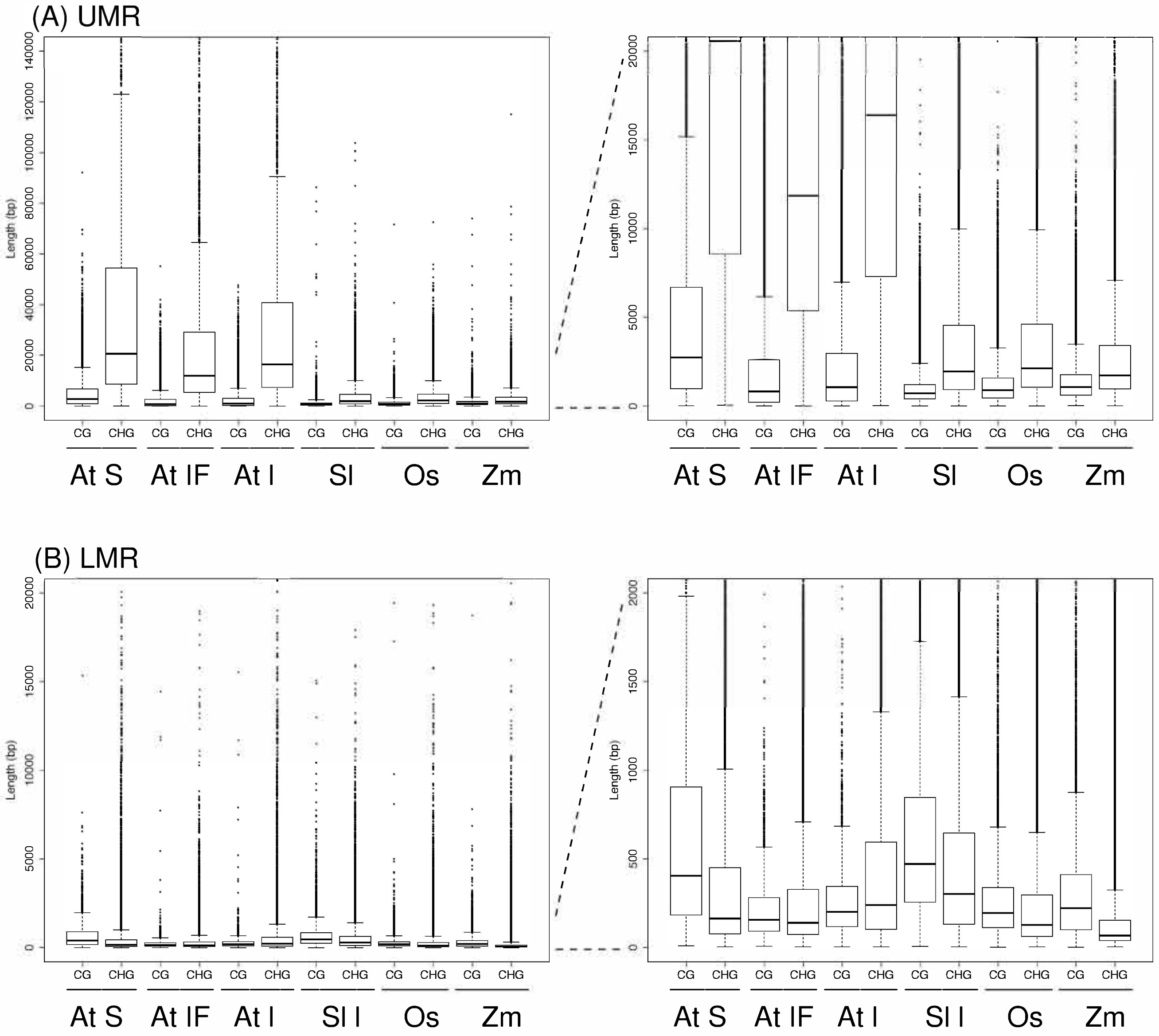


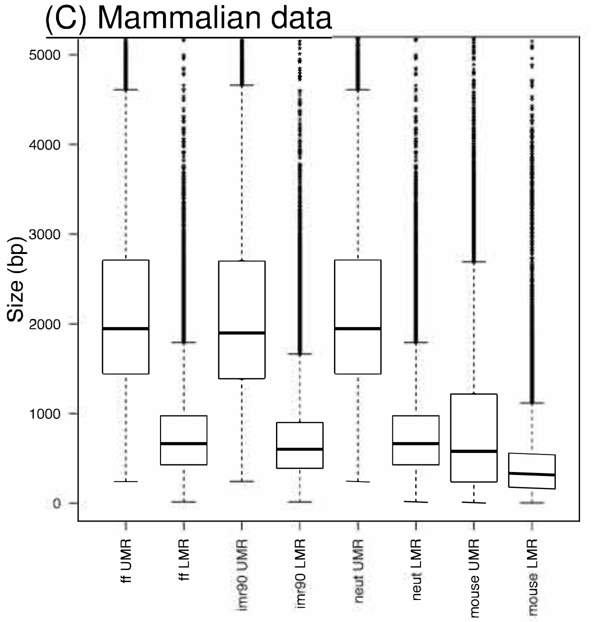


| | ff | imp90 | neut | mouse |
|---|---|---|---|---|
| **UMR** | 1947 | 1901 | 1947 | 580.0 |
| **LMR** | 665.0 | 603.0 | 665.0 | 323 |

Supplementary Figure S12. Sizes of (A) UMRs, and (B) LMRs in plants, and (C) mammals in base pairs. The median lengths (in bp) for mammalian data were given in the table. At: *Arabidopsis thaliana*, Sl: *Solanum lycopersicum*, Os: *Oryza sativa*, Zm: *Zea mays*, S: seedling, IF: Immature inflorescence and l: leaf, ff: foreskin fibroblast, imr90: fetal lung fibroblasts, neut: granulocytic neutrophils, mouse: mouse ES cells.

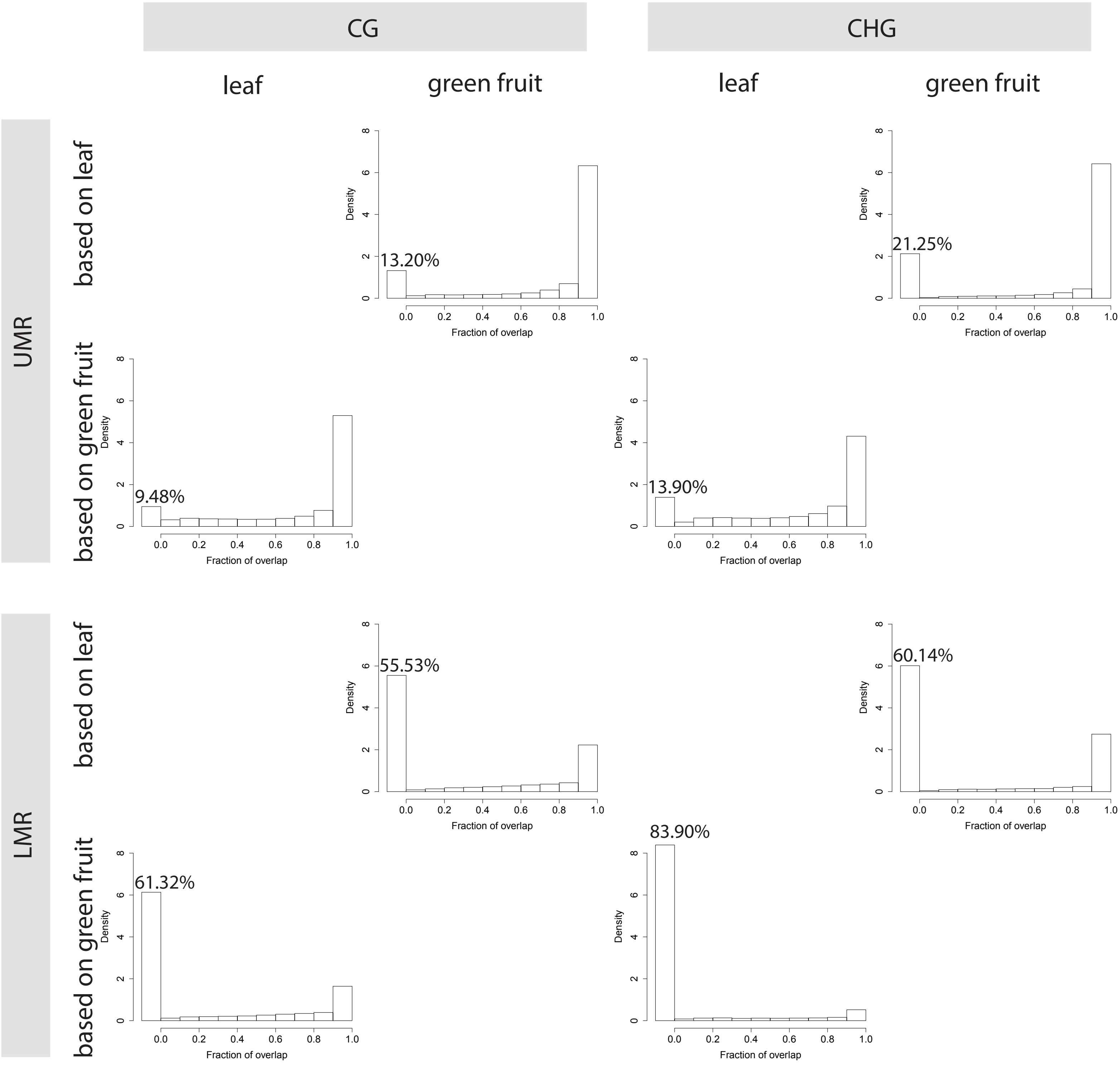


Supplementary Figure S13. Fraction of overlapping CG and CHG UMRs and LMRs between tomato leaf and green fruit. The density on the y-axis shows the fraction of UMRs or LMRs in each bin, whereby the areas of all the bars add up to 1. The percentage mentioned in each graph indicates the percentages of UMRs or LMRs without an overlap.

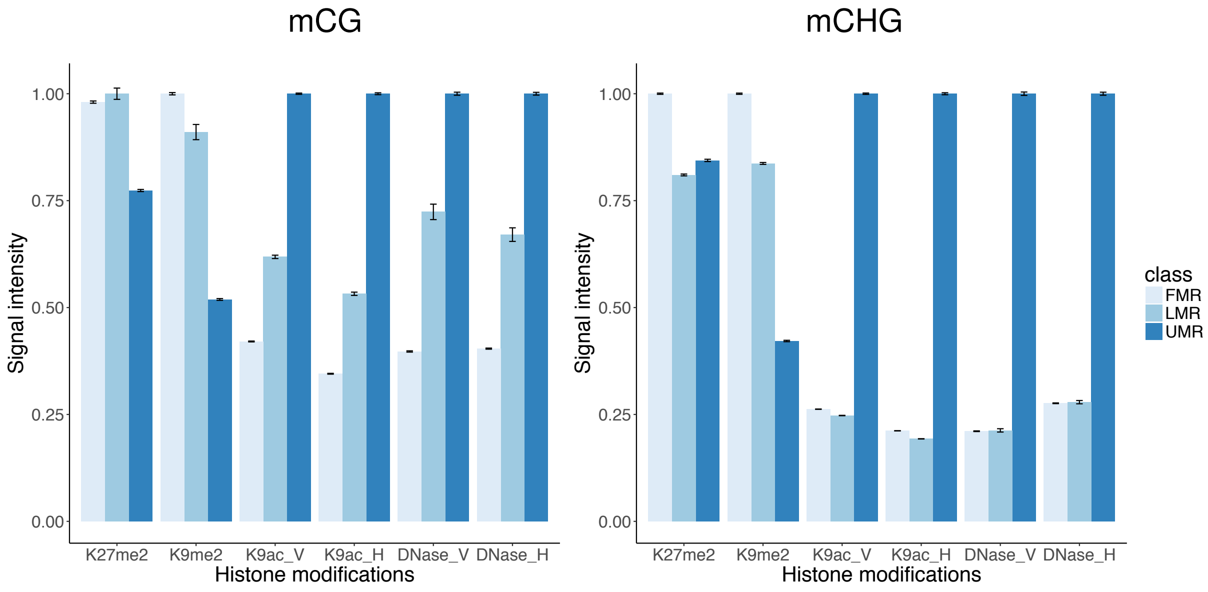


Supplementary Figure S14. Enrichment of chromatin marks in FMRs, LMRs and UMRs in the maize genome. For each mark indicated, the average signal intensities of median signal intensities in each region were calculated, and normalised to the highest bar for each mark. H3K9me2 and H3K27me2 data is derived from inner stem tissue of 1-month-old plants. The V and H at the end of K9ac and DNase indicate the data is derived from V2-IST or husk tissue, respectively.

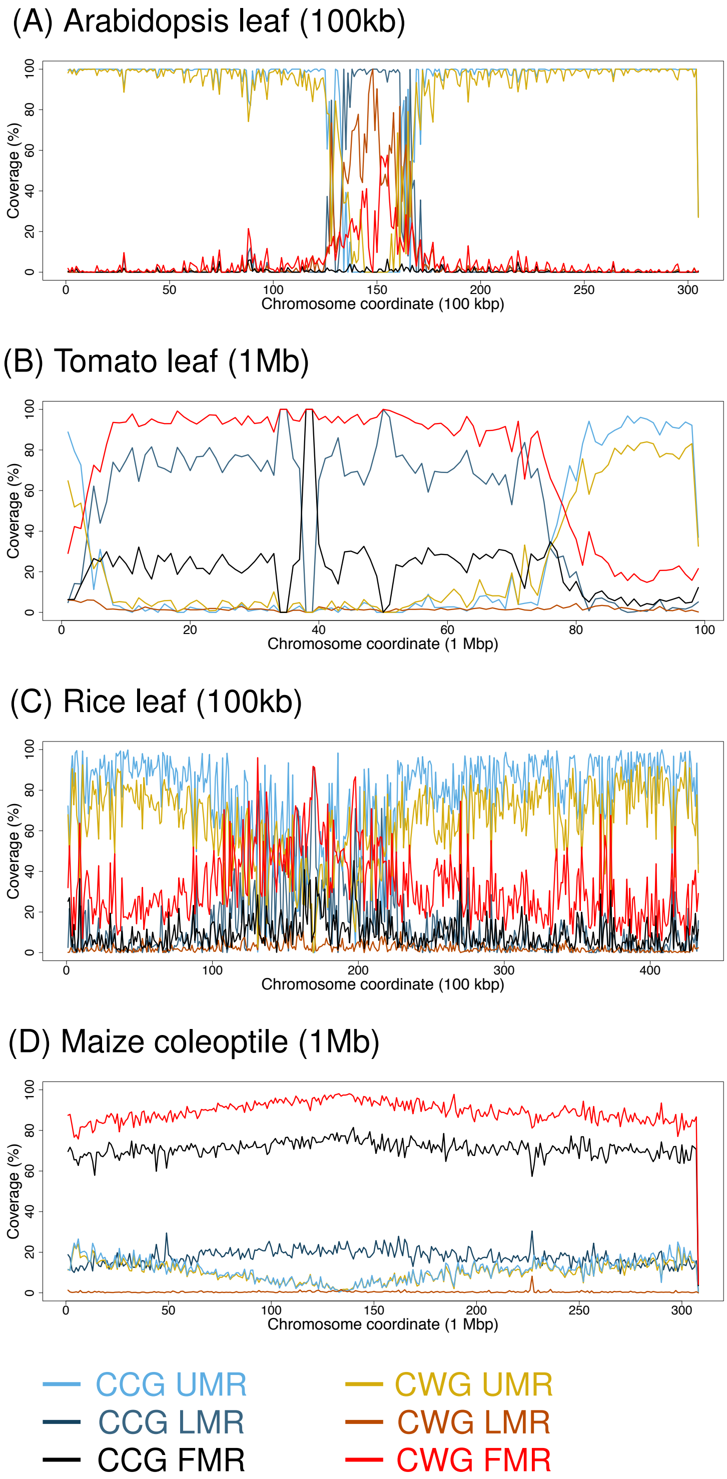


Supplementary Figure S15. Coverage of UMRs, LMRs or FMRs in CCG and CWG context over windows in chromosome 1 for (A) Arabidopsis leaf (0.1 Mb windows), (B) tomato leaf (0.1 Mb windows), (C) rice leaf (1 Mb windows), (D) Maize coleoptile (1 Mb windows).

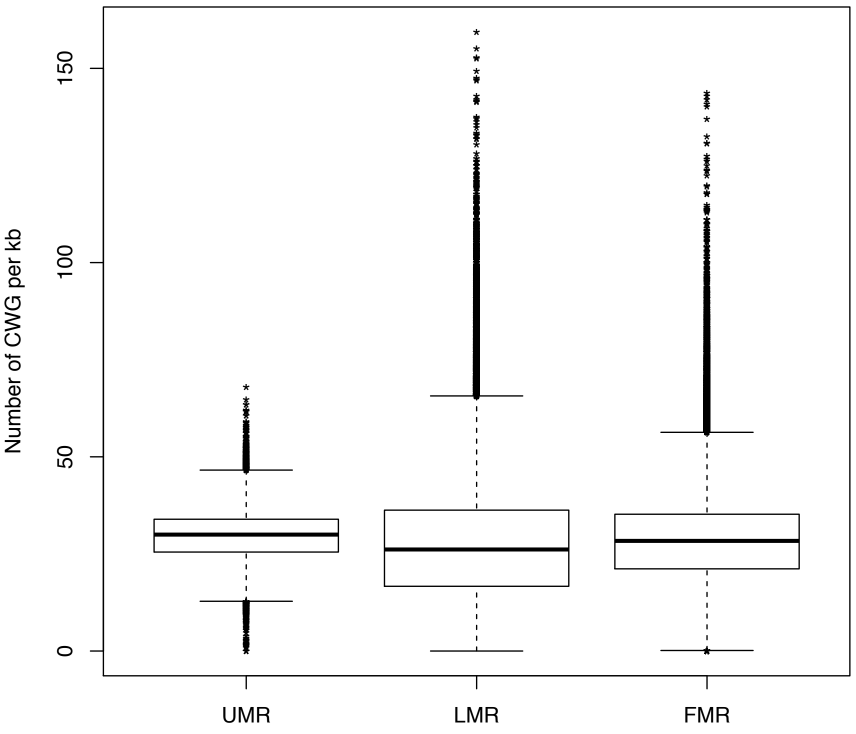


Supplementary Figure S16. Concentration of CWGs in every CCG UMR, LMR or FMR per kb in maize.

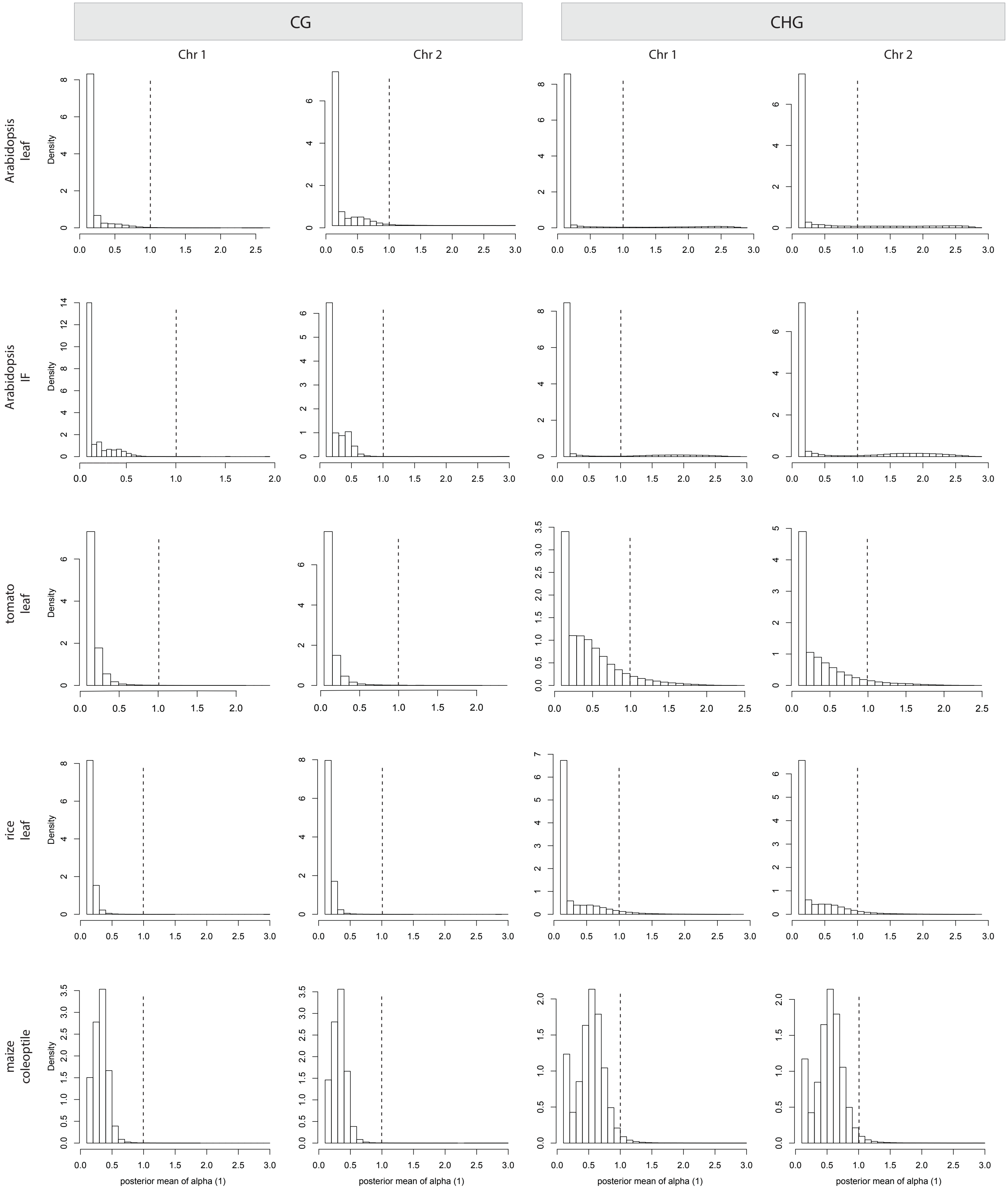

CG
CHG
Chr 1
Chr 2
Chr 1
Chr 2
Arabidopsis leaf
Arabidopsis IF
tomato leaf
rice leaf
maize coleoptile
Density
posterior mean of alpha (1)

Supplementary Figure S17. The distribution of α-values, which characterize the distribution of methylation levels in sliding windows of 100 consecutive CGs or CHGs along the genome, for chromosome 1 and 2 of the indicated species and tissues, for CG and CHG. The vertical dashed lines at $\alpha = 1.0$ indicates that the vast majority of α-values is $< 1$, indicating the absence of PMDs.

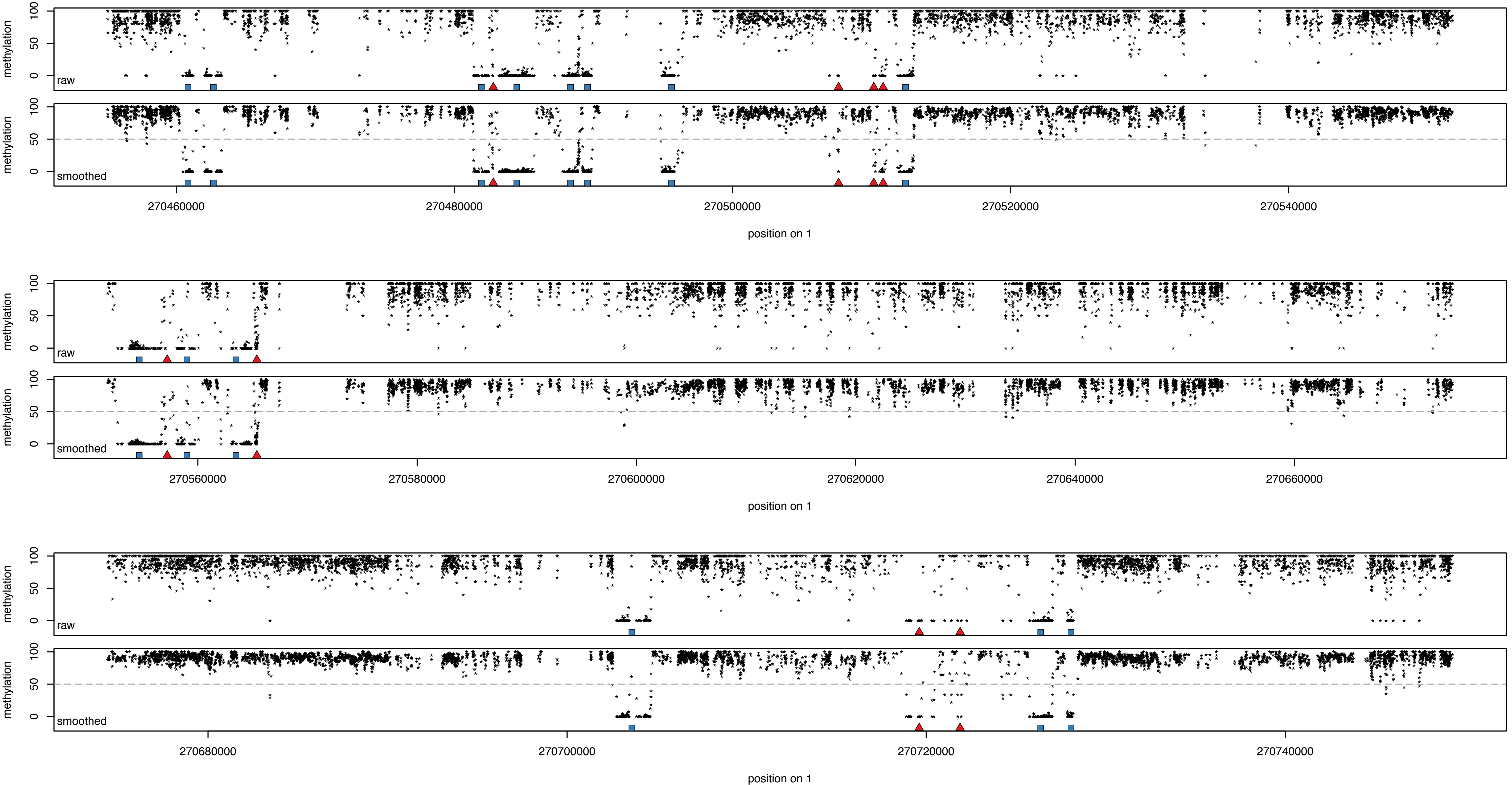

methylation
100
50
0
raw
smoothed
270460000
270480000
270500000
270520000
270540000
position on 1
270560000
270580000
270600000
270620000
270640000
270660000
270680000
270700000
270720000
270740000

Supplemental Figure S18. DNA methylation levels at randomly chosen regions of the maize genome are plotted using the plotPMDSegmentation function. Pairs of panels are shown, each pair showing the same region twice, with the raw methylation levels at the top and the methylation levels smoothened over 3 consecutive CGs or CHGs at the bottom. The horizontal grey dashed lines (bottom) indicate the 50% cut-off for LMRs. UMRs are indicated with blue squares, LMRs with red triangles.
