## additional tabels for "In plants distal regulatory sequences overlap with unmethylated rather than low-methylated regions, in contrast to mammals"

Table S1. Summary of datasets used in this study.

| Species | Tissue | Read length | unique reads | avg. cov. | Accession |
| --- | --- | --- | --- | --- | --- |
| Arabidopsis  (119 Mb) | Seedling | 150-150 PE | 7,235,544 | 18.2x | GSE99404 (Lauss et al 2018) |
|  | Leaf | 51 SE | NA | NA | GSM2078421 (Harris et al 2016) |
|  | IF | 51 SE | NA | NA | GSM1848703 (Zhai et al 2015) |
| Beet  (376 Mb) | Leaf | 100 SE  150 SE | 33,597,598  75,133 | 5.9x | GSM2096947 (Niederhuth et al 2016) |
| Tomato  (802 Mb) | Leaf | 101-101 PE | 117,411,390 | 29.6x | SRX151217 (Zhong et al, 2013) |
|  | Green fruit | 101-101 PE | 116,394,780 | 29.3x | SRX098309 (Zhong et al, 2013) |
|  | Leaf | 125-125 PE | 11,408,287 | 3.6x | SRX2008738 (Gouil et al 2016) |
| Rice  (373 Mb) | Leaf | 55 SE | 171,626,718 | 25.3x | GSM946552 (Chodavarapu et al 2012) |
| Maize  (2,106 Mb) | Coleoptile | 50-50 PE | 856,082,578 | 20.3x | GSE39232 (Regulski et al) |

Analysed data provided by the original study was used for the data indicated as NA (Not Available; no read number or average coverage were calculated for these). SE stands for single-end, PE for paired-end sequencing.

Table S2. Number of CG LMRs and UMRs in different human and mouse cell types.

| **Organism** | **Cell type** | **UMR** | **LMR** | **ratio** |
| --- | --- | --- | --- | --- |
| human | bcell | 21,078 | 71,020 | 2:7 |
|  | cd133hsc | 20,230 | 45,536 | 1:2 |
|  | neutrophil | 21,378 | 72,985 | 2:7 |
|  | h1 | 18,263 | 29,609 | 2:3 |
|  | imr90 | 16,695 | 35,030 | 1:2 |
|  | ads_adipose | 21,933 | 59,906 | 2:3 |
|  | ads_ipsc | 20,121 | 33,475 | 2:3 |
|  | ads | 21,754 | 55,031 | 2:5 |
|  | ff_ips_19_11_bmp4 | 19,430 | 80,417 | 1:4 |
|  | ff_ipsc_19_11 | 19,147 | 31,566 | 2:3 |
|  | ff_ipsc_19_7 | 19,178 | 32,079 | 2:3 |
|  | ff_ipsc_6_9 | 18,932 | 31,485 | 2:3 |
|  | ff | 17,395 | 37,205 | 2:3 |
|  | h1_bmp4 | 18,733 | 42,814 | 1:2 |
|  | h9_laurent | 19,549 | 31,633 | 2:3 |
|  | h9 | 18,182 | 36,850 | 1:2 |
|  | imr90_ipsc | 18,986 | 29,507 | 2:3 |
| mouse | ES cell | 33,675 | 26,335 | 6:5 |

The data is derived from human cell types (Burger et al 2013) and mouse ES cells (Stadler et al 2013).

Table S3. Median and maximum sizes of LMRs and UMRs in bp.

| **Median** | **UMR** | | | **LMR** | |
| --- | --- | --- | --- | --- | --- |
|  | **CG** | **CHG** | **CG** | | **CHG** |
| At Seedling | 2,741.0 | 20,570 | 405 | | 165 |
| At IF | 811.5 | 11,850 | 158 | | 141 |
| At leaf | 1,050 | 16,390 | 203 | | 241 |
| Tomato leaf | 709 | 1,942 | 471 | | 304 |
| Tomato green fruit | 860 | 2,232 | 388 | | 294 |
| Rice leaf | 876 | 2,109 | 196.0 | | 129 |
| Maize coleoptile | 1,059 | 1,713  (CCG: 4,196)  (CWG: 1,755) | 223 | | 69  (CCG: 390)  (CWG: 379) |

| **Max** | **UMR** | | | **LMR** | |
| --- | --- | --- | --- | --- | --- |
|  | **CG** | **CHG** | **CG** | | **CHG** |
| At Seedling | 92,130 | 807,000 | 27,060 | | 155,100 |
| At IF | 55,070 | 516,000 | 39,270 | | 54,610 |
| At leaf | 47,600 | 518,000 | 69,810 | | 77,800 |
| Tomato leaf | 625,600 | 625,700 | 363,300 | | 2,281,000 |
| Tomato green fruit | 280,300 | 671,500 | 2,281,000 | | 2,346,000 |
| Rice leaf | 71,590 | 72,470 | 23,370 | | 63,240 |
| Maize col | 299,600 | 299,900 | 33,090 | | 622,500 |

Table S4. Overlap between CHG UMRs/LMRs and DHSs, H3K9ac enriched regions, and enhancer candidates.

| (A) |  | **CHG UMR** | **CHG LMR** |
| --- | --- | --- | --- |
|  | **All** | 77,700 | 460,329 |
| **DHS** | **V2-IST** | 15,850 (20.4%) | 312 (0.1%) |
|  | **Husk** | 13,586 (17.5%) | 257 (0.1%) |
| **H3K9ac** | **V2-IST** | 16,268 (20.9%) | 116 (0.0%) |
|  | **Husk** | 26,520 (34.1%) | 881 (0.2%) |
| **Candidate** | **V2-IST** | 396 (0.5%) | 0 (0%) |
|  | **Husk** | 1291 (1.7%) | 0 (0%) |

| (B) | |  | **CHG UMR** | | **CHG LMR** | |
| --- | --- | --- | --- | --- | --- | --- |
|  | | **all** | **> 1 bp overlap** | **> 50% overlap** | **> 1 bp overlap** | **> 50% overlap** |
| **DHS** | **V2** | 18485 | 18,270 (98.8%) | 18,072 (97.8%) | 281 (1.5%) | 163 (0.9%) |
|  | **Husk** | 15261 | 15,063 (98.7%) | 14,902 (97.6%) | 238 (1.6%) | 152 (1.0%) |
| **H3K9ac** | **V2** | 17170 | 17,135 (99.8%) | 17,094 (99.6%) | 93 (0.5%) | 22 (0.1%) |
|  | **Husk** | 28948 | 27,927 (96.5%) | 27,560 (95.2%) | 63 (0.2%) | 63 (0.2%) |
| **Candidate** | **V2** | 398 | 398 (100%) | 398 (100%) | 0 (0%) | 0 (0%) |
|  | **Husk** | 1320 | 1320 (100%) | 1320 (100%) | 0 (0%) | 0 (0%) |

(A) Number of CHG UMRs and LMRs overlapping with DHS, H3K9ac enriched regions, or enhancer candidates. (B) Number of DHS, H3K9ac enriched regions or enhancer candidates overlapping > 1 bp or >50% with CHG UMRs and LMRs. V2-IST refers to inner stem tissue of maize seedlings in a V2 stage, husk to husk leaves (Oka et al 2017). Candidate refers to the enhancer candidates identified in Oka et al (2017).

Table S5. Overlap between CG UMRs/LMRs and DHSs, H3K9ac enriched regions, and enhancer candidates.

| (A) |  | **CG UMR** | **CG LMR** |
| --- | --- | --- | --- |
|  | **All** | 95,869 | 14,493 |
| **DHS** | **V2-IST** | 17,067 (17.8%) | 366 (2.5%) |
|  | **Husk** | 14,225 (14.8%) | 244 (1.7%) |
| **H3K9ac** | **V2-IST** | 17,120 (17.9%) | 257 (1.8%) |
|  | **Husk** | 29,710 (31.0%) | 991 (6.8%) |
| **Candidate** | **V2-IST** | 395 (0.4%) | 0 (0%) |
|  | **Husk** | 1304 (1.4%) | 0 (0%) |

| (B) | |  | **CG UMR** | | **CG LMR** | |
| --- | --- | --- | --- | --- | --- | --- |
|  | | **all** | **> 1 bp overlap** | **> 50% overlap** | **> 1 bp overlap** | **> 50% overlap** |
| **DHS** | **V2** | 18485 | 18,210 (98.5%) | 17,866 (96.7%) | 333 (1.8%) | 167 (0.9%) |
|  | **Husk** | 15261 | 15,031 (98.5%) | 14,815 (97.1%) | 228 (1.5%) | 149 (1.0%) |
| **H3K9ac** | **V2** | 17170 | 17,089 (99.5%) | 16,998 (98.9%) | 220 (1.3%) | 32 (0.2%) |
|  | **Husk** | 28948 | 27,845 (96.2%) | 26,961 (93.1%) | 862 (3.0%) | 58 (0.2%) |
| **Candidate** | **V2** | 398 | 398 (100%) | 398 (100%) | 0 (0%) | 0 (0%) |
|  | **Husk** | 1320 | 1320 (100%) | 1320 (100%) | 0 (0%) | 0 (0%) |

(A) Number of CG UMRs and LMRs overlapping with DHS, H3K9ac enriched regions, or enhancer candidates. (B) Number of DHS, H3K9ac enriched regions or enhancer candidates overlapping > 1 bp or >50% with CG UMRs and LMRs. V2-IST refers to inner stem tissue of maize seedlings in a V2 stage, husk to husk leaves (Oka et al 2017). Candidate refers to the enhancer candidates identified in Oka et al (2017).

Table S6. Number of CHG/CG UMRs/LMRs that overlap with ACRs and vice versa.

| (A) |  | **CHG UMR** | **CHG LMR** |
| --- | --- | --- | --- |
|  | **All** | 77,700 | 460,329 |
| **dACR** | **Leaf** | 7,827 (10.1%) | 448 (0.1%) |
|  | **Inflorescence** | 7,408 (9.5%) | 164 (0.0%) |
| **aACR** | **Leaf** | 21,785 (28.0%) | 767 (0.2%) |
|  | **Inflorescence** | 19,606 (25.2%) | 334 (0.1%) |

| (B) |  | **CG UMR** | **CG LMR** |
| --- | --- | --- | --- |
|  | **All** | 95,869 | 14,493 |
| **dACR** | **Leaf** | 7,933 (8.3%) | 144 (1.0%) |
|  | **Inflorescence** | 7,482 (7.8%) | 94 (0.6 %) |
| **aACR** | **Leaf** | 23,669 (24.7%) | 298 (2.1%) |
|  | **Inflorescence** | 20,943 (21.8%) | 174 (1.2%) |

| (C) | |  | **CHG UMR** | | **CHG LMR** | |
| --- | --- | --- | --- | --- | --- | --- |
|  | | **all** | **> 1 bp overlap** | **> 50% overlap** | **> 1 bp overlap** | **> 50% overlap** |
| **dACRs** | **L** | 10651 | 9,877 (92.3%) | 9,702 (91.1%) | 377 (3.5%) | 131 (1.2%) |
|  | **I** | 9330 | 9,035 (96.8%) | 8,938 (95.8%) | 155 (1.7%) | 87 (0.9%) |
| **aACR** | **L** | 32481 | 31,208 (96.1%) | 30,679 (94.5%) | 679 (2.1%) | 188 (0.6%) |
|  | **I** | 27288 | 26,717(98.0%) | 26,430 (96.9%) | 312 (1.1%) | 121 (0.4%) |

| (D) | |  | **CG UMR** | | **CG LMR** | |
| --- | --- | --- | --- | --- | --- | --- |
|  | | **all** | **> 1 bp overlap** | **> 50% overlap** | **> 1 bp overlap** | **> 50% overlap** |
| **dACR** | **L** | 10651 | 9,814 (92.1%) | 9,603 (90.2%) | 132 (1.2%) | 74 (0.7%) |
|  | **I** | 9330 | 8,986 (96.3%) | 8,850 (94.9%) | 90 (1.0%) | 63 (0.7%) |
| **aACR** | **L** | 32481 | 30,915 (95.2%) | 30,219 (93.0%) | 281 (0.9%) | 138 (0.4%) |
|  | **I** | 27288 | 26,503 (97.1%) | 26,088 (95.6%) | 169 (0.6%) | 100 (0.4%) |

(A-B) Number of (A) CHG and (B) CG UMRs and LMRs that overlap with accessible chromatin regions (ACRs), for all ACRs (aACRs) and for distal ACRs (dACRs) only. (C-D) Number of dACRs and aACRs overlapping > 1 bp or >50% with (C) CHG and (D) CG UMRs and LMRs. L refers to inner leaves of a VE stage seedling, I to inflorescence primordia of the ear (Ricci et al. 2019). The ACRs examined were identified by Ricci *et al.* (2019).

Table S7. Overlap between CCG and CWG UMRs, LMRs and FMRs.

| (A) |  | **CCG UMR** | **CCG LMR** | **CCG FMR** |
| --- | --- | --- | --- | --- |
|  | **Total bp in regions indicated** | 220,612,889 | 398,334,999 | 1496338533 |
| **CWG UMR** | 202,906,785 | 177,886,621 | 22,566,188 | 2,453,856 |
| **CWG LMR** | 10,284,930 | 1,397,362 | 6,804,508 | 2,083,041 |
| **CWG FMR** | 1,893,094,716 | 4,132,878 | 359,697,713 | 1,491,801,491 |

| (B) | **% of bp in CWG region indicated** | **CCG UMR** | **CCG LMR** | **CCG FMR** |
| --- | --- | --- | --- | --- |
| **CWG UMR** | 9.64 | 80.63; 8.37 | 5.80; 0.60 | 0.16; 0.02 |
| **CWG LMR** | 0.49 | 0.63; 1.30 | 1.75; 3.58 | 0.14; 0.28 |
| **CWG FMR** | 89.87 | 18.78; 0.21 | 92.46; 1.03 | 99.70; 1.11 |

(A) Number of base pairs (bp) covered by CWG and CCG regions, and the overlap between these regions in maize. The heading ‘total bp in regions indicated’ holds for the second column and second row. (B) Percentages of base pairs in CCG regions that overlap with base pairs in the CWG regions indicated; the ratio between observed and expected percentages.
